# A computational model for individual differences in non-reinforced learning for individual items

**DOI:** 10.1101/2022.03.20.484477

**Authors:** Tom Salomon, Alon Itzkovitch, Nathaniel D. Daw, Tom Schonberg

**Author notes:** Correspondence concerning this article should be addressed to Tom Schonberg, Department of Neurobiology, Faculty of Life Sciences and Sagol School of Neuroscience, Tel Aviv University, Ramat Aviv 6997801, Tel Aviv, Israel.

## Abstract

Cue-Approach Training (CAT) is a paradigm that enhances preferences without external reinforcmeents, suggesting a potential role for internal learning processes. Here, we developed a novel Bayesian computational model to quantify anticipatory response patterns during the training phase of CAT. This phase includes individual items and thus this marker is potentially of internal learning signals at the item level. Our model, fitted to meta-analysis data from 29 prior CAT experiments, was able to predict individual differences in non-reinforced preference changes using a key computational marker. Crucially, two new experiments manipulated the training procedure to influence the model’s predicted learning marker. As predicted and preregistered, the manipulation successfully induced differential preference changes, supporting a causal role of our model. These findings demonstrate powerful potential of our computational framework for investigating intrinsic learning processes. This framework could be used to predict preference changes and opens new avenues for understanding intrinsic motivation and decision-making.

**Teaser:** Bayesian modeling of response time predicts individual differences in non reinforced preference change.

## Introduction

Value construction and modification are commonly understood as the result of learning processes that rely on external reinforcements ^1–4^. While reinforcement-based learning dominated the research field of value-modifications, a recent review^5^ highlighted the various means to influence preferences and choices without external reinforcements covered in the literature, dating back to the *mere exposure effect*. In those studies preferences for stimuli could be enhanced merely by repeated exposure to stimuli ^6^. Similarly, another non-reinforced preference change paradigm that did not rely on external reinforcements was found in work showing preference modification following previous choices^7^. These studies provide a strong theoretical framing for efficacy of interventions such as advertisements on preferences and how life choices affect future decisions ^8,9^, above and beyond external reinforcement-based manipulations ^10,11^.

A decade ago, a unique procedure named the Cue-Approach Training (CAT) ^12^ task has been developed as a reliable means to change preferences without external reinforcements. The CAT procedure is a multiphase task with an initial training session on individual items and a subsequent binary choice phase. The first studies with the CAT procedure, showed that preferences for snack food stimuli could be modified via an association between images and a neutral cue and rapid motor response^12–14^. The task included a Go/NoGo training phase, during which a set of snack food stimuli were presented individually on the screen. Some snacks were associated with a Go cue, to which participants were required to respond with a rapid button press response (Go stimuli). Another (larger) set of snack food stimuli were passively presented for the same exposure period, without a cue or response (NoGo stimuli). The training phase included several repetitions (runs) in which participants could learn the association of individual Go stimuli with the cue and response. In the subsequent post-training probe phase Participants demonstrated enhanced preferences for the Go stimuli over NoGo stimuli. This phase included binary choices between two snacks for actual consumption and these two snacks had a similar initial subjective value ^12–14^.

Multiple studies with the CAT procedure exemplified its efficacy in modifying preferences for a wide range of stimuli beyond snack food items, including healthy food items, faces, positive affective stimuli and fractal art stimuli^15–18^. Overall, across dozens of studies with different types of stimuli, participants consistently demonstrated enhanced preference for Go stimuli over NoGo stimuli of similar initial value^12–21^. Studies that examined the long-term maintenance of the CAT effect in additional follow-up sessions, found that the preference modification effect persisted for months without any additional training sessions^12,15,18,21,22^. The fact that a preference modification effect was observed following a training procedure with no external reinforcement or feedback, on individually presented items, inspired the idea that CAT relies on a non-reinforced valuation pathway, putatively involving attention and motor neural circuits^5^.

Several mechanisms have been suggested to underlie non-externally reinforced paradigms. First, the preferences-as-memory theoretical framework suggests that value could actually be framed as retrieval of relevant knowledge about alternatives from memory^23^. Relatedly, studies showed that Go stimuli that were better remembered were also more preferred and that memory was positively correlated with likelihood of choosing Go stimuli^19,21^. Attention has also been shown to play a key role in value-based decision making both on its own and in synergy with memory^24,25^. Some support for the involvement of attention was found in eye-tracking studies with CAT, which found that during the probe phase, participants view chosen Go stimuli more ^12,13,18,26^, suggesting that increased gaze time during the probe phase indicated attentional evidence-gathering process, which increased the likelihood of choosing the Go stimuli over NoGo stimuli ^27–29^. However, most of these studies ^13,18,26^ did not find enhanced gaze for Go stimuli when they were not chosen, undermining the hypothesis that enhanced attention could drive the CAT preference modification effect on its own.

However, these previous mechanistic views have focused on the probe phase where preferences are being expressed, whereas preference is changed during the training phase. During training all stimuli are presented individually for the same duration, thus mere exposure or viewing time alone could not account for the preference modification effect. Studies that examined the necessary features of the training task discovered that simply responding slowly to the Go stimuli was not sufficient to induce preference changes, as preference modification did not occur in a training version in which the Go cue was presented immediately with the Go stimulus allowing a full 1 second to respond. Similarly, training with all Go cue items presented in a block of Go stimuli^13^ did not yield a preference change effect. These findings suggested that simple attention or motor response are not sufficient, and that the rapid motor response during CAT and the motor preparation learning are crucial factors for non-reinforced preference modification with CAT.

The critical question that persists is what the mechanisms are, that operate at the individual item level without external reinforcements to induce the subsequent observed behavioral change in the binary choice phase. A prominent clue as to the underlying mechanisms during training were identified in a recent imaging study with CAT^18^, which found that individual differences in the preference modification effect (measured during the binary choice probe phase) correlated with increased neural activity during training within the supplementary-motor cortex and the striatum. These regions are associated with motor-planning^30,31^ and reinforcement-based learning^32–35^, respectively. The striatum has been suggested to be key neural hub for learning thanks to its unique dual-role in both reward processing and motor regulation networks ^36,37^. The involvement of the striatum linking non-reinforced training and subsequent preference change, could putatively indicate the presence of internal reinforcement ^38^ during the training phase, particularly in the absence of external feedback.

These neuroimaging results, showing that enhance striatal activity during training correlated with preference modification were the key inspired the current work’s hypothesis. We utilized the unique structure of CAT that trains individual items, which allows us to shed light on valuation mechanisms at the individual item level. An important aspect of the CAT task is that training is performed at the individual item basis, and preference change is tested in a subsequent binary choice phase. Therefore, we surmised that identifying individual differences in motor learning during the training phase could serve as the key factor to explain differential non-reinforced behavioral change effect following CAT.

We tested our hypothesis by developing a novel Bayesian computational model with an individualized learning marker based on training motor-response data on single items and testing its association with the preference modification effect observed in the subsequent probe phase. For the development of the new computational framework, we utilized 29 different experiments that used the CAT procedure. Then, to examine the causal impact of our newly proposed mechanism we designed a new non-reinforced training procedure which aimed to directly impact learning and establish a framework for understanding the intrinsic mechanisms underlying non-reinforced preference modification through motor-learning.

The current work thus consists of two main parts. The first part includes a meta-analysis study, whereby using data of 864 participants from 29 previous CAT experiments, we devised a novel marker for learning. By combining multiple samples with the CAT task into one meta-analysis, we were able to create a unique dataset of a standardized non-reinforced preference modification task. To identify an individualized marker for learning, we first examined RT patterns during the training phase in an exploratory analysis and built a computational model of these patterns using a Bayesian computational framework. The Bayesian model included an individualized parameter, which modeled a transition from cue-dependent responses to anticipatory responses and was used as a foreseeing marker of individualized learning. We examined the association of the individualized learning marker with individual differences in preference modification, as measured in the subsequent binary-choice probe phase. We further examined the hypothesis that the mechanism of non-reinforced preference modification is operating both at the participants-level (i.e., some participants learn better than others), as well as on a more granular item-level (i.e., within participants, some stimuli are better learned than other). We expanded our computational model to capture variability in within-participant item-level learning and examined its association with subsequent choice pattern.

In the second part of this work, we implemented the findings from the Bayesian model to develop a novel CAT design and directly manipulate the computational marker in order to demonstrate a causal direction between the new model and preference change. In the new design, we used two different cue-contingency conditions to manipulate learning difficulty. We hypothesized that the cue contingency manipulation will affect the motor response and will be captured by the individualized learning marker devised in the first part, as well as predict subsequent preference modification in the probe phase. The second part included one preliminary experiment (*n* = 20), which was followed by a larger pre-registered replication study (*n* = 59). All the hypotheses, experimental design and analyses plans were preregistered before data collection of the replication study began (https://osf.io/nwr4v). By manipulating the task design to induce differential behavioral modification effect, we aimed to establish a causal relationship between the proposed cognitive mechanism and the preference modification effect.

By identifying an individualized computational marker for learning in CAT, the current work provides empirical evidence that non-externally reinforced preference modification occurs at the individual item level via motor-learning cognitive mechanisms. We establish a reliable quantification of this individualized learning, without the need to explicitly ask participants to reveal their preference overtly. The ability to exemplify a marker for non-externally reinforced preference changes at the individual item level, this work will shed light on the putative mechanism of this effect. Furthermore, it will allow us to identify learning patterns both between-participants and within-participant, at the individual item level. A computational marker for learning holds the potential to passively monitor learning in real-time and support the development of novel closed-loop interventions.

### Study 1: Meta-analysis of CAT studies

In the first part of the current work we performed a meta-analysis of 29 CAT experiments with a total of *N* = 864 participants, which had been conducted by our research group and colleagues (see methods for sample size, publication origin and demographics of the individual experiments). All experiments included the three main phases of the CAT procedure: initial preference evaluation, cue-approach training, and a preference modification probe task (Figure 1; see detailed description in the Methods section). In the first phase of the task, initial baseline preferences for a set of stimuli were evaluated in an auction procedure ^39^, when snacks were used, or a forced choice task ^15^ for all other stimulus types. Following initial preference evaluation, in the cue-approach training phase, approximately 30% of the stimuli were consistently associated with a neutral Go cue (Go stimuli), to which participants needed to respond with a rapid button press, while the rest of the stimuli were presented passively without cue or response (NoGo stimuli). Participants were instructed to wait for the Go cue and respond as fast as they could, once they perceived the cue. In the final probe phase, preference modification following CAT was evaluated. Participants were asked to indicate their preferred stimulus out of pairs of stimuli with similar initial value, in which one of the two stimuli was a Go stimulus and the other was a NoGo stimulus.

**Figure 1.**
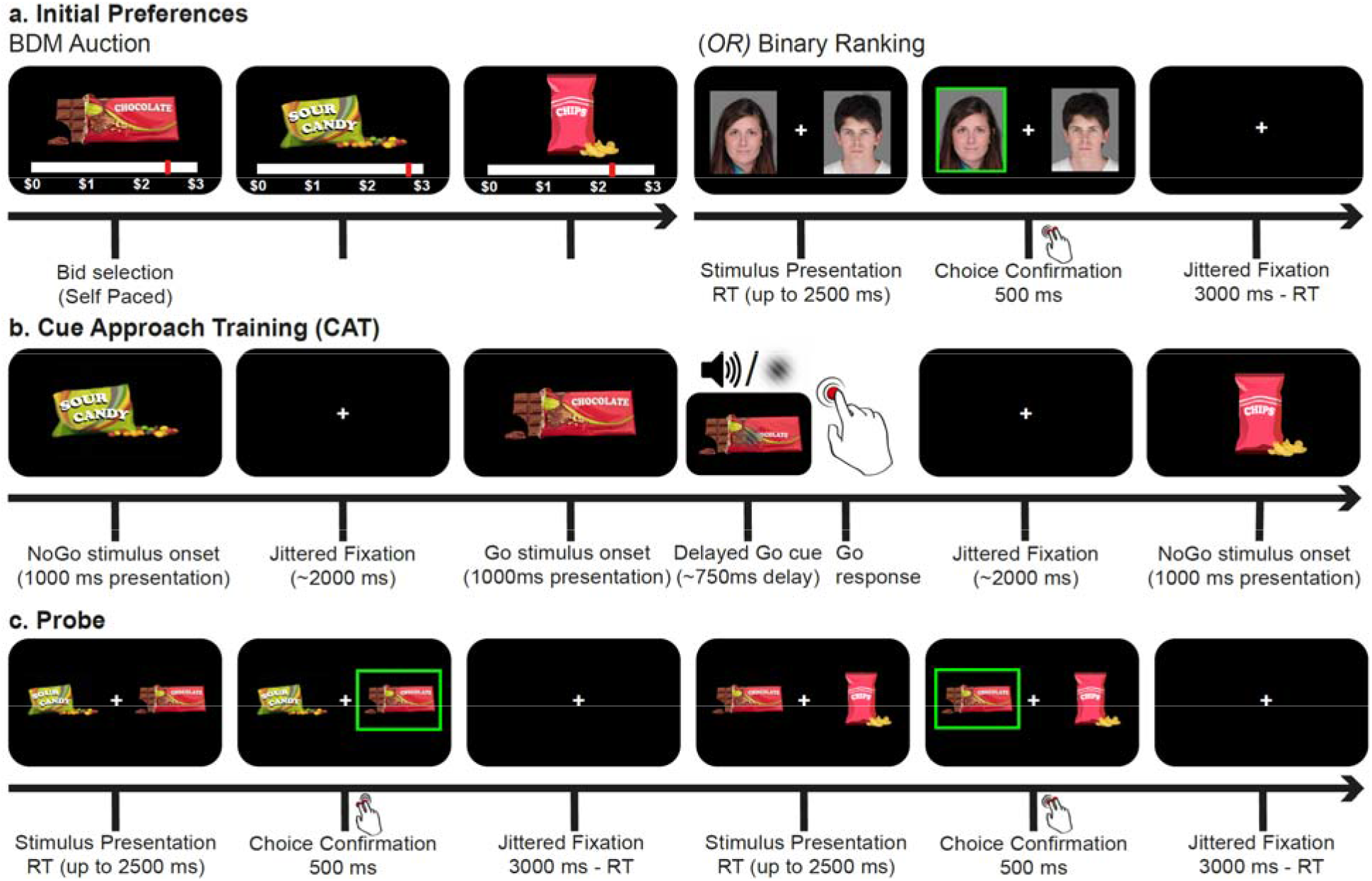
General outline of the three main procedural components of the Cue-approach training (CAT) paradigm. (a) Initial preference evaluation task. Baseline preferences for all stimuli were evaluated either with using a Becker-DeGroot-Marschak (BDM) auction (for consumable stimuli) or using a forced choice binary ranking task (for non-consumable stimuli). (b) In the cue approach training (CAT) task, approximately 30% of stimuli were presented in association with a delayed cue (auditory or visual) to which participants responded with a rapid button press (Go stimuli). All other stimuli were presented without cue and response (NoGo stimuli). (c) In the probe phase, preference modification was evaluated using a binary forced choice between pairs of stimuli of similar initial value, where one was a Go stimulus and the other a NoGo stimulus. Face images are included with permission from the copyright holder ^40^.

The meta-analysis study focused on identifying response patterns in the training task which could indicate learning efficacy and predict the behavioral change measured in the subsequent probe task. We hypothesized that faster Reaction Time (RT) could indicate improvement in learning.

## Results

### RT analysis of CAT

The training task in the different experiments consisted of 8-20 training runs (the number of training runs varied between experiments and identical within each experiment). During each run, all Go and NoGo stimuli were presented once individually on-screen. During NoGo trials, stimuli were presented without a Go cue and required no response. Whenever Go stimulus appeared (approximately 30% of trials), a Go cue was followed, to which participants were asked to respond with a rapid button press, before stimuli offset. Participants were instructed to respond *only* when they hear or see the cue. Each trial started with stimulus onset, which was presented on screen for 1000ms. During Go trials, a Go cue appeared approximately 750ms following stimulus onset (thus leaving participants approximately 250ms to respond). The cue onset changed according to participants’ performance, so that each successful Go response was followed with an increase in Go onset time (thus leaving less time to respond and making the next trial more challenging; see detailed description in the methods section).

To identify unique response patterns, we performed an exploratory analysis of the RT distribution during Go trials. Before analyzing RT in the CAT task, invalid trial trials were excluded. Resulting in the exclusion of missed trials, in which participants failed to make a response within the allocated 1500ms from stimulus onset timeframe (0.90% of trials), and trials in which the response was shorter than a threshold of 100ms (0.02% of trials), as these trials are likely to be indicative of inattentional response. In total, 1731 trial out of 188,224 Go trials were excluded (0.92% of trials).

Examining the RT density distribution in the training task as a function of training run, revealed a distinct pattern, wherein as training progressed, participants’ RTs were less homogeneous and started to form a growing peak of early RTs (Figure 2a). In all 29 experiments, the Go cue changed in each trial, according to participants’ performance (see methods), thus, a more informative measurement of RT was the time from cue onset (RT minus cue-onset), referred to here as *effective RT*. In earlier training runs, when the Go stimuli were associated for the first time with the Go cue, most effective RTs were clustered around a unimodal center, approximately 300ms following the Go signal onset (effective RT *M* = 293ms) with 99% of effective RTs larger than 145ms. We use here this empirical quantile of 145ms as threshold to define rapid *anticipatory responses*, as these responses can be elicited when the participants predicted a go cue would appear. As training progressed, a consistent pattern appeared in the effective RT data - the main peak of the RT distribution was reduced, while a growing portion of participants’ responses consisted of faster anticipatory RTs, in many cases preceding the Go cue onset (Figure 2b). For example, while the proportion of anticipatory responses comprised only 1% of RTs in the first training run, they comprised 2.7% of RTs by the 5^th^ run, 12.2% of the trials by the 10^th^ run, 20.5% of trials by the 15^th^ run, and finally 29.4% of trials in the final 20^th^ run (aggregated across all samples).

**Figure 2.**
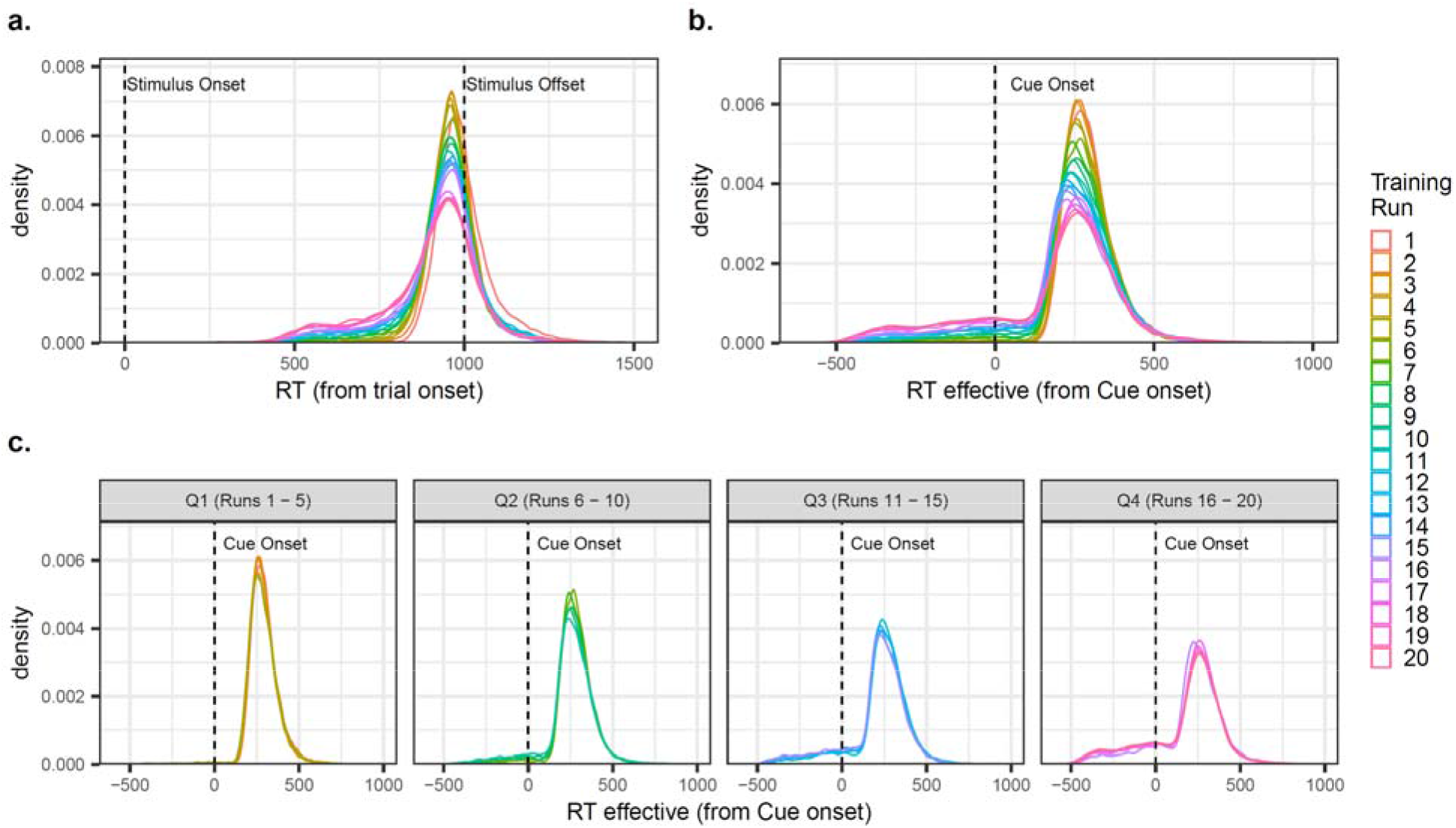
Reaction time distribution of data pooled from meta-analysis with 29 studies. Density plots of (a) RTs time-locked to the trial onset and (b) RTs time-locked to the varying Go signal onset (*effective RT*). Color indicates training run (repetition). As training progressed, th RT distribution shifted from a cue-dependent unimodal distribution to left-tailed distribution with increasing proportion of early anticipatory responses. (c) a clear differentiation in effective RT is apparent when splitting the data (from Figure 2b) to four quartiles, according to the training run (each quartile indicating five consecutive training runs).

Thus, RT data suggest a temporal-dependent response pattern. As training progress, participants tended to rely less on the Go cue (late cue-dependent RTs) and generated more early anticipatory responses. Interestingly, the mode of the RT distribution seemed to remain stable, and only the relative proportion of the two distributions changes. We hypothesized that this transition pattern from the cue-dependent response to a stimulus-triggered anticipatory response reflected a process of learning about the stimuli that could be associated with behavioral change in the later preference probe. Thus, we next sought to develop a model to characterize the strength of this effect within-individual.

### Individualized-learning computational model

Based on the unique RT pattern observed in the exploratory analysis of CAT task data, we developed a novel computational Bayesian model. To capture the form of RT distribution with rapid anticipatory responses and slower cue-dependent responses, we modeled the effective RT as a mixture of two Gaussian distributions, with two different means (*µ*_1_ and *µ* _2_ free parameters), two standard deviations 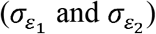, and a mixture proportion (Θ _*i,t*_) indicating the relative mixture proportion for participanti at trialt. The model was implemented with the Stan probabilistic programming language ^39^, which optimized the model parameter estimates using a Markov-chain Monte Carlo (MCMC) approach. See methods section for a detailed description of the model and Supplementary Code S1. The first Gaussian was restricted to have lower mean than the second Gaussian to maintain their correspondence across different MCMC sampling chains (i.e., *µ*_1_, 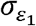 would correspond with the early anticipatory RT distribution). To capture the gradual change in anticipatory responses over training, the mixture proportion was modeled as a time-dependent variable, expressed by the following formula:

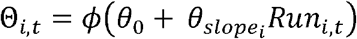

Where the mixture probability Θ _*i,t*_ for participant_i_ at trial_t_ was modeled as a linear function (with a fixed *θ* _0_ intercept and random participant-level 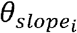 free paramter) of training run (where the *Run*_*i,t*_ ranged between [0,1]). The linear trend was scaled non-linearly to probability-appropriate values of [0, 1] range using a normal distribution cumulative distribution function (normal CDF; denoted with *ϕ*), resulting in a sigmoid-like link function. See further details in the methods section.

To associate all learning with a single parameter and a simple interpretation of 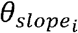 free parameter, we chose to fix the *θ* _0_ intercept term, which represents the mixture proportion at the very first run (*Run*_*i,t*_ = 0). The *θ* _0_ intercept term was fixed at -3.1, corresponding with Θ_*i,t*_= *ϕ* (− 3.1) ≈ 0.1% mixture proportion at the very first run for all participants. Thus, higher 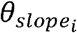 parameter estimate indicates an individual participant with faster transition from slow cue-dependent RTs to fast anticipatory responses, which corresponds with larger Θ _*i,t*_mixture proportion at the very last run. For example, parameter estimate of 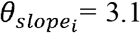, indicated that at the last run (*Run*_*i,t*_ = 1), the mixture proportion of anticipatory responses would be *ϕ*(0) = 50%, while 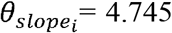 would correspond with mixture proportion of *ϕ*(1.645) ≈ 95%, at the last run. We hypothesized that this individualized parameter would be indicative of improved learning in the training phase of CAT and would also be positively associated with stronger preference modification effect, measured in the subsequent probe phase.

Fixing the expected anticipatory responses to very low number at the first run, was an assumption we chose to adapt in the current work to be able to quantify different participants’ learning using 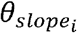 as a single comparable parameter which captured all aspects related to learning. If *θ*_0_ intercept term would also have been left as a free parameter, we would have needed to account for both parameters and their interaction to define a computational learning marker.

After running four independent chains, the Bayesian model converged to a stable solution (see Supplementary Fig. S1). The converged model estimated effective RTs as a time-dependent mixture of two RT distributions: one of early anticipatory responses (*µ*_1_ = 107.50ms, 95%CI [103.77, 111.21], 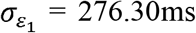, 95%CI [274.13, 278.42]) and late cue-dependent responses (*µ*_2_ = 286.12ms, 95%CI [285.67, 286.58], 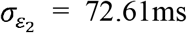, 95%CI [72.23, 73.00]). Overall, participants demonstrated an increase in proportion of anticipatory responses as training progressed, as manifested in the positive group-level parameter (*θ*_*slope*_ = 4.19, 95%CI [3.84, 4.54]), with variation between participants (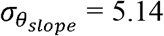, 95% CI [4.85, 5.46]), indicating some participants made faster transitions and some generated nearly only cue-dependent responses (non-positive 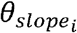 estimates).

Posterior predictive checks (simulated distributions of RT based on estimated model parameters) revealed a good fit of the model to the actual data. Simulated posterior distributions recreated the patterns observed empirically, showing more rapid transition to anticipatory responses in participants with higher 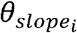 parameter estimate (Figure 3; Supplementary Fig. S2).

**Figure 3.**
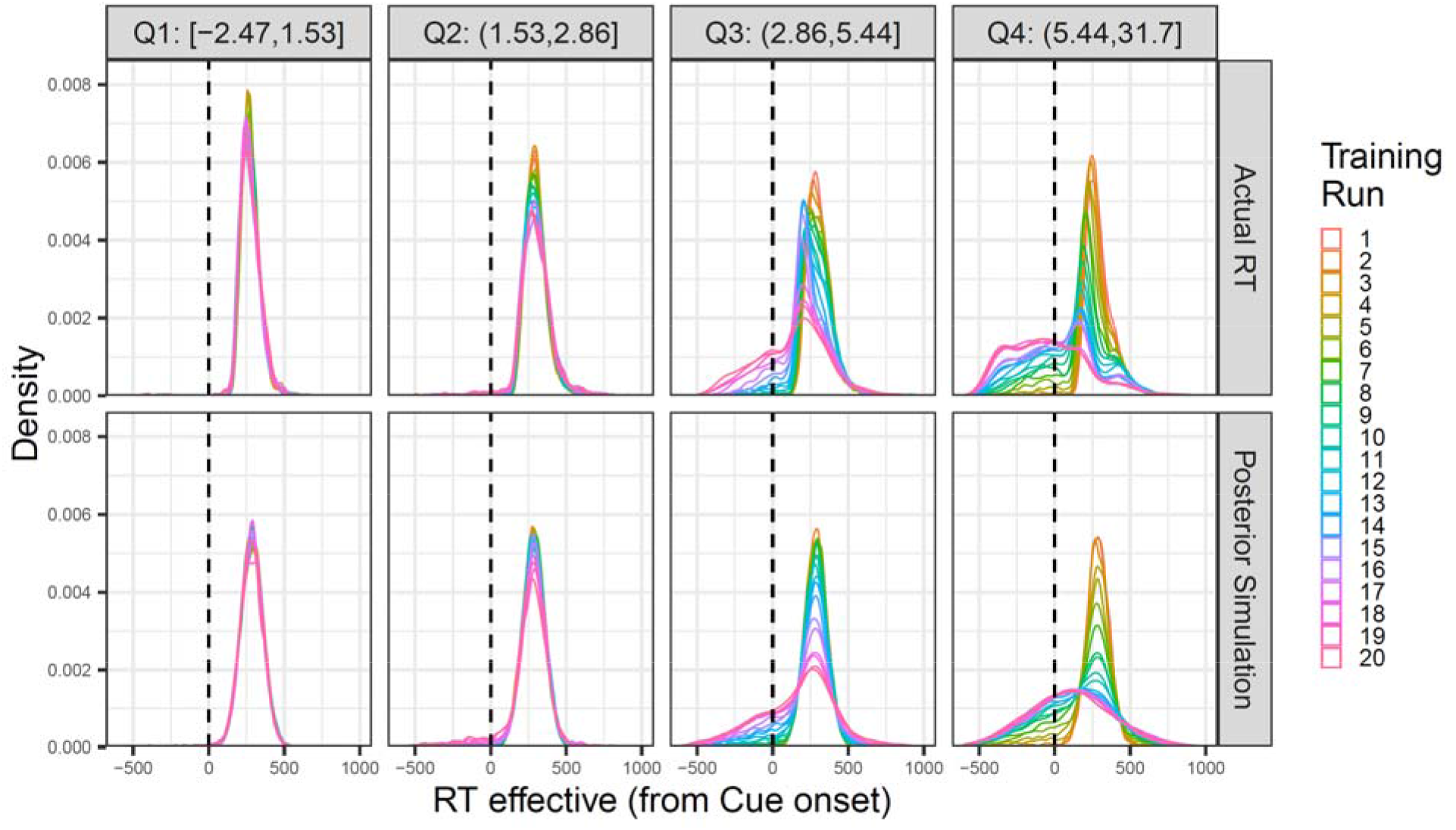
Actual RT distributions versus simulated posterior distributions, by quantile group. Participants were categorized into four equal quantile groups, according to their parameter estimates (denoted here as Q1-Q4; columns). Participants with higher parameter estimates were characterized with faster transition to anticipatory responses (top row). Posterior simulated RT distributions using mixture of Gaussians (bottom row) recreated relatively well this transition pattern. Vertical dashed line represents cue-onset. See also Supplementary Fig. S2 for more detailed comparison.

### Learning parameter association with choices

Previous findings with CAT consistently showed that preferences for Go stimuli were enhanced following CAT, as manifested in choice behavior during the probe phase. When presented with a choice between a Go stimulus and a NoGo stimulus of similar initial value, participants consistently chose the Go stimulus ^12,14–20,22^. To test the hypothesis that this probe performance is associated with RT patterns during CAT, we examined whether variation in the slope parameter fit to RTs correlated with probe performance. Note that, under the model, the slope parameter controls the final proportion of anticipatory responses, which we understood as a measure of the extent of learning about th stimulus achieved by the endpoint of CAT. To evaluate the association of the parameter with preference-change effect following CAT, we analyzed the proportion of probe trials in which participants chose the Go over the NoGo stimulus, as a per-participant linear function of (mixed logistic regression model, including random intercept and slope terms for the 29 experiments; see methods).

The meta-analysis showed was positively associated with the preference modification effect – i.e., participants with higher parameter estimate, also demonstrated greater odds of choosing Go stimuli (OR = 1.04, 95% CI = [1.02, 1.06], *Z* = 5.17, *p* = 4.6E^-7^; two-sided mixed model logistic regression; Figure 4). The model’s intercept was significantly greater than zero, i.e., even when extrapolated to very low value, the model forecasted enhanced preference for Go stimuli (intercept odds = 1.26, 95% CI = [1.18, 1.34], *Z* = 7.03, *p* = 2.0E^-12^).

**Figure 4.**
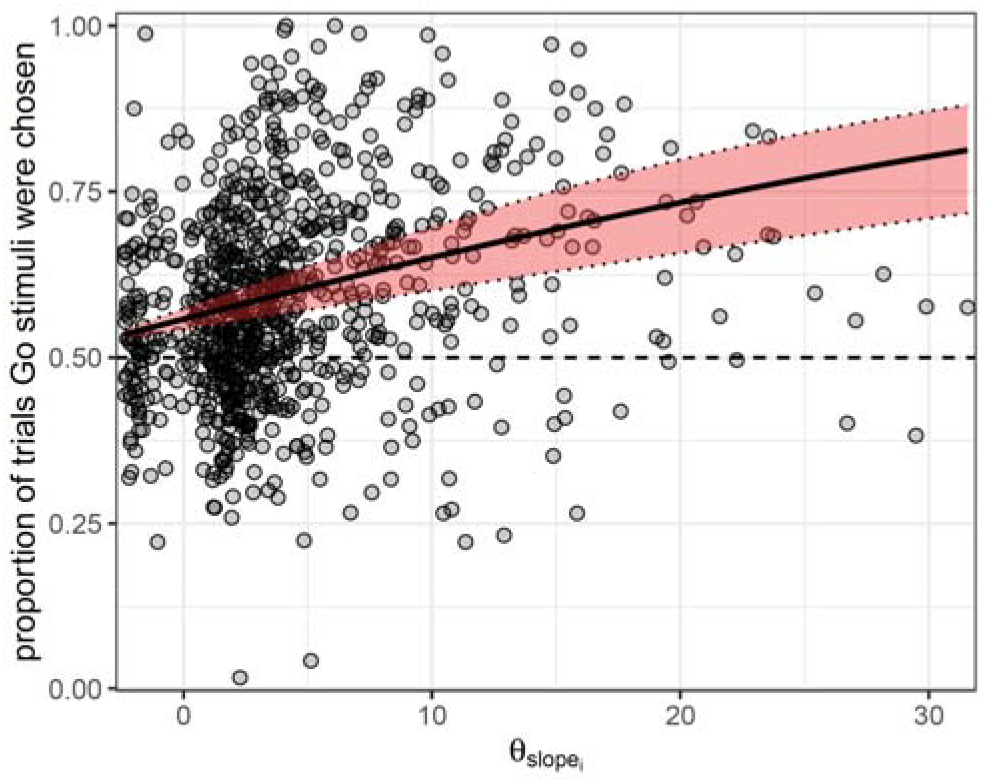
Meta-analysis results - computational marker and preference modification effect. Participants who transitioned faster to anticipatory response during cue-approach training (larger estimates) also demonstrated stronger preference modification effect (proportion of trial Go stimuli were chosen). Trend line and surrounding red margins represent estimated preference modification effect and 95% CI, respectively (mixed model logistic regression). Dots represent individual participants.

The estimated variances associated with the fixed-effect terms, random-effect terms and residuals were used to evaluate two scores for the generalized (logistic) linear mixed mode: a marginal (representing the relative proportion of variance associated with the fixed-effects) and a conditional score (representing the relative proportion of variance associated with the fixed-effects and random-effects; see more details in the methods section). The fixed effect accounted for = 0.311 of the variance. With the random intercept and slopes, the fixed and random effects combined accounted for = 0.828 of the overall variance.

### Additional model validation

In a post-hoc analysis (see Supplementary materials), we examined the explanatory power of 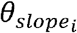 above and beyond a simpler RT-based marker. Using the proportion of anticipatory responses (RTs which were faster than the top 1% of RTs in the first run) each participant made as an alternative marker, we found the simpler marker was also associated with subsequent probe choices. Furthermore, 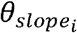 showed no significant contribution above and beyond this simpler RT-based marker.

In an additional post-hoc analysis, we used an alternative Bayesian model in which RT were modeled using log-normal distributions instead of Gaussians (see Supplementary). This new model better captured the right-tail shape of RTs (Supplementary Fig. S13) and replicated similar correlation pattern between 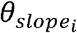 parameter estimate and Go stimuli choices during probe (OR = 1.05, 95% CI = [1.03, 1.07], *Z* = 6.05, *p* = 1.4E^-9^; see Supplementary Fig. S14).

### Stimulus-specific learning parameter

While the previous model aggregated the data within participants, in fact, each participant has learned the Go cue association with several different Go stimuli. Thus, it is possible that learning was not uniform across the entire training stimuli, meaning every participant had a slightly different learning rate for each of the Go stimuli she or he encountered. In an additional model, we aimed to expend our model by examining behavior on a more granular scale – focusing on the variability in RT and choice for individual Go stimuli within participant. Accordingly, the computational model was elaborated by decomposing the per-subject learning parameter 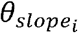 into a set of subject- and stimulus-specific parameters 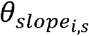, indicating the transition speed to anticipatory RTs of each Go stimulus_s_ encountered by participant_i_ during training. The model was otherwise identical in design (see methods). This reanalysis of the data was conducted after the collection and analysis of the following two studies which are reported in the current work.

The reanalyzed model converged around slightly different parameter estimates: estimated early anticipatory RT distribution was characterized with lower mean (*µ*_1_ = 88.30ms, 95% CI [84.1, 92.34], 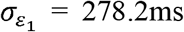, 95% CI [275.84, 280.65]), late cue-dependent responses were similar (*µ*_2_ = 289.31ms, 95% CI [288.82, 289.79], 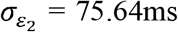, 95% CI [75.23, 76.05]), and the 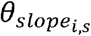 parameters tended to be lower but with greater variability (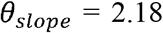, 95% CI [2.01, 2.34], 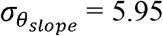, 95% CI [5.76, 6.16]; see Supplementary Fig. S3).

Similarly to the previous logistic regression analysis with per-participant parameter, using the current analysis’ per-stimulus parameters also significantly predicted choices of the corresponding stimuli in the subsequent probe phase (Intercept: Odds = 1.45, 95%CI [1.36, 1.54], 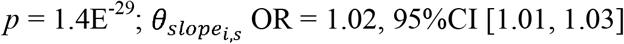, Z = 3.65, *p* = 2.6E^-4^; two-sided mixed model logistic regression; see Supplementary Fig. S4). Thus, Go stimuli which were characterize with larger proportion of anticipatory responses were also chosen more during the probe phase. The models’ fixed effects accounted for 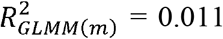 of the total variance and the fixed effect with mixed effects accounted for 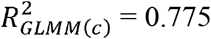 of the total variance.

However, this overall effect comprises both between-participant and within-subject, between-stimulus contributions. To examine the unique contribution of 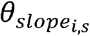 above and beyond participant-level 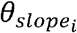 parameter (which was evaluated in the previous analysis), in a third logistic regression model we included both per-participant and per-stimulus parameters from the two different Stan models, as independent variable (and their random effects; see methods) to predict choices during probe. Our results indicate that each parameter provided some unique contribution above and beyond the other parameter (Intercept: Odds = 1.37, 95%CI [1.27, 1.47], 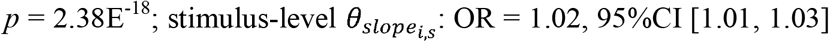, Z = 3.34, *p* = 4.2E^-4^; participant-level 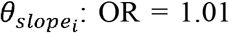, 95%CI [1.00, 1.03], Z = 2.07, *p* = 0.019). Thus, stronger learning of anticipatory RTs, both per-individual and per-stimulus within individual, were associated with increased odds of choosing Go stimuli.

### Interim discussion

Examining RT patterns during CAT revealed for the first time a distinct time-dependent RT pattern, in which participants gradually transition from slow cue-dependent responses to rapid nticipatory RTs. Using Bayesian modeling, we were able to quantify this RT transition pattern with stable parameter estimates, of which the 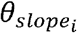 parameter provided a promising computational marker to evaluated individualized differences in learning during training. Furthermore, as expected, this computational marker was also found to be associated with the preference modification effect in the subsequent probe phase, as participants with a more robust learning marker also demonstrated stronger preference modification effect. Examining the data on a stimulus-level learning parameter, demonstrated an improved explanatory power for the variability in probe choices, above and beyond the wider participant-level parameter. Since the computational marker was calculate based on measurements that preceded the probe phase, it holds a potential to provide predictive marker for future preference modification.

Several challenges were noted in modeling the original experimental data with the new computational model, largely owing to the fact that the original experimental procedures were not designed with the present analyses in mind. First, in the training task, Go cue onsets were derived based on performance using a staircase procedure. Participants who performed well and responded rapidly were presented in the next trials with a more challenging cue, appearing later into the trial, while participants who performed poorly were exposed to earlier occurring (less difficult) cues. Consequently, anticipatory responses were not time-locked to the same event as the late cue-dependent responses. While a model that uses RTs time-locked to the stimulus onset could potentially model anticipatory responses shape well, using a model that is agnostic of the cue-onset time, would result in difficulty to differentiate between cue-dependent and anticipatory responses due to the changing cue onset. For example, a response at 750ms could be attributed to cue-dependent response of a participant with very poor performance which was presented with a cue at 500ms, or an anticipatory response of a well-performing participant that encountered a challenging 800ms cue onset delay. Thus, making on the surface the two response patterns hard to distinguish and introducing a potentially confounding source of variation across participants. Therefore, in the current study we use RTs time-locked to the cue onset, which improved the distinction between cue-dependent and anticipatory responses, at the expense of less accurate tracking of the anticipatory response distribution shape.

Some lack of fit of the predicted data could also be indicative that the model’s simplistic assumption that anticipatory RTs are normally distributed around the same mean for all participants is unlikely. This challenging effect may have led to some counter-intuitive results; for instance, due to high variability in anticipatory responses, the fitted model predicted that very slow responses (RT effective > 500) would be more likely in the fastest learners’ late runs, compared with earlier runs or with slower learners. In an additional post-hoc analysis (performed after the initial write-up of this manuscript), we modeled RT using a log-normal distribution, which better captured the skewed right-tail shape of RTs, which replicated the conclusions of the current design. Nonetheless, the large heterogenous data structure (with over 800 participants from 29 different experiments) provided a computational challenge when trying to fit more complex models (e.g., with random effects per participant on this parameter or using non-normal distributions). The current model was selected as a reasonable compromise that provided sensible results. The fact that using an additional alternative modeling approach resulted in similar conclusions as those gathered with the preregistered Gaussian model, provided vital evidence that the conclusions of the current study are stable above and beyond implementation choices.

Furthermore, the task instructions of CAT specifically mentioned that participants are required to respond *after* cue-onset. An additional crucial factor which could not be controlled in the current study was how rigorously participants complied with this instruction. It is possible that some participants might have learned the stimulus-cue association well and could have produced anticipatory responses but were reluctant to deviate from the task’s guidelines. If so, their 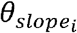 would not reliably reflect their actual learning of the task. It is important to note that although the logistic regression results showed a significant positive association of choices with participant-level and even stimulus-level individualized learning parameters, significant preference change remains even when the model extrapolated for no learning effect. The effect sizes reported above are quite modest and in a post-hoc analysis they were not found significantly better compared to predicting choices with a simpler RT-based marker. Thus, suggesting of additional contributions to the choice pattern which are not otherwise accounted for by the computational marker. The small effect sizes might be due to the inaccurate measurement of additional constructs within the computation parameter, such as the instruction adherence and interaction with performance-based dynamic cue onset, which were proposed here.

Despite these drawbacks, our unique model provided a prospective computational marker for learning and subsequent preference changes. Based on the promising results of Study 1 meta-analysis and considering the challenges raised above in fitting the computational model to the CAT task data, in Study 2, we devised a novel design of the CAT procedure, which was tested with two independent experiments, a smaller preliminary study (*n* = 25) and a larger pre-registered replication study (*n* = 59). In these new experiments we addressed the limitations of the previous study design. We also modified the training design to answer a more directional hypothesis which does not only examine correlational association, but also try to provide evidence for more consequential directionality, by inducing a differential behavioral change.

### Study 2: Novel CAT design – preliminary and replication experiments

A novel experimental design was tested in two pre-registered experiments - a preliminary experiment and a larger replication experiment. Based on the conclusions from Study 1, the CAT procedure and instructions were altered to optimize the task for the Bayesian computational framework and manipulate behavior in accordance with the hypothesized cognitive mechanism.

In the training task, the Go cue onset time was fixed at 850ms from stimulus onset and participants were clearly instructed that they were permitted to make anticipatory responses before cue onset. Furthermore, based on the conclusions from the meta-analysis study, we introduced within the training task a manipulation of the predictability of the stimulus-Go cue contingency, which we expected would affect the 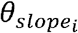 parameter and subsequently also manipulate the preference modification effect. Of the Go stimuli, half of the stimuli were always associated with the Go cue (100% contingency condition), while the rest of the Go stimuli were only associated in half of the presentations with the Go cue (50% contingency condition). The rest of the stimuli were never associated with a Go cue (NoGo stimuli; See methods section for detailed description of the new design).

We hypothesized that in the 50% contingency condition, learning the association of Go stimuli with the Go cue would be more challenging, and would therefore be identified by a lower 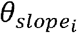 learning parameter. We also hypothesized that manipulating learning efficacy during training would therefore induce a differential preference modification effect in the subsequent probe phase, with more robust preference modification for Go stimuli trained in the 100% contingency condition, compared with the more challenging 50% contingency condition.

The new task design was first tested in a preliminary (not preregistered) study with *n* = 20 valid participants, aimed to evaluate the efficacy of the new task and the replicability of the meta-analysis conclusions. Following the preliminary experiment, we run an additional pre-registered direct replication experiment with a larger sample size (*n* = 59), a size chosen based on the preliminary experiment results. As the two experiments were identical in design, they are reported together in all sections.

## Results

### Reaction times in the training task

As in the meta-analysis study, RT patterns of the training task were examined. Before examining RT patterns, we removed Go trials where participants did not respond at all (preliminary exp. = 6.33%. replication exp. 5.97% of Go trials) or trials with an unlikely fast RT (RT< 250ms, preliminary exp. = 0.31%, replication experiment 0.21% of trials). In total, 636 (of 9600; 6.62%) and 1747 (of 28,320; 6.17%) of trials were excluded from the preliminary and replication experiments, respectively.

In both experiments, we were able to replicate the RT pattern observed in the meta-analysis study – while in early training runs, participants mostly relied on cue-dependent RTs, as training progressed, an increased portion of RTs were of earlier anticipatory RTs (Figure 5). Examining mean (*SD*) effective RT at Run 1, revealed similar RTs in all contingency conditions - preliminary exp.: *M*_50%*Cont*_. = 326.88ms (85.13ms), *M*_100%*Cont*_. = 336.60ms (64.39ms); replication exp.: *M*_50%*Cont*_. = 322.49ms (63.68ms), M_100%*Cont*_. = 319.90ms (78.88ms). As training progressed, participants were able to anticipate the cue and respond before cue-onset. Examining the differential pattern between the 100% contingency condition and the 50% contingency condition showed, as expected, that participants initiated more anticipatory responses in the 100% contingency condition, compared with the 50% contingency condition, resulting in faster mean-RTs at Run 20 - preliminary exp.: *M*_50%*Cont*_. = 253.27ms (187.65ms), *M*_100%*Cont*_. = 175.55ms (245.5ms); replication exp.: *M*_50%*Cont*_. = 256.57ms (188.94ms), *M*_100%*Cont*_. = 81.39ms (271.99ms); see Figure 5.

**Figure 5.**
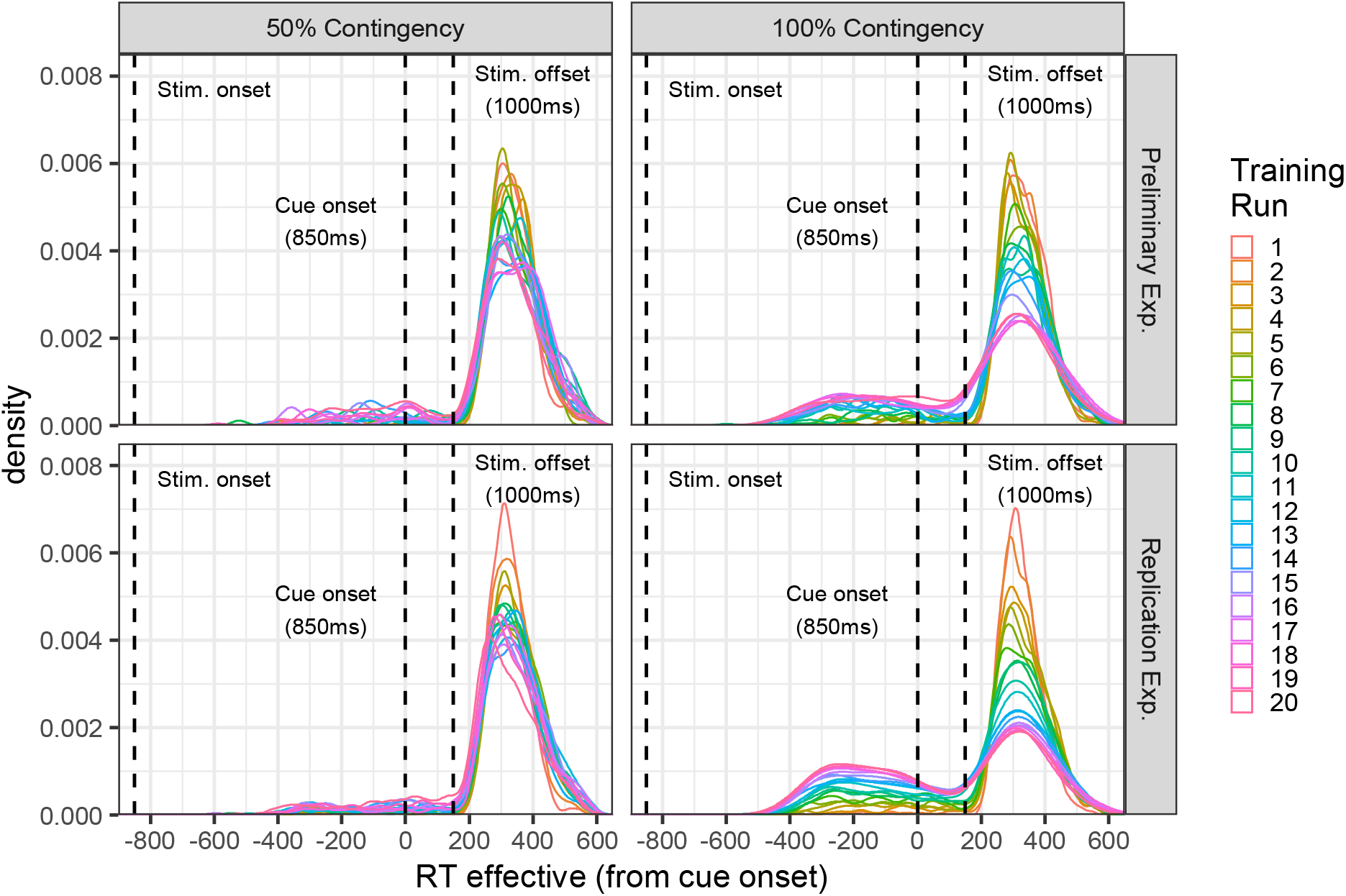
Density plots of effective RT distribution (time-locked to cue onset) in Study 2 (preliminary experiment on top row and replication experiment on bottom row). The Go cue onset was fixed at 850ms from stimulus onset - vertical dashed lines represents the trial onset (time 0), cue-onset, and trial offset (850ms and 1000ms from trial onset, respectively). As training progressed (indicated by the training run in color), responses were less homogeneous following cue onset, with increased proportion of anticipatory responses. The transition to anticipatory responses was more robust in the 100% contingency condition (right column) compared with the 50% contingency condition (left column).

#### Computational marker of learning

As in the meta-analysis study, RTs within each condition were modeled with a time-dependent mixture model, implemented with Stan Bayesian framework. RT data within each condition (100% contingency and 50% contingency) were fitted using two Gaussian distribution parameters (two means and two SDs). A 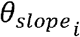 time dependent parameter was modeled for each condition and determined the rate of transition from cue-dependent responses to anticipatory responses of individual participants. Since the Go cue in Study 2 was fixed at 850ms, the effective RT (relative to stimulus rather than cue) measurement perfectly correlated with actual RT; thus, we used only the actual RT in the model.

In both the preliminary and replication experiment, the computational model converged around similar hyperparameter estimates (see MCMC trace plot and posterior simulations fit in Supplementary Fig. S6 and Supplementary Fig. S7, respectively). Cue-dependent responses were characterized with later mean RT and smaller variance, compared with earlier anticipatory responses (preliminary experiment: *µ*_1_ = 735.07, 95%CI [677.14, 792.87], 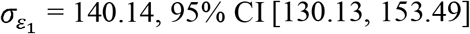, *µ*_2_ = 1190.23, 95% CI [1188.46, 1192.03], 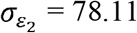, 95%CI [76.78, 79.47]; replication experiment: *µ*_1_ = 740.44, 95% CI [710.10, 771.76], 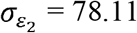, 95%CI [135.79, 144.74], *µ* _2_ = 1187.46, 95% CI [1186.25, 1188.68], 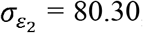, 95% CI [79.46, 81.15]). As the cue onset was fixed in Study 2 at 850ms, the mean cue-dependent RTs of approximately 1190ms corresponded with an effective RTs center of approximately 340ms in both experiments. Anticipatory RT mean varied considerably between participants, as identified by the 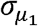 parameter, which modeled between-participant variability in 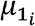 parameter estimates (preliminary experiment: 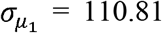, 95%CI [73.29, 166.69]; replication experiment: 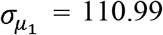, 95%CI [90.92, 136.34]).

Transition from cue-dependent to anticipatory responses, as captured by 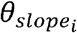 parameter estimates of the preliminary experiment, indicated faster transition in the 100% contingency condition, compared with the 50% contingency condition (*θ* _*slope100%*_, 95%CI [1.03, 2.75], *θ*_*slope50%*_;*=0*.*90*, 95%CI [-0.01, 1.68]; mean difference in *θ*_*slope*_, 95%CI = [-0.17, 2.23]), with prominent variability between participants (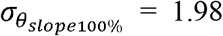, 95%CI [1.45, 2.67], 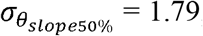, 95%CI [1.21, 2.57]). This effect was replicated even more distinctly in the larger replication experiment (*θ* _*slope100%*_=3.25, 95%CI [2.54, 3.89], *θ*_*slope50%*_=1.54, 95%CI [1.11, 1.92]; mean difference in *θ* _*slope*_, 95%CI = [0.90, 2.47]; 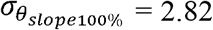, 95%CI [2.37, 3.42], 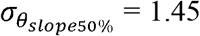, 95%CI [1.16, 1.82]). The 95% credible interval of the difference between the two conditions 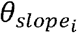 parameters indicated a general trend in the preliminary experiment which was distinct in the replication experiment.

#### Preference modification in probe task

Based on the results of the meta-analysis in Study 1, we hypothesized that manipulating the Go cue contingency would correspondingly manipulate the preference modification effect, manifested as more pronounced preference modification for stimuli in the 100% contingency condition, compared with the 50% contingency condition.

As expected, in the preliminary experiment, participants chose Go stimuli over NoGo stimuli above chance level, both in the 50% contingency condition (prop. = 57.11%, *Z* = 2.85, *p* = 0.002, odds = 1.33, 95% CI [1.09, 1.62]; one-sided logistic mixed model), as well as in the 100% contingency condition (prop. = 64.45%, *Z* = 3.40, *p* = 3.4E^-4^, odds = 1.81, 95% CI [1.29, 2.56]). The preference modification effect was more robust in the 100% contingency condition, compared with the 50% contingency condition (*Z* = 1.92, *p* = 0.028, OR = 1.36, 95% CI = [0.99, 1.87]). These effects were prominently replicated in the larger replication experiment (50% contingency condition: prop. = 58.41%, *Z* = 5.50, *p* = 1.9E^-8^, odds = 1.40, 95% CI [1.24, 1.59]; 100% contingency condition: prop. = 66.31%, *Z* = 6.41, *p* = 7.1E^-11^, odds = 1.97, 95% CI [1.60, 2.42]; condition difference-effect: *Z* = 3.81, *p* = 7.0E^-5^, odds = 1.40, 95% CI [1.18, 1.67]; one-sided logistic mixed model), see Figure 6.

**Figure 6.**
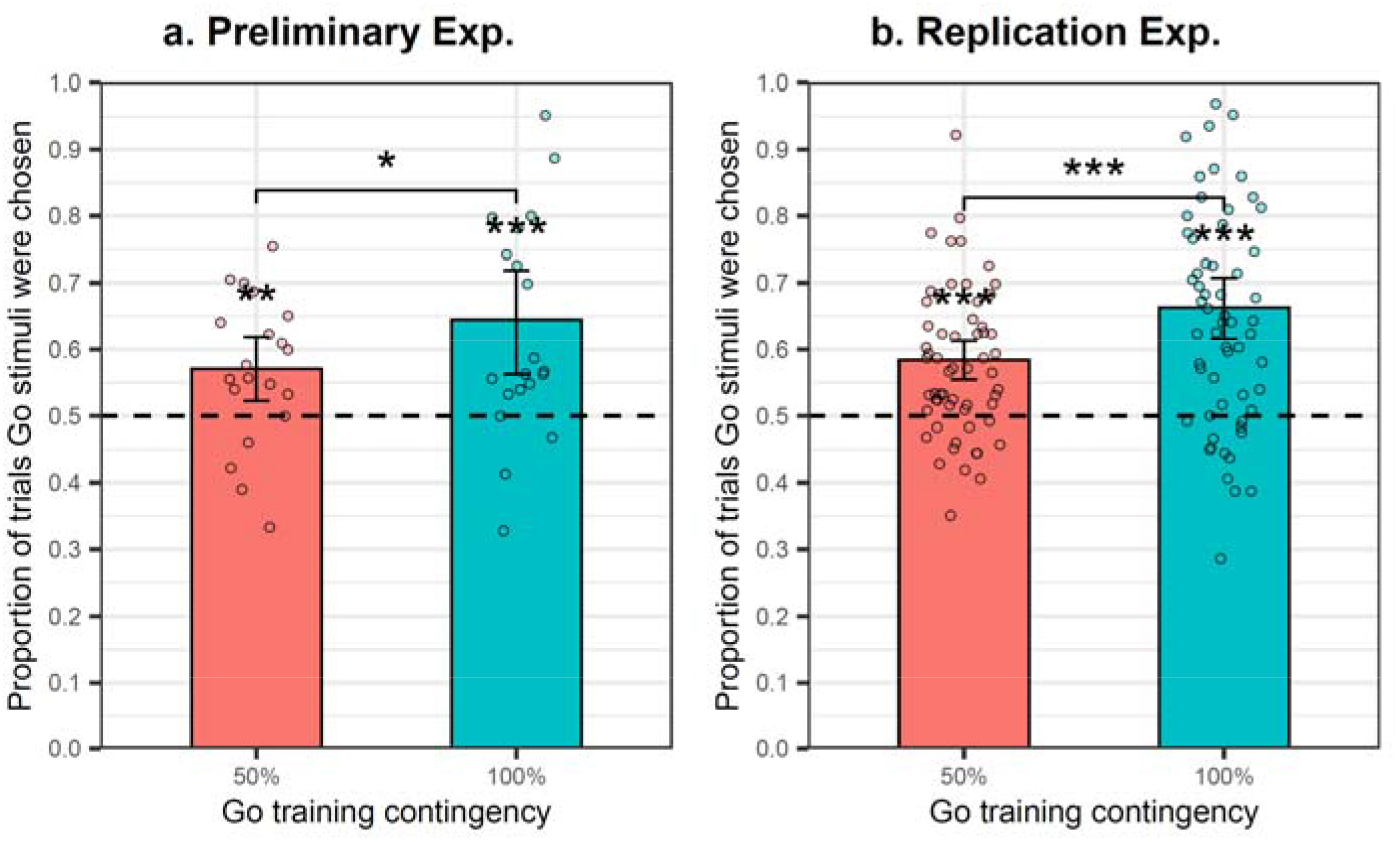
Probe results. Proportion of trials participants chose Go stimuli over NoGo stimuli of similar initial value, in the preliminary experiment (a) and replication experiment (b). Dots represent individual participants, error-bars represents 95% CI based on a mixed model logistic regression. Participants demonstrated enhance preference both for Go stimuli in the 50% contingency condition and to a larger extent in the 100% condition. Statistical significance is denoted with asterisks (* *p* < 0.05, ** *p* < 0.01, *** *p* < 0.001; one-sided mixed model logistic regression). Dashed line represents 50% chance level.

We also examined the impact of initial subjective value on the reported effects by controlling for the initial value difference within each probe choice, as well as by examining the interaction of contingency effect with value categories. In all of the analyses, our effects of interest remained consisted, i.e., we observed overall enhanced preferences for Go over NoGo stimuli, and a more robust preference for Go stimuli within the 100% contingency condition (see supplementary materials).

#### Prediction of probe using computational model

Most importantly, we aimed to examine whether the preference modification effect could be predicted using the individually fitted 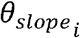 parameter estimated for each participant. Adding the individual learning parameter as an additional independent variable to the logistic regression model revealed that indeed 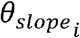 parameter estimated from the training task, could be used to predict preference modification in the subsequent probe task (Figure 7). In the preliminary experiment a significant contribution was found for 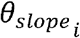 in the 50% contingency condition (log-OR = 0.12, *Z* = 2.30, *p* = 0.011, OR = 1.13, 95% CI [1.02, 1.26]; one-sided mixed model logistic regression), as well as in the 100% contingency condition (log-OR = 0.26, *Z* = 3.59, *p* = 1.7E^-4^, OR = 1.29, 95% CI [1.12, 1.49]). No significant difference was found between the slopes under the two conditions (log-OR = 0.13, *Z* = 1.52, *p* = 0.13, OR = 1.14, 95% CI [0.96, 1.36]; two-sided mixed model logistic regression). These results were replicated also in the larger replication experiment, where 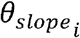 parameter estimates predicted the preference modification effect, both in the 50% contingency condition choices (log-OR = 0.10, *Z* = 2.39, *p* = 0.008, OR = 1.11, 95% CI [1.02, 1.20]; one-sided mixed model logistic regression), as well as in the 100% contingency condition (log-OR = 0.11, *Z* = 3.49, *p* = 2.4E^-4^, OR = 1.12, 95% CI [1.05, 1.19]). No differential slope effect was found between the two conditions (log-OR = 0.01, *Z* = 0.26, *p* = 0.79, OR = 1.01, 95% CI [0.93, 1.10]; two-sided mixed model logistic regression).

**Figure 7.**
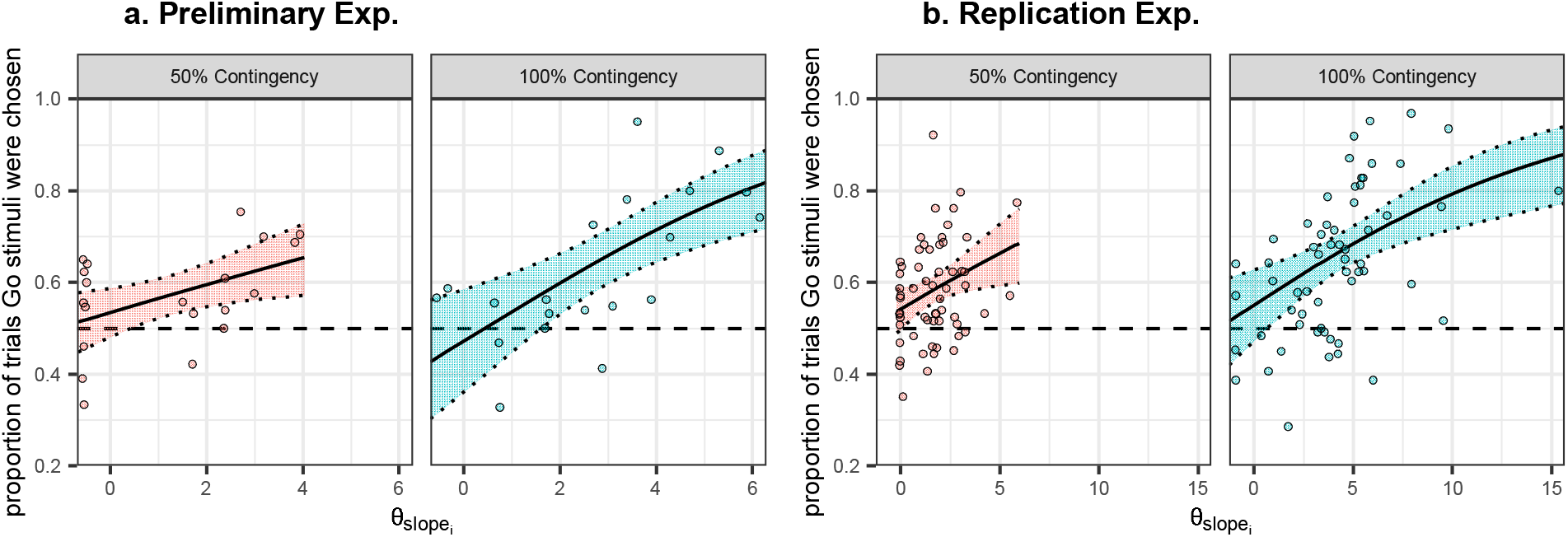
Association of probe choices with training computational marker in study 2. Larger 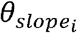 estimates parameter estimates were positively associated with subsequent preference modification in probe phase, both in the preliminary experiment (a) and in the larger replication experiment (b). The 100% contingency condition was characterized with larger 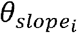 values, and respectively also stronger preference modification effect. Trend line and surrounding color margins represent estimated preference modification effect and 95% CI, respectively (mixed model logistic regression). Dots represent individual participants. Horizontal dashed line represents 50% chance level.

Examining the sizes of estimated fixed-effect, random-effect and residual variances, revealed that the model accounted for a large portion of the variance in participants’ choice pattern in both experiments, both when examining the contribution of the fixed effects only 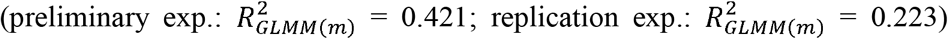, and furthermore when examining the joint contribution of fixed and random effects 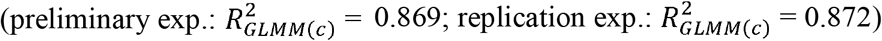.

To examine the overall effect of contingency condition above and beyond 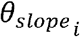, using a likelihood ratio test, the logistic regression model was compared with a restricted (nested) model that did not contain contingency fixed effects (contingency intercept and slope interaction; restriction of 2 parameters). In both experiments, comparing the two models revealed no significant contribution of contingency above and beyond 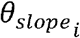 (preliminary experiment: Δ*AIC*_(*rest*.*-full*)_ = -1.64, 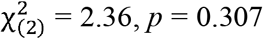 replication experiment: Δ*AIC*_(*rest*.*-full*)_ = -3.61, 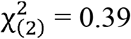, *p* = 0.824; two-sided likelihood ratio test).

#### Post-hoc model validation

Like in Study 1, in a post-hoc analysis we fitted a non-Gaussian model (log-normal distributions) and found similar fit to actual RT data (see Supplementary Fig. 16), as well as positive association of the 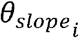 parameter estimates with subsequent probe choice both in preliminary experiment (50% condition: log-OR = 0.03, *Z* = 2.45, *p* = 0.024, OR = 1.03, 95% CI [1.02, 1.04]; 100% condition: log-OR = 0.04, *Z* = 3.75, *p* = 0.001, OR = 1.04, 95% CI [1.03, 1.51]; one-sided mixed model logistic regression), and in the replication study in 50% condition (50% condition: log-OR = 0.014, *Z* = 2.12, *p* = 0.038, OR = 1.01, 95% CI [1.008, 1.02]; one-sided mixed model logistic regression). We also performed a post-hoc analysis in which we compared the explanatory power of 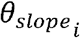 with a simpler marker using the proportion of anticipatory responses each participant made (i.e. the proportion of RTs which were faster than the top 1% of RTs in the first run; see Supplementary). In contrast to the previous meta-analysis results, in the two Study 2 experiments, we found that 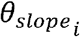 had a significant predictive power, above and beyond a simpler marker which is based on the proportion of anticipatory responses participants made during training.

Examining the interplay with choice RT. Finally, to examine the hypothesis that 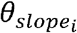 impacts choices via faster automated response to Go stimuli, we introduced two new exploratory analysis which examined whether 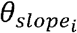 correlated with faster RT during probe. We found no evidence supporting that probe choices were associated with choice RT during probe (preliminary experiment - 50% contingency choices: *b* = 4.56, *t*(18) = 0.18, *p* = 0.859, 100% contingency: *b* = 9.27, *t*(18) = 0.42, *p* = 0.682; replication experiment - 50% contingency choices: *b* = 1.18, *t*(57) = 0.09, *p* = 0.928, 100% contingency: *b* = 3.79, *t*(57) = 0.57, *p* = 0.568; two-sided linear mixed model).

We further examined whether the predictive power of 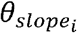 remained consistent above and beyond the time it took participants to make their probe choices. We introduced to the logistic regression analysis used to examine the predictive power of 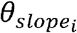 an additional regressor of the choice RT (in seconds). We found that choice RT often had a significant explanatory power, wherein faster choices were associated with higher likelihood of choosing to Go stimuli (preliminary experiment – 50% contingency: log-OR = 0.35, *Z* = 1.31, *p* = 0.19, OR = 1.42, 95% CI [0.84, 2.40], 100% contingency: log-OR = -0.72, *Z* = -2.33, *p* = 0.019, OR = 0.49, 95% CI [0.26, 0.89]; replication experiment – 50% contingency: log-OR = -0.58, *Z* = -3.50, *p* = 4.6E^-4^, OR = 0.56, 95% CI [0.41, 0.78], 100% contingency: log-OR = -1.01, *Z* = -5.59, *p* = 2.2^-8^, OR = 0.36, 95% CI [0.26, 0.52]; two-sided mixed logistic regression). Nonetheless 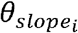 was consistently found to be predictive of choosing Go stimuli, above and beyond choice RT (preliminary experiment – 50% contingency: log-OR = 0.12, *Z* = 2.44, *p* = 0.007, OR = 1.13, 95% CI [1.02, 1.25], 100% contingency: log-OR = 0.26, *Z* = 3.36, *p* = 3.8^-4^, OR = 1.30, 95% CI [1.12, 1.52]; replication experiment – 50% contingency: log-OR = 0.1, *Z* = 2.42, *p* = 0.008, OR = 1.11, 95% CI [1.02, 1.21], 100% contingency: log-OR = 0.12, *Z* = 3.53, *p* = 2.0^-4^, OR = 1.12, 95% CI [1.05, 1.20]).

#### Stimulus-specific learning parameter

In an additional analysis (mentioned but not detailed in the preregistration), we extended this analysis to stimulus-level effects. Each Go stimulus_s_ presented to participant_i_ was fitted a 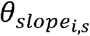 parameter, which was expected to capture within-participant variability in learning - i.e., model for difference in learning of stimuli within a contingency condition. We aimed to use these participant and stimulus-specific learning parameters as independent variables in a mixed model logistic regression (with a random intercept, random slope for contingency and random slope for 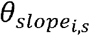 independent variables, see methods for full description of the statistical models).

Fitting a full Stan model for both stimuli of 50% and 100% contingency simultaneously did not converge. We hypothesized that the full model, with 2 conditions × 16 stimuli × n participants inter-dependent 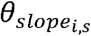 parameters was too complex to resolve using MCMC process. Thus, we decided to reduce complexity by eliminating the participant-level effects on the mean RT for the early anticipatory responses (estimating only group-level *µ*_1_, without participant-level 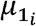 parameters; as was done in the previous meta-analysis study). We further split the data based on contingency condition and analyzing each contingency as an independent dataset. Using this approach resolved the convergence issues. This solution for an unexpected issue was not preregistered or planned prior to data analysis.

Examining the association between the 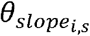 parameter estimates and the participant-level 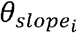 parameter estimates of the previous model, revealed high similarity (preliminary exp.: *r* = 0.84; replication exp.: *r* = 0.80). In addition, participants with negative 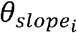 parameter estimates had very small variance (SD < 0.1) in their 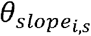 estimates (see Supplementary Fig. S8). When such 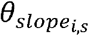 were introduced in the mixed-model logistic regression, the low within-participant variability caused convergence warnings. Therefore, all participants with 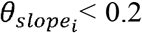 (which is equivalent to SD < 0.1 of 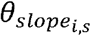) in either contingency condition, were excluded from the analysis. This resulted in the exclusion of nine participants in the preliminary experiment and 14 participants in the replication experiment.

Analyzing the remaining 11 and 45 participants in the preliminary and replication experiment, resulted in similar conclusions, as found with the participant-level 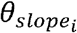 parameter estimate. Overall, in both contingency conditions, a significant positive association was found between 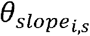 and preference of Go stimuli over NoGo stimuli (preliminary experiment - 50% contingency: *Z* = 4.34, *p* = 7.0E^-6^, OR = 1.33, 95% CI [1.17, 1.52]; 100% contingency: *Z* = 2.85, *p* = 0.002, OR = 1.21, 95% CI [1.06, 1.38], overall model 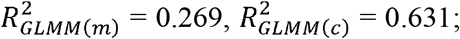; replication experiment - 50% contingency: *Z* = 3.14, *p* = 8.4E^-4^, OR = 1.09, 95% CI [1.03, 1.15]; 100% contingency: *Z* = 5.09, *p* = 1.7E^-7^, OR = 1.09, 95% CI [1.05, 1.12], overall model 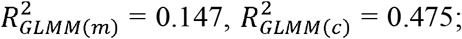; one-sided mixed model logistic regression; Figure 8). No significant differences were found between the slopes of the two conditions (preliminary experiment: *Z* = -1.35, *p* = 0.18, OR = 0.91, 95% CI [0.79, 1.04]; replication experiment: *Z* = - 0.08, *p* = 0.94, OR = 1.00, 95% CI [0.95, 1.05]; two-sided mixed model logistic regression).

**Figure 8.**
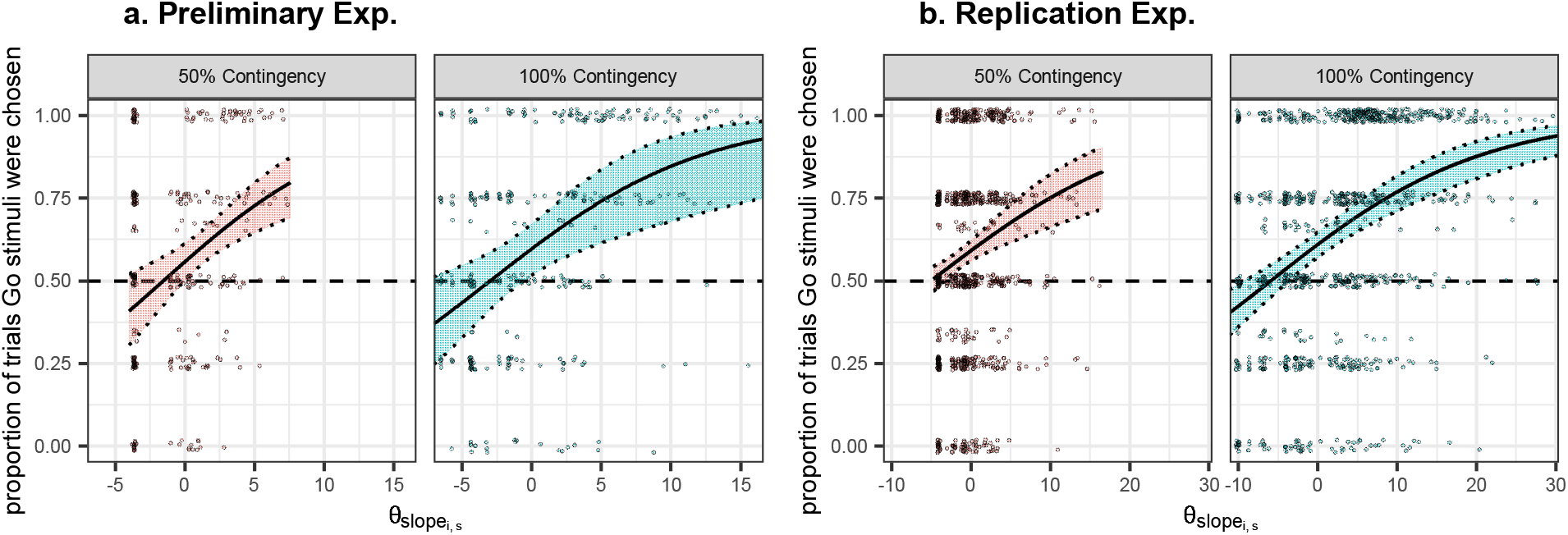
Association of probe choices with stimulus-level computational marker in study 2. Larger 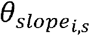 estimates parameter estimates were positively associated with subsequent preference modification in probe phase, both in the preliminary experiment (a) and in the larger replication experiment (b). Dots represent individual stimuli (small vertical jitter was added to better visualize high density areas). Trend lines and surrounding color margins represent estimated preference modification effect and 95% CI, respectively (mixed model logistic regression). Horizontal dashed line represents 50% chance level.

The contribution of stimulus-level 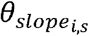 parameter to predicting preferences was examined by comparing a full model, containing both stimulus-level 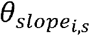 and participant-level 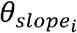 individualized learning parameters to a restricted model containing only 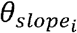 as independent variable. The addition of 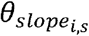 had a significant contribution both in the preliminary experiment (ΔAIC = 59.65, 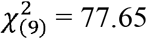, *p* = 4.7E^-13^; likelihood ratio test), as well as in the replication experiment (ΔAIC = 93.97, 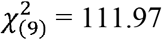, *p* = 5.8E^-20^).

### Interim discussion

Based on the conclusions of the meta-analysis study, a novel preference modification procedure was designed and tested in Study 2. In the new design, two cue contingency conditions (50% and 100% contingency) were tested within-participant, which we hypothesize would manipulate the difficulty of learning the stimulus-cue association during the training phase of the CAT paradigm. The introduction of the two contingency conditions allowed us to experimentally impact the 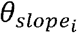 individualized computational marker of learning, which in turn provided a reliable predictor for the subsequent preference modification effect in the probe phase. As expected, we found a more robust preference modification effect for Go stimuli in the 100% contingency condition, compared with the 50% contingency condition. The fact that a directed experimental intervention in the preceding training phase induced a differential effect in the probe phase, provides support for a directional causal impact of training on preference change. In light of these results, it is plausible to conclude that the association demonstrated in Study 1 between the individualized learning parameter and behavioral change might represents a causal relationship, in which improved learning in the training phase induces stronger behavioral change in the subsequent probe phase.

Furthermore, between-participant variability in the 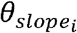 computational marker was also predictive of between-participant difference in preference modification, as measured during the subsequent probe phase. Participants with more robust 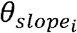 also demonstrated stronger preference for Go stimuli over NoGo stimuli of similar initial value. This effect was observed both in the 100% and 50% contingency conditions, with no significant difference in the effect between the two conditions. This suggests that 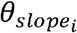 had a similar predictive pattern for both conditions, one unit increase in 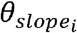 resulted in similar increase in the odds of choosing Go over NoGo, above and beyond the contingency condition. However, as the 100% contingency condition was characterized with larger 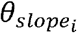 parameter estimates, so was the probe phase characterized with greater odds of choosing Go stimuli (i.e., stronger preference modification effect). Taken together, these results provide evidence for novel means to both predict individual differences in learning and preference modification between participants, as well as means to manipulate this preference modification effect. All the non-exploratory results in the preliminary experiment were carefully detailed in a preregistration, and were all replicated in a larger replication experiment, which included the exact same procedure and analysis pipelines. It is important to note that using a larger sample size not only replicated the significance of the effects, but also the descriptive trends, resulting in outcomes of similar effect size.

In contrast to post-hoc analysis in Study 1, which found that the computational marker performed similarly well as a simple RT-based marker, in a post-hoc analysis of the two new experiments, we did find that 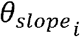 captured a unique predictive signal above and beyond a simple proportion of anticipatory responses. These results reinforce our hypothesis, that by optimizing the training procedure for our computational model (by fixing the Go cue onset time and explicitly encouraging participant to make anticipatory responses), we were able to better capture learning signals using 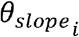 marker.

It is interesting to note that descriptively the new design induced a strong preference modification effect in the 100% Go contingency (group estimate of approximately 65% in both experiments), compared with other CAT and Go/NoGo experiments, were a more modest effect of 55%-60% preference bias for Go stimuli was commonly observed ^12–20^. Without carefully controlling for all factors, it is hard to conclude whether this difference is significant, and which one of the changes in the experimental design induced this enhanced effect, however, it would be interesting to try and address this directly in future work.

In an exploratory analysis, we examined variation across stimuli as well as participants, focusing both on the variability in learning patterns between different participants, as well as within-participant variability in learning for different stimuli. Due to the technical requirements, such an in-depth analysis was applicable only for a smaller subset of our data, however, the partial results we found suggest that the prediction model is applicable in all levels of granularity, both at the individual participant level, as well as in the individual stimulus level.

### General discussion

In the current work, we aimed to study and characterize the cognitive mechanisms underlying non-reinforced preference modification with CAT^12^, focusing on the training phase where individual items are presented. While behavioral change is manifested in a binary probe phase, actual preference change occurs at the individual item level. Based on previous studies that alluded to the potential involvement of internal non-reinforced learning in CAT, the current work aimed to use computational modeling to measure this internal process at the training phase with individual items and establish its causal role on the subsequent preference modification effect. The CAT task is a multiphase standardized procedure, which includes an initial preference evaluation task, a non-reinforced training phase on individual items, and a binary probe phase. Therefore, the task provided a unique opportunity to aggregate data from multiple studies and build a large corpus of training data.

In the first part of the current work, we examined previously collected data of n=864 participants from 29 different CAT experiments in a meta-analysis and identified a distinct time-dependent RT pattern during the training task. We found that while in early stages of training, participants depended on the Go cue to initialize responses (homogeneous RTs following cue onset), as training progressed, participants relied less on the Go cue and started to generate fast anticipatory responses. This raised the hypothesis that a marker of transition from cue-dependent responses to early anticipatory responses could be indicative of internal non-reinforced learning and subsequent preference changes.

Using a Bayesian computational framework implemented with Stan, a statistical model of RT patterns was formulated. RTs were postulated to derive from a mixture model of two Gaussian distributions and a time-dependent mixture proportion, which accounted for the transition speed from late cue-dependent responses to early anticipatory responses. The model parameters were estimated with MCMC algorithm, with a key participant-level 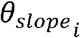 parameter, which modeled the individualized difference between participants in transition to anticipatory responses rate. This Bayesian parameter was used as a passive computational marker of learning.

Examining the association between the individualized learning parameter and preference modification effect in the subsequent probe task, showed a positive association in which participants with more robust learning parameter estimates during training also demonstrated stronger preference modification effect in the subsequent probe phase (i.e., enhanced preference for the Go stimuli over NoGo stimuli of similar initial value). Thus the 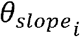 parameter, which was optimize to model RT pattern during training, was found to be good predictor of preference modification effect in a future probe task. An additional analysis modeling RT using a more finely tuned learning 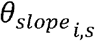 parameter, fitted for the individual stimuli, further expended these findings, and demonstrated positive predictive power of 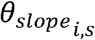 in evaluating preference modification effect on a stimulus-level basis, above and beyond the more general participant-level parameter.

Based on the results of Study 1, we derived a theoretical hypothesis with implication for shaping learning efficacy and consequently also the effect of training on preference behavior. If faster transition to anticipatory responses is related to learning, then hindering the ease of cue anticipation could also obstruct learning and resulting in less pronounces preference modification effect. To examine this hypothesis, we design a novel CAT procedure, which was tested in a preliminary experiment and an additional larger preregistered replication experiment. Cue anticipation was manipulated by introducing two training conditions within the CAT task - in the 100%-contingency condition, the Go stimuli perfectly anticipated Go cue onset, while in the 50% contingency condition, the Go stimuli only anticipated the Go cue in half of the trials.

Manipulating cue-contingency during CAT induced the expected effects both in the training and in the probe task. During training, participant relied less on early anticipatory responses and transitioned slower to anticipatory responses in the 50% contingency condition. This effect was captured by the 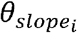 estimates, which were larger in the 100% contingency condition compared with the 50% contingency condition. More importantly, in the subsequent probe phase, a consistent pattern of stronger preference modification effect was observed for the 100% contingency Go stimuli. Thus, manipulating training difficulty has also affected future preference modification measured in the probe phase. When individual differences in preference modification were examined, 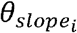 marker accounted for a large portion of variability in choices, while no significant difference was found between the two contingency conditions above and beyond the 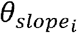 effect. This suggest that the more robust preference modification effect observed for the 100% contingency Go stimuli, was captured by the stark differences in 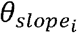 learning marker. Furthermore, the fact that intentionally manipulating the training procedure in Study 2 induced differential learning parameter estimates as well as differential behavioral change effect, further supports the hypothesis of a causal nature of the association with the behavioral change. Thus, it is plausible to deduce that shaping training at the individual item level, which was captured by the individualized learning parameter, has driven the induced differential behavior modification effect, as evaluated in the subsequent probe phase. It is important to note that although the cue-contingency was different between the two conditions, the actual exposure-time to the stimuli was identical. Thus, the enhanced preference for Go stimuli in the 100% contingency condition could not be accounted for by mere exposure effect ^6^.

### What is the mechanism?

These findings, showing a consistent link between motor-planning during training at the individual items level and preference modification in the subsequent binary choice probe phase, correspond with previous works. These works showed that a rapid response is a crucial feature for preference modification with CAT^13^, and neural finding showing that increased striatal and premotor activity during training, were associated with more robust preference modification effect in the probe phase^18^. Thus, the results suggest that learning efficacy in the training phase is manifested as stimulus-specific motor planning. Importantly, all participants were alert and attentive throughout the training phase, as demonstrated by the negligible rates of non-response. However, it evident that mere attention is not sufficient to induce strong preference modification, but rather an attention that can be translated to action, suggesting a unique valuation pathway in the absence of external reinforcements, putatively based on parieto-frontal circuits involving attention with motor planning ^5^.

To produce an anticipatory response for a Go stimulus, participants must also rely on memory. Thus, it is possible that the anticipatory response pattern captured in our data identified individualized difference in memory which in turn mediated a change in preferences^23^. This hypothesis is in line with previous findings with non-reinforced training, which identified a positive association between preferences modification and enhanced memory both at the participant-level ^15^ and at the stimulus-level^19,21^. Since memory for the trained Go stimuli was not examined in the current work, it would be interesting to examine in future experiments whether 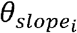 computational marker correlates with declarative memory and does it provide independent predictive power of subsequent non-reinforced preference modification above and beyond memory measurements. Future imaging studies could also examine the new task design with fMRI to identify whether differential neural activation patterns characterize the two contingency conditions.

One question that is often raised in non-reinforced preference modification research is whether the behavioral effect of choosing Go stimuli reflects an internal change in the value of the chosen Go stimuli, or rather habitual automated responses to choose the Go stimuli originating from mechanisms similar to operant conditioning. On the one hand, CAT experiments showed that not all cue-response associations result in enhanced preference for Go stimuli – CAT was found to be less effective in enhancing preferences for stimuli of negative affective valence ^15^ and low-value snack food items ^12^, suggesting that the stimulus pretraining value interacts with the effect of CAT on preferences. In addition non-challenging CAT designs such as CAT where the cue starts with the stimulus onset or when the response association was made as a block of stimuli to which participants were required to respond also failed to induce preference change^13^, which provide evidence that the mechanism impacting preferences requires more than a simple motor response association. On the other hand, some findings elude to the involvement of rapid impulsive decision making mechanisms - CAT experiments in which participants were asked to make slow well considered decisions found that CAT effect on preferences diminished when participants made slow non-impulsive choices. This result further resonates the finding that during the CAT probe phase, participants were faster in their choices when Go stimuli were chosen ^17,41^.

To test these two competing notions that CAT non-reinforced preference modification is driven by impulsive motor response habitual mechanisms versus internal value representation change, we examined in our data whether choice RT could account for the enhanced likelihood of choosing Go over NoGo stimuli during probe. Our results showed most of the times that indeed faster choice RTs were associated with increased likelihood of choosing Go stimuli, in agreement with past findings. However, and more importantly, we found that the computational marker was not correlated with choice RT, and maintained its predictive power of choice above and beyond choice RT. Thus, our results indicate that while some aspects of non-reinforced preference change could be attributed to rapid impulsive choices, our proposed computational marker tracks an independent preference modification mechanism, which could putatively be indicative of value representation change. Future work could attempt to find behavioral or neural correlates which could corroborate this hypothesis.

An explanation offered by the model for the modification of preferences revolves around the presence of an internal reinforcement process during the training phase. Following the initial exposure to stimulus-cue contingencies, participants attempted to respond based on the stimuli rather than the cue itself, displaying anticipatory responses. When participants correctly pressed earlier, an internal feedback mechanism was activated, further enhancing the learning process. This internal reinforcement can be seen as a broader mechanism for facilitating learning. It is challenging to differentiate this internal reinforcer from memory or attention processes, and our study does not aim to do so. Instead, the current model integrates these mechanisms and incorporates the internal reinforcer as an integral component of the learning process. Drawing on previous research and considering the correlation between brain regions associated with the reward system ^18^, we propose that this inner mechanism plays a role in reinforcement. However, since no external feedback was provided during the training task, our hypothesis suggests that the reinforcement mechanism is likely to be internal in nature. The current work’s unique contribution establishing a new passive marker for non-reinforced learning could be tested in future neuroimaging studies. These studies could shade light on whether the marker is associated more with parietal attention mechanisms, temporal memory-related regions or striatal reward and motor-learning neural mechanisms (ref).

In addition to a basic understanding of the cognitive mechanisms of non-reinforced learning, identifying a learning marker based on early training runs could be used to design powerful prediction tool for learning efficacy, with relevance to other paradigms and situations that share the basic training/transfer structure of the current studies. A naïve approach to evaluate learning-efficacy during training tasks such as CAT might suggest to introduce direct measurements of behavior-change throughout the training process, e.g. by asking participants to make active value-based choices ^42,43^. However, it has been shown that the mere act of choice could induce a longitudinal effect of enhanced preference for the chosen stimuli ^8,9,44^ and introduced bias in the effect of preference modification following non-reinforced training ^21^. Therefore, probing preference within a preference modification training, is likely to alter the learning process and undermine its validity. Moreover, in more standard learning procedures, such as conditioning-based interventions, to examine the efficacy of learning researchers observe the subject’s response to the conditioned stimulus when the associated unconditioned stimulus is omitted ^45–52^. Over repetition of probe tests in all of these procedures might initiate a new learning procedure – where the conditioned stimulus is no longer associated with the unconditioned stimulus, which would eventually lead to extinction-learning ^53–56^. Thus, evaluating learning by active probing of the behavior-change effect could interfere with the learning process. However, a passive marker for learning which can predict behavioral change efficacy based on earlier training data, does not evoke the drawbacks of introducing an active choice task to reveal preferences. Using a passive learning marker which is evaluated based on independent training data that preceded the probe phase, overcomes this obstacle. The temporal primacy 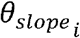 estimation could provide a prediction tool for future preference modification in CAT, without tainting the results with a direct evaluating of preferences using a choice task. An interesting hypothesis could anticipate that the 50% cue-contingency would resemble partial reinforcement learning schedule, and thus will have potentially higher long term sustainability and resistance to extinction ^57^. While the current work did not examine long term maintenance of CAT effect and its association cue-contingency, this could be an interesting subject to test in future studies.

The benefits of real-time individualized markers for learning could also be of great value for additional learning and behavior change procedures beyond CAT. In many learning tasks (including CAT), the behavioral change effect on choices is measured in a separate probe phase, proceeding the training phase. For example, in experiments testing pain perception, participants are trained to associate neutral stimuli with differential level of painful heat stimulation before probing the impact of interventions such as drug versus placebo administration ^58,59^. Studies examining habit formation may use lengthy free-operant learning protocol, in which participants repeatedly perform an action (such as pressing a button) to gain food rewards, in order to test in a later probe phase whether the participants demonstrate habitual behavior and continue to perform the action even when the reward is devalued (e.g., pressing to get food when satiated) ^60–62^. And even outside the field of value-based decision making, in the clinical psychological wellbeing domain, attention bias modification (ABM) procedures use computerized attention training interventions similar in nature to CAT to treat depression and anxiety ^63–65^. In such experimental settings, where training is either unpleasant or exhausting, with plausible odds that learning would not be well-established if under-trained, monitoring learning efficacy individually based on the training data could assist in optimizing training efficacy and efficiency. While the current work focuses on CAT, we assert that the basic mechanisms identified by the current work, could be adapted to accommodate a need in a wider range of learning procedures. With an appropriate adaptation to the desired experimental design, this unique feature of a passive computational marker which is predictive of subsequent change, opens a new path for potential interventions that will allow more efficient training via monitoring and real-time feedback ^66,67^.

The current work’s approach in modeling learning using RT could also be applicable in experimental designs which do not separate the probe from the training task, such as reinforcement learning tasks, where learning can be evaluated as it progresses on a trial by trial basis ^68–71^. Since RT patterns could be indicative of the individual’s confidence in her choice during reinforcement learning tasks such as probabilistic selection ^51,52^ or multi-arm bandit task ^72,73^, incorporating RT data could improve reinforcement learning models and capture learning more accurately ^74^.

Some limitations were not directly tested in the current work and should be addressed in future studies. Primarily, in Study 2 of the current work, we demonstrated that learning and preference modification could be hindered using a more difficult/partial association procedure. However, our model also predicts that easing the association procedure could enhance learning efficacy and preference modification effect. Further work should empirically test this hypothesis, for example by examining the effect of enhancing the Go cue saliency or attention during predefined Go stimuli ^27,75,76^, or by manipulating the temporal features of training schedule ^77^. Furthermore, by instructing participants to respond when they have enough confidence that a Cue will follow, we might have introduced an undesired confound to the task in the subjectivity of confidence each participant needed to respond; i.e., some participants might have learned well the association between a stimulus and the Go cue but were reluctant to respond before they validate that in a 50% contingency condition a cue will appear. Although such confound does not undermine the validity of the findings, future studies should avoid it and aim to examine a differential effect in which the difference between conditions reflect only task difficulty, for example, by asking to respond to stimuli which at any point were associated with a cue, or by clarifying in the instruction that a response based on guess is also valid.

Another limitation of the current work could raise a concern for a confound which might undermine the ability to deduce causal relationship between learning and choice in our new experimental design. In two experiments, we found that participants demonstrated an enhanced preference modification effect for stimuli that were more consistently associated with Go cue. By maintaining a similar presentation time of both 50% and 100% contingency stimuli, we eliminated the confound of the standard mere exposure effect ^6^, as participant viewed the stimuli in both conditions for the same time duration and putatively had to maintain high alertness to both type of stimuli, which both required response with a high likelihood. However, one might argue that participants were more engaged with the 100% contingency stimuli, to which they pressed twice as much. Thus, an alternative explanation could claim that increased engagement (i.e., more press responses) is the true causal factor which induced a stronger preference modification effect, rather than the learning procedure identified by the computational marker.

Our statistical analysis provides evidence that suggests that the learning marker had a stronger impact on choices than the level of exposure. When analyzing the factors predicting choice patterns, the training condition had no explanatory power above and beyond the computational marker of learning. Nonetheless, a dedicated experimental design could aim to directly discern between these two competing hypotheses. One such proposed training design could present the 50% contingency stimuli twice as often as 100% contingency stimuli. This would result in a training design with more consistent association for the 100% contingency condition (similarly to the current design), while maintaining equal engagement levels between the two conditions (manifested as identical number of press responses in both training conditions) and an increased exposure to the 50% contingency stimuli, which would be presented for twice the duration of time as the 100% contingency condition stimuli. Our theory hypothesizes that despite the increased exposure and similar engagement with the 50% contingency stimuli in this design, participants will still demonstrate faster learning in the consistent 100% contingency condition and would thus show stronger preference modification effect for the 100% contingency condition stimuli. Future work could try to run this dedicated design and provide empirical evidence which would settle whether our proposed model of learning overcome mere exposure effect combined with more equal engagement-level, which we did not fully address in the current work.

We carefully documented the experimental choices in a preregistration before data was collected for an independent replication experiment. Utilizing a Bayesian computational framework, such as the one used here, provided great flexibility required to fit complicated theoretical model. We aimed to use a relatively straightforward model, with as similar features as possible in all studies. Future studies could take the liberty to use the openly accessible data from this work and try to improve the model by changing our current assumptions including distributions’ shape, free parameters, and priors.

In conclusion, the current work laid the foundations for understanding and quantifying non-externally reinforced learning at the individual item level via a novel Bayesian modeling approach of RT pattern during training. Using a large meta-analysis dataset of previous CAT studies and two new original experiments, we demonstrated a unique method to evaluate a learning marker at the individual item level with a robust predictive power of subsequent preference change. Thus, we propose that motor response patterns could provide a passive marker for value-change, which does not require direct measurement of preferences.

## Methods

### Study 1 – CAT meta-analysis

#### Data collection and sample sizes

For study 1 meta-analysis, the data from 29 previous CAT experiment was combined. The dataset included 21 experiments from published works by our research group and collaborators ^12–16,18,19,22^, as well as eight additional unpublished works, which were collected as part of the preparations for previously published manuscripts, or are planned to be published in the future. The experiments comprised of a median sample size of *n* = 26 (*M*_*sample size*_ = 29.79, range = [23, 70]; see Table 1 for information of the sample size in each experiment, and the supplementary data for more detailed description of the unpublished data).

**Table 1.** Experiment included in the meta-analysis.

| Exp. | Stimuli | n | Training runs | Go cue | Publication (exp. number in publication<sup>a</sup>) |
| --- | --- | --- | --- | --- | --- |
| 1 | Fractal art | 25 | 12 | Auditory | Salomon et al., 2018 (2) |
| 2 | IAPS positive | 27 | 12 | Auditory | Salomon et al., 2018 (3) |
| 3 | IAPS negative | 28 | 12 | Auditory | Salomon et al., 2018 (4) |
| 4 | Snacks (IL) | 25 | 20 | Visual | Salomon et al., 2018 (5) |
| 5 | Snacks (IL) | 25 | 20 | Neg. auditory | Salomon et al., 2018 (6) |
| 6 | Faces | 25 | 20 | Auditory | Salomon et al., 2018 (7) |
| 7 | Fractals | 25 | 20 | Auditory | Salomon et al., 2018 (8) |
| 8 | IAPS positive | 29 | 20 | Visual | Salomon et al., 2018 (9) |
| 9 | IAPS negative | 32 | 20 | Visual | Salomon et al., 2018 (10) |
| 10 | Faces: politic. | 25 | 20 | Auditory | Unpublished |
| 11 | Faces: politic. | 39 | 20 | Auditory | Unpublished |
| 12 | Faces | 42 | 16 | Auditory | Salomon et al., 2019 |
| 13 | Faces: affective | 42 | 20 | Auditory | Unpublished |
| 14 | Faces: affective | 70 | 20 | Auditory | Unpublished |
| 15 | Fractals | 29 | 16 | Auditory | Aridan et al., 2019 |
| 16 | Snacks (US) | 26 | 12 | Visual | Unpublished |
| 17 | Snacks (IL) | 24 | 20 | Visual (Reward)<sup>b</sup> | Unpublished |
| 18 | Snacks (IL) | 30 | 16 | Auditory | Botvinik-Nezer, Bakkour, et al., 2021 (1) |
| 19 | Snacks (IL) | 25 | 16 | Auditory | Botvinik-Nezer, Bakkour, et al., 2021 (Pilot) |
| 20 | Snacks (IL) | 23 | 12 | Auditory | Unpublished |
| 21 | Snacks (IL) | 25 | 20 | Auditory | Unpublished |
| 22 | Snacks (US) | 29 | 12 | Auditory | Schonberg et al., 2014 (1) |
| 23 | Snacks (US) | 25 | 8 | Auditory | Schonberg et al., 2014 (2) |
| 24 | Snacks (US) | 25 | 12 | Auditory | Schonberg et al., 2014 (3) |
| 25 | Snacks (US) | 27 | 16 | Auditory | Schonberg et al., 2014 (4) |
| 26 | Snacks (US) | 26 | 16 | Auditory | Schonberg et al., 2014 (7) |
| 27 | Snacks (US) | 30 | 12 | Auditory | Bakkour et al., 2017 |
| 28 | Snacks (US) | 25 | 16 | Auditory | Bakkour et al., 2016 |
| 29 | Snacks (IL) | 36 | 16 | Auditory | Botvinik-Nezer, Salomon, et al., 2019 |

<sup>a</sup> In papers with multiple experiments, we note in parenthesis the experiment number or title as can be found in the related manuscript.
<sup>b</sup> In experiment 16 a visual cue was used indicating to participants they were awarded with a sum of money, which accumulated throughout the training phase.

All participants gave their informed consent to take part in the experiments. In most experiments, participants received monetary compensation for their time, and in few experiments, some participants took part in the experiment in exchange for course credit. All experiments were approved by the ethical review board of the institutes where they were performed (Tel Aviv University, The Hebrew University of Jerusalem, University of Texas at Austin, and McGill University).

#### Stimuli

The different experiments included in Study 1 examined the effect of CAT on preferences for various stimuli (see table 1 for a summary of all experiments). In most of the experiments, the stimuli set comprised of images of familiar local snack food items, popular in the US (eight experiments) or in Israel (eight experiments), which participants received for actual consumption as part of the experiment. Other experiments used face stimuli of unfamiliar figures posing neutral expression from the Siblings dataset ^40^ (two experiments), unfamiliar faces with neutral and happy expression from the Karolinska directed emotional faces dataset ^78^ (two experiments) or familiar faces of famous Israeli politicians (two experiments). Unfamiliar abstract stimuli of fractal art ^79^ were used in three experiments. Two experiments included positive affective stimuli from the international affective picture system ^80,81^ dataset and two experiments included negative affective stimuli from the IAPS dataset.

The face and snack stimuli were modified using Photoshop to remove any background features and create visual standardization within each experiment. In each experiment, all stimuli were colored images of identical dimensions, where the subject of the image (the snack or the face) was centered in the middle of the image frame and the background was replaced with homogenous background (either black or gray). In experiments with fractal art and IAPS stimuli, the images were only cropped to identical pixel dimensions.

##### Go stimuli used in the CAT task

In most experiments (23 out of 29) a neutral auditory cue was used as Go cue during the cue-approach training task. In the remaining experiments a neutral visual cue (Experiments 4, 8, 9, and 16), an aversive auditory cue (Experiment 5), or a visual cue indicating a reward (Experiment 17), was used. Both types of visual cues (neutral and reward related) comprised of a semi-transparent Gabor shape, presented on top of the associated Go stimuli. In experiment 17, the appearance of the visual cue indicated to the participant that she or he had won an additional amount of money, which accumulated across the training task.

#### Procedure

While the different experiments diverged in several aspects of their procedure, all experiments maintained three key phases: an initial-preferences evaluation task, followed by a training task, and a probe task (see Figure 1 for illustration of the procedural design).

##### Initial preferences evaluation task

In the first task of the experimental procedure, participants were exposed to the complete set of stimuli for the first time and were required to indicate their subjective preferences using one of two tasks. To evaluate participants’ preferences in experiments using consumable snack food stimuli, participants performed a Becker-DeGroot-Marschak (BDM) auction procedure ^39^. Participants were allocated with either 3 USD (in experiments conducted in the US) or 10 ILS (in experiments conducted in Israel; approximately equivalent to 3.1 USD) which were used to bid on the snack-food stimuli, presented one by one. Participants were informed that at the end of the experiment, one of the trials will be randomly selected, for which the computer will generate a counter bid. If the participants’ bid was higher than that of the computer, they were required to purchase the snack for the lower price bided by the computer. Participants purchased the snack for actual consumption at the end of the experiment. However, if the computer’s bid was higher, participants got to keep the allocated sum of money. Participant were explicitly instructed that the best strategy for the task is to indicate their true subjective preference. Prior to their participation, participants were asked to fast for at least 3 hours, to make sure they were hungry and incentivized to purchase the food items, according to their subjective preferences.

In experiments with of non-consumable stimuli (such as fractals and faces), which are less appropriate to be evaluated using a monetary scale, preferences were evaluated using a binary ranking procedure. Participants were presented in each trial a random pair of stimuli and were required to choose the stimuli they prefer better. Based on the idea of choice transitivity, binary choices were quantified to produce subjective value ranks using the Colley Matrix ranking procedure ^82^.

Following the BDM or binary choice task, stimuli were ranked-ordered according to subjective preferences. The ranks of the stimuli were used as a basis to form two value-groups - one of high-value stimuli (above-median rank) and a group of low-value stimuli (below median rank). The size of the value groups differed between experiments (having 8, 12 or 16 stimuli per value group). In each of the value-groups, half of the stimuli were allocated to be associate with the Go stimuli and response in the subsequent CAT task (Go stimuli), and the other half of stimuli was allocated to be presented in CAT without Go cue (NoGo stimuli).

Allocation for Go and NoGo stimuli within each value category maintained an equal mean rank for Go and NoGo stimuli -e.g., for a high-value group consisting of eight stimuli, the stimuli ranked 8, 11, 12, and 15 were allocated to be Go stimuli, while stimuli ranked 9, 10, 13, 14 were allocated to be NoGo stimuli, such that both allocations were characterized with a mean rank = 11.5. The Go / NoGo allocation was counterbalanced across participants.

##### Cue Approach Training (CAT)

In the CAT task, stimuli were presented individually on the screen center for a fixed duration of one second (except for Experiment 27, where the duration was extended to 1.2 seconds for compatibility with fMRI scanning protocol). In each experiment, all training stimuli were presented once in each training run, thus the number of training runs indicate the number of stimulus repetitions. A fixed proportion of stimuli per experiment (commonly 30% of stimuli; range 25%-40%) were Go stimuli. When a Go stimulus appeared, a delayed Go cue appeared after a Go signal delay. Participants were asked to respond to the Go cue with a button press as rapidly as possible, before the stimulus offset (one second after the stimulus onset). The Go signal delay was adjusted according to the participant’s performance - a failure to respond before stimulus offset resulted in 50ms shortening of the Go signal delay (thus reducing the task difficulty for the following trial), while a successful response on time resulted in 16.667ms increase of the next Go signal delay (making the task more difficult; 1:3 ratio of signal delay increase to decrease). The Go signal delay commonly started at 750ms, and ranged around 700ms (across experiments, *M* = 691.78, *SD* = 102.77).

The training phase in each experiment, consisted of 8 to 20 training runs (see Table 1). Participants were not informed in advance of the contingency between Go signal and Go stimuli. However, as they were repeatedly exposed to the same stimuli, in the later runs of the task, some of the participants were able to identify the Go stimulus and cue association, thus producing accurate faster responses, sometimes even preceding the Go cue onset. This measurement of RT to Go stimuli trials was used in this current work as our main behavioral measurement for the CAT learning model. The details of the model are further discussed below.

##### Probe

In the final probe phase, preference modification following CAT was evaluated. The probe phase was usually preceded by a ‘filler’ task such as filling up questionnaires or ranking liking for fractal art images. This task generally provided some time for consolidation and a dissociation between the CAT and probe phases. In the probe task, participants were presented with pairs of stimuli of the similar initial subjective value (both high-value stimuli or both low-value). In each pair, one of the stimuli was a Go stimulus and the other was a NoGo stimulus. Preference modification was evaluated as the proportion of trials in which participants chose the Go stimulus over the NoGo stimulus, above and beyond the expected 50% chance level. This measurement was used as the main outcome variable which indicated preference modification following CAT.

#### Analysis

In Study 1, we aimed to identify a marker for learning in the CAT task. To achieve this goal, we examined RT patterns in the task. In an exploratory analysis of the meta-analysis data, we identified that as training progressed, mean RT in the task was reduce. We hypothesized that as the training task progresses, participants that were able to identify and learn the stimulus-cue contingency pattern, and thus could generate faster Go responses which do not rely on the delayed Go cue onset, but rather on the Go stimulus onset. We formally modeled this process in a single experiment before testing it on the entire meta-analysis dataset.

##### Computational model of CAT

To model our proposed cognitive mechanism we utilized a Bayesian modeling approach of the RT in the CAT task, using the R implementation of Stan programing language ^83^. RTs were modeled as a mixture of two gaussian distributions (Equation 1) – one Gaussian of shorter mean RT, representing the early anticipatory responses generated when the stimulus-cue association is predicted by the participants; and a second Gaussian with a later mean RT following the cue-onset (i.e., a distribution representing standard cue-dependent responses).

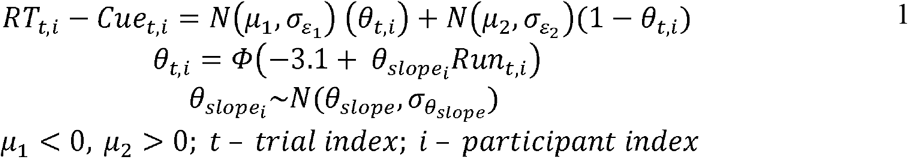

The *θ* _*i,t*_ mixture probability of the two Gaussian distribution was used to determine the proportion of trials participants were expected to produce early anticipatory response (1). It was defined using a linear function of time-dependent 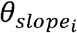 parameter, which was multiplied by the scaled training run index (training repetition), scaled to 0-1, where 0 and 1 indicating the first and last (20th) training repetition, respectively. This 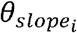 parameter was fitted individually for every participant and was therefore defined conceptually as the individualized learning parameter of interest. To scale the linear function to a range suitable for proportions (0 – 1), we used the normal CDF as a link function.

The baseline proportion of early anticipatory responses (at the first training run) was fixed at the value *θ*_*0*_ = -3.1, corresponding with probability of 0.1% to generate an anticipatory response at the very first run. Thus, the individualized learning parameter could be interpreted as the sole parameter effecting the mixture proportion for the two Gaussian distributions, e.g., 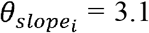 would be interpreted as a *ϕ* (0) = 0.5, 50% proportion of anticipatory responses at the final (20^th^) training run. The individually fitted 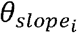 parameter was determined in the current work as the computational marker of learning and was our main parameter of interest. See Supplementary Code S1 for a complete specification of the model parameters and priors.

The model’s parameters were evaluated using Markov-chain Monte Carlo (MCMC) gradient algorithm, implemented with RStan ^83^. Each model was evaluated four independent times, with chains length of 2000 (1000 chain links burn-out time). The mean values of each parameter of the converged model were used as the parameter estimates and are reported along with 95% credible interval (CI). All reported results converged onto stable solution with 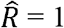.

To maintain stable interpretation of μ_1_ and μ_2_ as the centers of anticipatory and cue-dependent responses, respectively, an upper threshold of 150ms from cue onset was imposed on the μ_1_ parameter, and a lower threshold of 200ms was for *μ* _2_ parameter. Thus, in every chain μ_1_ was imposed the role of the earlier anticipatory responses. Alternative formulation forms of the computational model were also considered and tested, including using 0ms as both upper and lower threshold for the distribution means, using a log-normal distribution instead of a normal distribution, and fitting a unique 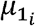 parameter for each participant. However, these models either did not converge when applying to the entire dataset or converged with some issues. Initially we used 0ms as an upper threshold for *µ*_1_, which caused the parameter to converge at this maximal threshold. Using a unique 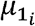 parameter for each participant was theoretically favorable (and indeed used in Study 2), but could not be applied in Study 1, potentially either due to computational challenges (fitting many more inter-dependent parameters) or due to the complex structure of the data (e.g., where anticipatory responses were intertwined with late responses due to changing go signal delay)

##### Computational model of CAT with stimulus-level parameters

In an exploratory model, designed following the analysis of Study 2, we aimed to fit a stimulus-level learning parameter. For each stimulus_s_ which was presented to participant_i_, we fitted an individualized 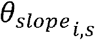 parameter, which was used instead of the participants-level individualized learning parameter 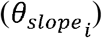, see Supplementary Code S2. Initially, we attempted to model a participant-level dependence, by modeling 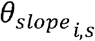 as a parameter derived from higher-level 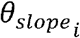 participant-level parameter, similarly to the design in Study 2 (see below). However, such model with tens of thousands of 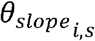 parameter estimates that covaried with hundreds of 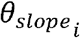 parameters was computationally too demanding. To simplify the model structure, 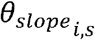 parameter was modeled independently of participants’ identity (i.e., not considering within-participant possible effect; see Supplementary Code S2).

##### Deviation from previous version and pre-registration

In our pre-registration, we used an identical Stan model with one key difference – to enforce an order in which *µ* _1_, *σ*_1_ would relate to early anticipatory RTs and *µ* _2_, *σ*_2_ would relate to late cue-dependent RTs, a different restriction was set in which the upper and lower limit of *µ* _1_ and *µ* _2_, respectively, were set to 0 (Cue onset), instead of the final limits set to 150ms and 200ms. Running the model with these different restrictions converged to a stable solution. Early anticipatory responses were modeled as having an earlier mean (*µ* _1_ = -0.04ms, 95%CI [-0.16, 0.0], 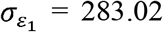, 285.62]) and late cue-dependent responses were very similar to the new model (*µ* _2_ = 287.71ms, 95%CI [287.26, 288.16], 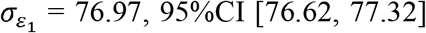]). Other parameters were also very similar *(θ* _*slope*_ = 3.49, 95%CI [3.20, 3.77]), with variation between participants (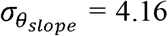, 95% CI [3.93, 4.39]). All reported association with probe phase remained very similar.

While the model showed no formal convergence errors (see trace plots of parameters in Supplementary Fig. S9), the μ_1_ parameter estimate seemed to have been affected by the artificial limitation which was imposed on its maximal value for technical reasons (to avoid inversion with μ_2_ across the different chains). Throughout the different chains, the final parameter converged around the upper limit set to 0 (the Cue onset). After several experimental attempts to change this limit, we found the increasing *µ*_1_ lower limit to 150ms resulted in a better solution, in which μ_1_ parameter estimate did not converge around the uppermost limit. This model is the final model we chose to use and report here.

##### Probe analysis

To evaluate the effect of CAT on preferences we analyzed the proportion of trials in which participants chose the Go stimulus over the NoGo stimulus in the probe phase, using mixed-model logistic regression. As Go and NoGo stimuli were matched based on initial value, under the null hypothesis, participants were expected to choose Go stimuli at 50% of trials (log-odds = 0; odds = 1). The results of this analysis were of main interest in previous publications and is reported in the current work for unpublished data (see supplementary materials).

While early work with CAT task showed a differential effect of value on CAT effect on preferences - i.e. CAT usually had induced more prominent preference modification for stimuli of initial high value, compared to stimuli of initial low value ^12,13^, more recent work with CAT found that this effect was not a dominant feature of CAT ^15,18,19,22^. Thus, in the current work, we pooled together data of stimuli with both high- and low-initial value.

##### Probe and individualized-learning parameter association

Our main analysis of interest aimed to evaluate the association of the CAT preference modification effect with the individualized learning parameter 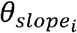, derived from the computational modeling of CAT. In a mixed model logistic regression analysis implemented with lme4 R package ^84^, the number of trial each participant chose the Go versus NoGo stimuli was explained using the independent variable of 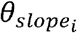 of each participant, as calculated in the preceding CAT task. The data was aggregated over all participants in the 29 experiments. To account for the aggregation of different experiments in the dataset, we also included random intercept and random slope terms for each of the 29 experiments, see logistic regression formula in Equation 2.

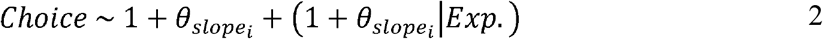

In an additional exploratory analysis (which was conceived after the analysis of Study 2), we modeled choices using a stimulus-specific 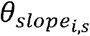 parameter. We aimed to examine if modeling a learning parameter at a more precise stimulus-level (contrary to the pre-registered participant-level 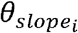 parameter), would serve as a more accurate predictor of future preference modification. For this aim we merged the 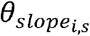 which were estimated for each contingency with the probe data. Choices (number of trials each Go stimulus_s_ was chosen or not by each participant_i_) were modeled with two additional models using 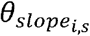 as an independent variable. In the first model, 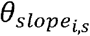 parameter estimates were used instead of 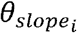 estimates. Since 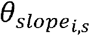 varied within participants, which were nested within experiment, a nested random slope effect for 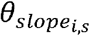 was also added (Equation 3a). In two additional models, probe choices were explained using a full model with both 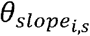 and 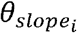 (Equation 3b), and a nested model with the same IVs except for the fixed and random effects of eslope (Equation 3c). Using a likelihood ratio test for nested models (comparing model 3b with the nested model of 3c), we examined whether 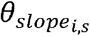 provided additional explanatory above and beyond 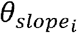.

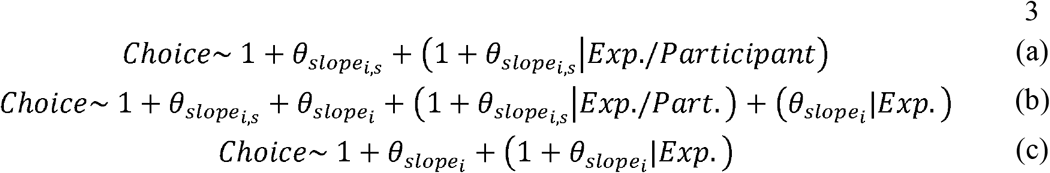

##### Effect size estimation for mixed models with 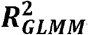

For the reported logistic regression mixed model, we included an additional effect size score using a generalized linear mixed effect model (GLMM) *R*^2^ estimate, based on the work by Nakagawa, and Schielzeth (2013), implemented with R’s Multi-Model Inference (MuMIn) package ^85–88^. Like *R*^2^ in linear mixed model, 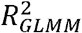 was used to quantify the relative proportion of variance accounted by the three generalized mixed model’s variance components: variance explained by the fixed effects 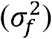, variance explained by the random effects 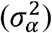, and unexplained residual variance 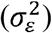. In accordance with the developers’ suggestion ^85,86^, we report here two *R*^2^ scores – the marginal *R*^2^ (equation 4a), which signifies the proportion of total variance accounted by the fixed effects 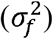; and the conditional *R*^2^ (equation 4b), which signifies the relative proportion of variance explained by both the fixed and random effects of the model.

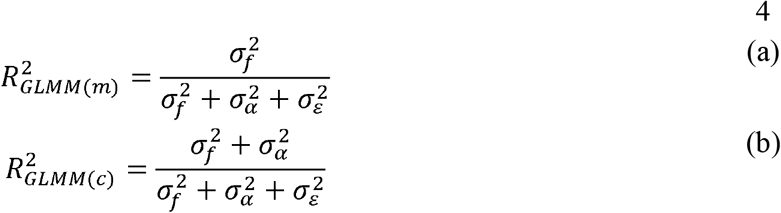

In mixed models examining the contribution of stimulus-level computational marker (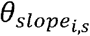; Equation 3) 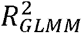 values were similarly evaluated. However, it is important to note that the basic unit of analysis in these model (choices per Go stimulus within participant) differ from the unit of analysis in Equation 2 (choices per participant). Thus the 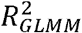 values are not comparable between the two models The 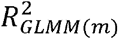 used the participant-level model is the most similar in structure and interpretation to “traditional” *R*^2^ in ordinary linear model, representing the proportion of variance explained by the (fixed) effects of interest, using the participant as the basic unit of analysis.

### Study 2: Novel CAT design

#### Stimuli

In the two novel experiments, the stimuli set included images of 80 unfamiliar faces from the Siblings dataset ^40^, 40 male and 40 female characters. As in previous CAT studies with this stimuli set ^15,18^, the images were processed in Photoshop, so that all stimuli were cropped to identical size (400×500 pixels), with the character’s pupils positioned at the same spatial coordinates and a homogeneous gray background. The characters poised similar neutral expression and had minimal salient artificial characteristics such as jewelry, make-up, or distinct facial hair. Previous CAT studies with these stimuli found that a similar preferences enhancement for both high- and low-value faces which were associated with a Go cue during CAT ^15,18^. Meaning, CAT had similar effect on preferences both for stimuli of initial low-value and of initial high-value. Using face stimuli allowed us to pool together a larger sample of stimuli of different initial values, under the reasonable assumption that the effect would show similar pattern across the different initial value categories.

During the CAT task, a visual Go cue was associated with some of the face stimuli. The Go cue was identical to the one used in a previous study ^15^, it comprised of a semi-transparent Gabor image (38×38 pixels, alpha = 0.7), which appeared at the center of the screen, on top of the face stimuli.

#### Procedure

Like previous CAT experiments, the two novel preliminary and replication experiments included three phases: a baseline preference evaluation task, a modified cue-approach training task, and a post training probe phase which examined preference modification.

##### Initial preference evaluation task

To evaluate initial preferences, participants underwent a binary forced-choice task between random pairs of stimuli, as in previous CAT experiments with non-consumable stimuli ^15,18^. The task included 400 unique choice trials (meaning, no choice between the same two faces repeated more than once), during which each of the 80 face stimuli was presented exactly 10 times. Each choice trial lasted 3000ms, of which participants were given a 2000ms time window to make their choice. Choice trials were followed by a 500ms confirmation screen, showing a green frame around the chosen stimulus, and a fixation cross, which was presented for the remaining trial duration as inter-stimulus interval (ISI), for at least 500ms. In case no choice was made during the 2000ms time-window, a screen saying “You must respond faster” appeared for 500ms, followed by a 500ms ISI.

Binary choices were transformed into individual preference scores using Colley ranking algorithm ^82^. Stimuli were ranked based on each participant’s individual initial preferences from the highest value (1) to lowest value (80). Ranks were used to categorize the stimuli into 10 equal-size value groups (each containing eight stimuli; ranks 1-8, ranks 9-16, etc.). The value categories and internal ranks within each category were used to allocate conditions in the subsequent training and probe task, ensuring initial values were balanced across 100% and 50% contingency conditions, as well as across Go and NoGo stimuli within each value-category, which would be pitted against each other in the subsequent probe phase.

##### Cue-approach training

The CAT task consisted of 20 training runs, in each run all 80 stimuli were presented in a random order for 1000ms each, with a 500ms ISI. In 30% of trials a visual semi-transparent Go cue appeared 850ms following the stimulus onset for 100ms at the center screen position, on top of the face stimulus. Unlike previous CAT experiments, the Go cue onset did not change throughout the training task. Thus, actual RT was consistent with effective RT (RT from cue onset).

Three Go association conditions were included in this modified version of the CAT – 16 stimuli were always presented with the Go cue (Go stimuli, 100% contingency condition), 16 stimuli were associated with the Go cue during half of the presentations (Go stimuli, 50% contingency condition), and the rest of the stimuli were never followed with a Go cue (NoGo stimuli). Participants were instructed to respond to Go stimuli by pressing a keyboard button as fast as they can, before the face-stimulus disappear. Participant were told in the instruction of the three Go cue contingencies, and that they may respond when they see the or even when they anticipate the cue will appear shortly (see the task instruction as presented to the participants in Supplementary Fig. S5). Unlike previous CAT studies, participants were explicitly told of the Go contingency and were thus encouraged to initiate anticipatory responses preceding the actual cue onset, while maintaining high accuracy by only responding when they have sufficient confidence that a cue will follow.

Stimuli were allocated with a go contingency condition based on initial preferences. Stimuli were categorized into 10 equal sized initial value categories, each containing eight stimuli. The stimuli in the highest and lowest initial value categories were allocated to be all NoGo stimuli. Of the middle eight value categories, four were allocated to be 100% contingency and 50% contingency condition category. Within each of the eight middle categories, half of the stimuli were Go stimuli and the other half were NoGo stimuli. The role allocation between categories and within them was designed so that initial values were balanced across the groups E.g., within the second value categories, stimuli ranked 9, 12, 13, and 16 were allocated to be Go stimuli, while stimuli ranked 10, 11, 14, and 15 were allocated to be NoGo stimuli (thus, each group had the same mean rank of 12.5; see illustration in Figure 9). The conditions allocation by initial value was counter-balanced across participants.

**Figure 9.**
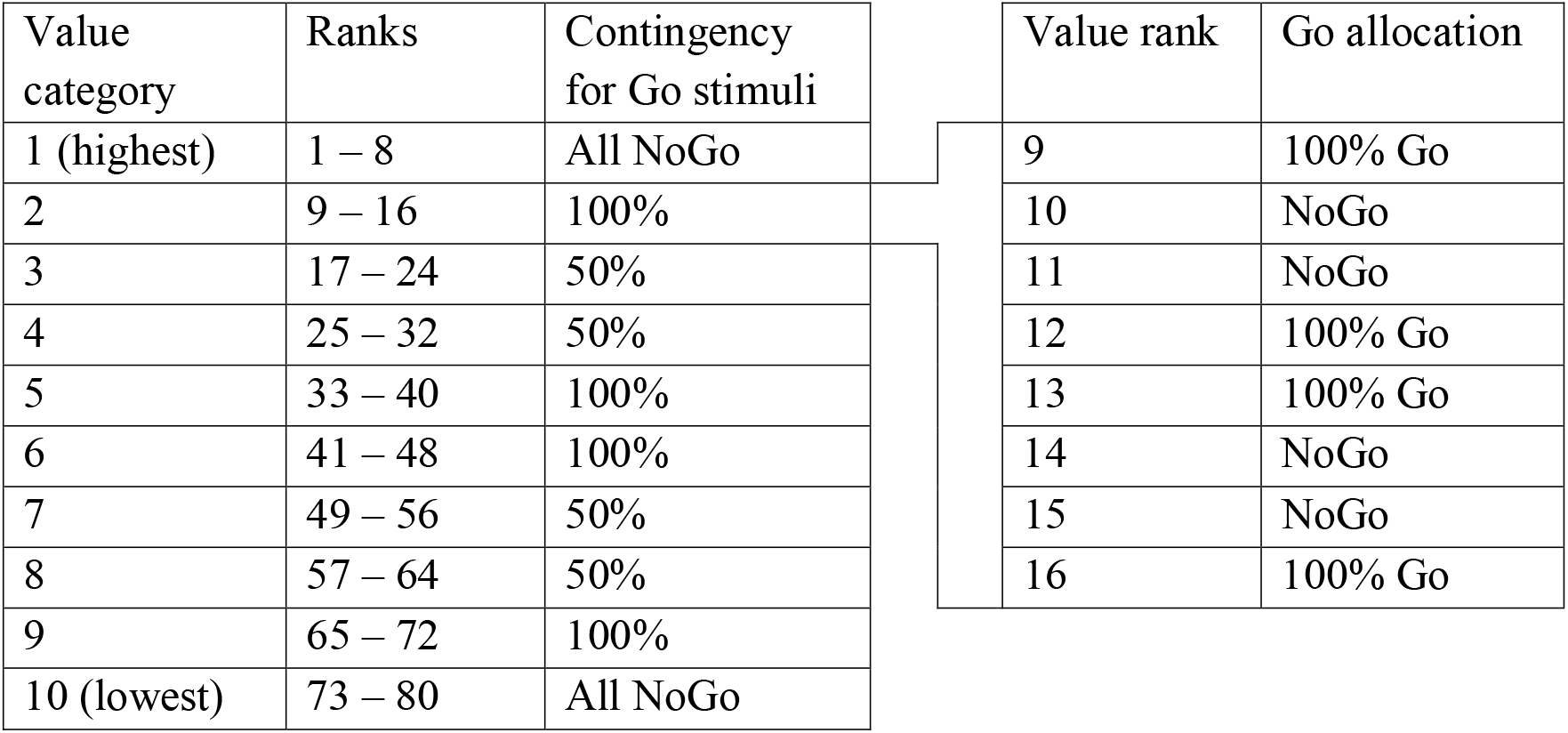
Go stimuli allocation illustration. Both allocations to Go contingency condition (100% versus 50%) and Go/NoGo stimuli were balanced based on initial subjective value. Go stimuli of 50% contingency were associated in 10 out of the 20 training runs with the Go cue. Different designs were counter-balanced across participants by switching between 100% and 50% contingency categories positions and within category Go/NoGo positions.

NoGo stimuli. Of the middle eight value categories, four were allocated to be 100% contingency and 50% contingency condition category. Within each of the eight middle categories, half of the stimuli were Go stimuli and the other half were NoGo stimuli. The role allocation between categories and within them was designed so that initial values were balanced across the groups E.g., within the second value categories, stimuli ranked 9, 12, 13, and 16 were allocated to be Go stimuli, while stimuli ranked 10, 11, 14, and 15 were allocated to be NoGo stimuli (thus, each group had the same mean rank of 12.5; see illustration in Figure 9). The conditions allocation by initial value was counter-balanced across participants.

All Go stimuli of the 50% contingency condition were associated with the Go cue in the last two runs as well as in eight additional runs (in total - 10 out of 20 run). This was done for potential future implementation of the task in an MRI scanner, which have not been done at the time of this manuscript write-up.

#### Probe

In the probe phase, preference modification effect was examined using a binary forced choice task, in which participants chose their preferred stimulus of pairs Go versus NoGo stimuli, of similar initial value. Each Go stimulus was pitted against the four NoGo stimuli within the same initial value category (e.g., Go stimuli ranked 9, 12, 13, and 16 were pitted against NoGo ranked 10, 11, 14, and 15). Participants had 1500ms to make their choice, which was followed by a 500ms confirmation feedback (or a message prompting faster response, in case no choice was made), and an ISI of varying duration (1000ms – 9500ms, *M* = 3000ms), drawn from a truncated exponential distribution with 100ms precision. Like in previous CAT experiments with non-consumable stimuli (and in contrast to CAT experiments with consumable stimuli), choices in the probe phase were not incentive compatible. Previous studies showed that CAT effect is consistent both across incentive compatible choices of consumable stimuli, as well as in non-incentive compatible choices ^15^.

Choices of the 100% contingency condition and 50% contingency condition were merged across the different value categories and analyzed with a mixed-logistic regression model. We hypothesized that participants would choose the Go stimuli over NoGo stimuli, and that this preference modification effect would be more robust for the 100% contingency condition. We also hypothesized that individual 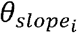 parameter from computational modeling of the training task, would predict which participants would demonstrate stronger preferences modification effect.

### Participants

In the preliminary experiment and replication experiment *n* = 20 and *n* = 59 valid participants completed the experiment and were included in the analysis, respectively. The sample size of the preliminary experiments was based on the minimal sample size used in previous CAT studies. The sample size required for the replication sample was based on a power analysis using the results of the preliminary experiment. A sample size of *n* = 59 was expected to be sufficient to achieve 95% power to detect a significant correlation effect (α = 0.05) between probe choices and the 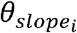 parameter estimates, in both 100% and 50% contingency conditions. The power analysis and the resulting sample sized was documented in the pre-registration and its supplementary materials (https://osf.io/nwr4v).

Three quality-assurance measurements were used as exclusion criteria, based on previous studies with CAT ^15,18,22^ – (1) low variability of Colley scores in the initial preference evaluation task (which indicate intransitive choice pattern), (2) proportion of false alarm during training, and (3) proportion of missed Go trials during training. In each experiment, we excluded participants with extremely low transitivity, high false alarm, or high miss rate (defined as 3SD from the group mean). The exclusion criteria were pre-registered for the replication experiment, along with the planned sample size.

In the preliminary experiment, no participant was excluded due to the mentioned above exclusion criteria (Transitivity score: *M* = 0.205, 3*SD* cutoff = 0.122, min valid score = 0.137; False alarm rate: *M* = 3.62%, 3*SD* cutoff = 15.80%, max valid score = 15.27%; Miss rate: *M* = 6.33%, 3*SD* cutoff = 24.13%, max valid score = 21.04%). In the replication experiment, five participants were excluded due to these pre-registered exclusion criteria: Two participants had low transitivity score, one participant had high rate of false-alarm, and two participants had high rate of missed Go trials (Transitivity score: *M* = 0.213, 3*SD* cutoff = 0.141, *M*_valid_ = 0.216, min valid score = 0.143; False alarm rate: *M* = 4.74%, 3*SD* cutoff = 39.00%, *M*_valid_ = 3.49%, max valid score = 37.59%; Miss rate: *M* = 7.59%, 3*SD* cutoff = 40.22%, *M*_valid_ = 5.97%, max valid score = 38.75%).

### Analysis

Like the procedural design, the analyses of the two experiments were also identical, and generally resembled that of the meta-analysis study, introducing some analysis improvements which could not be applied in the meta-analysis.

#### CAT computational model

The general goal and design of the Bayesian computational framework in the two new experiments resembled that of Study 1. Participants’ RTs were modeled as a mixture of two Gaussian distributions - one distribution of late, cue-dependent responses, and another distribution of earlier anticipatory responses. *θt,i* mixture proportion determined the probability of making an anticipatory response by the participant_i_ at trial_t_. The *θt,i* probability was modeled as a monotonic function of time, using as an individual rate parameter. Unlike the meta-analysis model, in Study 2, we introduced two distinct contingency conditions, thus for each participant two 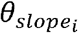 parameters were modeled, namely one for the 100% contingency condition, and one for the 50% contingency condition. Another discrepancy between the meta-analysis model and Study 2 model, was an introduction of individual parameter for anticipatory responses center 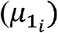, which allowed flexibility in modeling individual differences in anticipatory responses onsets (Equation 5). This model improvement was possible thanks to the more homogenous nature of the data. Each experiment (preliminary and replication) was modeled separately. See the complete model and priors in Supplementary Code S3.

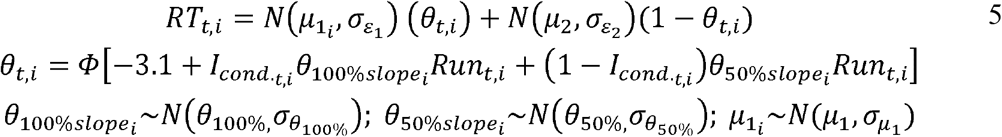

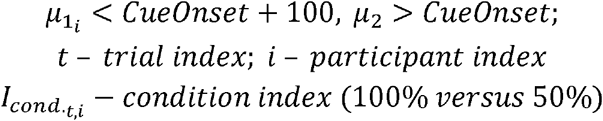

In an additional exploratory analysis (not pre-registered), we aimed to create an even more accurate learning computational marker by modeling a 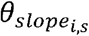 for every stimulus_s_ that participant_i_ was trained with. Assuming that some variability in training speed could be measured not only between participants, but also withing each participant. Since the actual stimuli which were allocated to be Go stimuli varied between participants (i.e., for each participant the condition and role of a certain image stimulus was different), we assumed no mutual information is shared between the same stimulus index of different participants. Stimuli and participants were treated as random effects (Equation 6).

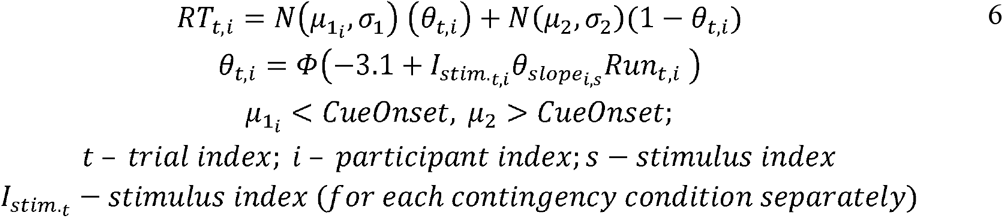

In the process of testing this novel approach, the model did not converge when presented with the full data which included both 100% and 50% contingency conditions. However, when trained twice, using each contingency-condition as an independent dataset, the model did converge. See the complete model and priors in Supplementary Code S4.

#### Probe

As in previous CAT studies, preference modification was evaluated following CAT. Participants’ preferences for Go over NoGo stimuli were categorized into two conditions corresponding with the CAT conditions – of 100% contingency and 50% contingency conditions. Trials of the different value groups were pooled together and analyzed with a mixed model logistic regression. In a post-hoc analysis we validated that the initial value had no significant impact on the conclusions of the analysis (see supplementary materials).

Based on the results of the meta-analysis, we hypothesized that better learning (manifested as larger 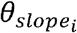 parameter estimate) would induce stronger preference modification effect. Thus, we also hypothesized that a stronger preference modification effect would be observed for the 100% contingency condition, compared with the 50% contingency condition. To examine this hypothesis, we run a one-sided mixed model logistic regression with an additional fixed and participant-based random explanatory variables (EVs) of the contingency condition (Equation 7).

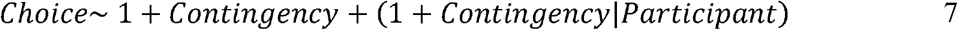

#### Probe choice prediction based on individual learning parameter

Most importantly, we hypothesized that using the 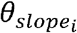 parameter estimate for learning would predict future preference modification effect observed during the subsequent probe phase. To test this hypothesis, we introduced the 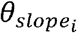 as an additional independent variable in the mixed model logistic regression. Choices were modeled using contingency condition (both as fixed effect and as a random slope between participants), 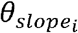, and an interaction term. This model is equivalent to modeling choices using an intercept and 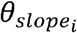 slope separately for each contingency condition with a random intercept modeled within participants (Equation 8a).

In an additional (not preregistered) exploratory analysis, we modeled a stimulus-specific 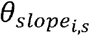 parameter. We aimed to examine if modeling a learning parameter at a more detailed stimulus-level (extending the pre-registered participant-level 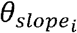 parameter), would serve as a more accurate predictor of future preference modification. For this aim we merged the 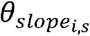 which were estimated for each contingency with the probe data. Choices were modeled with two additional models using a 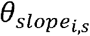 as an independent variable. In the first model, 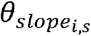 parameter estimates were used instead of 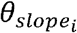 estimates. Since 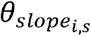 varied within participant, a random slope effect for 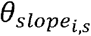 was also added (Equation 8b). In another model, probe choices were explained using a full model with both 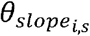 and 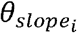 (Equation 8c). This model was used to examine whether 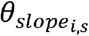 provided additional explanatory power in a likelihood ratio test for nested models (comparing the model of 8c with the nested model of 8a).

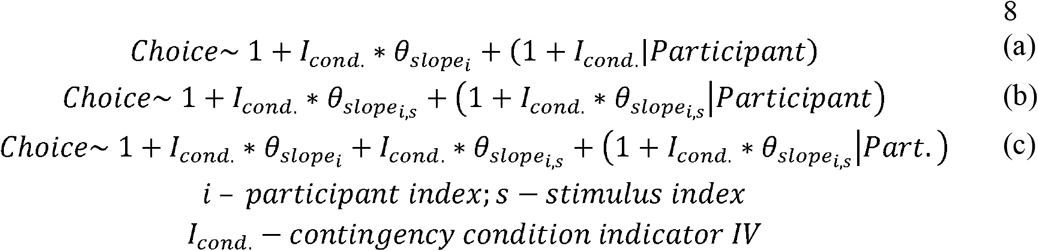

When results with 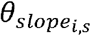 were analyzed, we found that some participants were fitted a 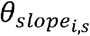 estimates of extremely small variability (SD<0.1, meaning their 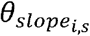 estimates were effectively fixed; see Supplementary Fig. S6). In such cases a mixed model with 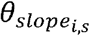 random slope (Equation 8b and 8c), resulted in convergence warnings (this phenomenon was not observed in Study 1). Thus, for these analyses, we excluded all participants which demonstrated such low variability of 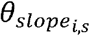 estimates in either one of the two contingency conditions. This resulted in exclusion of nine participants in the preliminary experiment, and 14 participants in the replication experiment (out of 20 and 59 participants, respectively). For the nested model likelihood ratio analysis (comparing a full model to model without 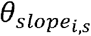 independent variables), both models were examined using the same subset of 11 and 45 valid participants.

#### Effect size estimation for mixed models with 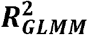

As in the meta-analysis study, we report for the logistic regression mixed models two 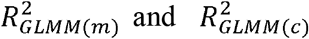, denoting respectively the marginal and conditional R2 effect of the generalized linear model ^85– 88^.

## Funding

This work was supported by the European Research Council (ERC) under the European Union’s Horizon 2020 research and innovation programme (grant agreement n° 715016) and Under the European, and the Israeli Science Foundation (1798/15 and 1996/20) granted to Tom Schonberg. Tom Salomon was supported by the Nehemia Levtzion fellowship and the Fields-Rayant Minducate Learning Innovation Research Center.

## Competing interests

The authors declare they have no competing interests.

## Data and materials availability

All materials and analysis codes are available online at: https://github.com/tomsalomon/CAT_Individualized_Learning. Preregistration hypotheses, experimental design data and codes used for power analysis are available in the Open Science Foundation (OSF) depository (https://osf.io/nwr4v).

## Notes

https://github.com/tomsalomon/CAT_Individualized_Learning

### Competing Interest Statement

The authors have declared no competing interest.

### Summary of Updates

The current version of the manuscript addresses suggestions made by external reviewers during the submission process for publication. We included new analyses that demonstrate the added value of the computational model over more simplified solutions and control for potential confounds or intervening factors such as response speed during probe choices and the initial value of the stimuli. We also elaborate more on topic points made by the reviewers in the intro and the discussion and include additional findings and analyses in the supplementary materials.

