## Supplementary Materials for "A computational model for individual differences in non-reinforced learning for individual items"

**This PDF file includes:**

Supplementary Text

Figs. S1 to S18

Codes S1 to S5

Table S1

References

Supplementary Text

Unpublished data included in the CAT meta-analysis

The meta-analysis of CAT included data from 29 different CAT experiments; ten of these experiments were from data which were not published as they were collected for pilot testing before running another published study or would be published in the future. In this section we describe the unique features in these experiments and their results.

Overall, in most experiment we found that CAT resulted in enhanced preferences for the associated Go stimuli. The results of the studies are summarized in Supplementary Fig. S10 and its corresponding Supplementary Table 1. For each experiment below we describe its unique context and methods, as well as the results if those differed from the general trend of enhance preference effect for CAT stimuli over NoGo stimuli of similar initial value.

The proportion of trials in which participants chose Go over NoGo stimuli was analyzed using mixed model logistic regression (with random intercept modeled per participant). For consistency with previous work with the CAT task (Bakkour et al., 2016; Schonberg et al., 2014), we analyzed separately choice between high-value stimuli, and choices between low-value stimuli. However, based on more recent finding with non-consumable stimuli as well as work that applied an appropriate mixed-model analysis on the data (Botvinik-Nezer, Bakkour, Salomon, Shohamy, & Schonberg, 2021; Salomon et al., 2018; Salomon, Botvinik-Nezer, Oren, & Schonberg, 2019), we did not expect to find a significant difference between the two value categories. Thus, we did not directly compare the effect of value category.

*Experiments 10-11: familiar faces.* In two experiments we performed CAT study using stimuli of familiar faces of famous past and present Israeli parliament member. The stimuli set comprised of equal proportion of left-wing, right-wing, and central-trending political figures. The aim of these experiments was to examine CAT effect with stimuli for which participants would have strong initial preference variability (e.g., a participant with right-wing tendencies should have strong preference for right-wing politicians over left-wing politicians). The procedure followed the protocol of similar CAT experiments with non-consumable stimuli – initial preferences evaluation using binary choices, training phase with 20 training runs, and probe phase in which Go stimuli were pitted against NoGo stimuli with similar initial value (both either high-value or low-value). Both experiments were identical in design, as the second experiment was designated to be a direct replication of the first experiment (*n* = 25) using larger sample size in the replication experiment (*n* = 39).

*Experiments 13-14: emotional faces.* In two experiments, participants were exposed to stimuli set of 80 faces from the Karolinska face dataset (Lundqvist, Flykt, & Öhman, 1998); the stimuli set comprised of 40 unique figures (21 male and 19 female), posing a neutral facial expression and a positive facial expression. Participants performed a CAT study design in which (unbeknownst to the participants) preferences were evaluated for each emotional category separately. During probe participants chose from pairs of similar value and same facial expression (happy or neutral), in which one stimulus was a Go stimulus. One important goal of these experiments was to examine whether CAT had a stronger preference modification effect for faces of positive affect than neutral one.

In both experiments we found that CAT enhanced preferences for the associate Go stimuli, both for the face stimuli of positive affect and neutral affect). However, contrary to our initial hypothesis, we did not consistently observe a stronger preference modification effect for the positive affective face stimuli (see Supplementary Fig. S10 and Supplementary Table 1).

*Experiments 16: snacks adolescence.* In this CAT experiment with snack food stimuli, the effect of CAT was probed in a group of adolescent participants and young adults, aged 14 – 20 (mean age 18.2, SD = 1.90). The experiment protocol followed that of other CAT experiments with snacks (Bakkour, Lewis-Peacock, Poldrack, & Schonberg, 2017; Schonberg et al., 2014), using a relatively short training of 12 training runs. The effect of CAT on preferences was not fully replicated in this sample. While participants demonstrated enhance preferences for the low-value Go stimuli over low-value NoGo stimuli, for the high-value Go stimuli an enhance preferences effect was not observed (see Supplementary Fig. S10 and Supplementary Table 1). This unexpected result could be due to less robust effect associated with the shorter training procedure or unique features of the population sampled this study.

*Experiments 17: snacks with rewarding cue.* This CAT experiment with snack food stimuli, followed a similar protocol to previous CAT experiments. In the training phase, the neutral Go Cue was replaced with a visual cue, indicating to the participants that they have won a monetary reward, which accumulated as training progressed. Reward was not dependent of performance in the task, and thus did not explicitly serve as an incentive to engage with the task. This experiment was used as pilot testing for a future experiment comparing reinforced with non-reinforced association learning.

*Experiments 20-21: CAT pilot experiments with snacks*. These two experiments were used as pilot experiments, before running Experiment 29 in fMRI setting (Botvinik-Nezer, Salomon, & Schonberg, 2020). In Experiment 20 a short training procedure with 12 training runs was used, which did not result in a significant preference modification effect (Supplementary Figure S9 and Supplementary Table S1). Following these results, in the second pilot experiments, which we refer to in this work as Experiment 21, a longer training procedure with 20 training runs was used. This procedural design was eventually selected for the final imaging experiment (Experiment 29).

Using simple alternative marker of learning

In an additional analysis, conducted following all other analyses, we examined the robustness of $\theta_{slope_{i}}$ as a prediction of subsequent probe choices above and beyond a simple alternative marker. As an alternative to $\theta_{slope_{i}}$, we calculated a simple anticipation score per participant. Training RTs were classified as ‘*anticipatory*’ based on the threshold of the bottom 1% of all RTs during the first training runs. In the meta-analysis this threshold was RT of 144.5ms or less, while in Study 2’s experiments this threshold was 210.2ms. We then calculate per participant the proportion of RTs below this threshold and used that proportion score as a simple alternative marker, which should model similar pattern as the $\theta_{slope_{i}}$. We also repeated the process for individual stimuli, to define a participant-stimulus alternative marker comparable with $\theta_{slope_{i,s}}$, and in Study 2, a proportion score was calculated for each condition (50% and 100% contingency).

We first examined the similarity of the proportion of anticipatory responses score with $\theta_{slope_{i}}$. In the meta-analysis, the proportion of anticipatory responses were found to be strongly correlated with $\theta_{slope_{i}}$, *r* = 0.68, *t*(862) = 27.02, *p* = 5.5e^-117^, 95%CI [0.63, 0.73]. However, in study 2 preliminary exp., the proportion of anticipatory responses were positively correlated with $\theta_{slope_{50{\%}_{i}}}$ (*r* = 0.57, *t*(18) = 1.93, *p* = 0.008, 95%CI [0.17, 0.98]), but to a lesser degree with $\theta_{slope_{100{\%}_{i}}}$ (*r* = 0.41, *t*(18) = 2.15, *p* = 0.08, 95%CI [-0.05, 0.86]). In Study 2 replication exp. there was no significant association between anticipatory responses and $\theta_{slope_{i}}$ within each condition; replication exp. 50% condition: *r* = 0.11, *t*(57) = 0.8, *p* = 0.428, 95%CI [-0.159, 0.369]; replication exp. 100% condition: *r* = 0.10, *t*(57) = 0.78, *p* = 0.441, 95%CI [-0.161, 0.366]. Examining the correlation plots (see Supplementary Fig. S11), it seems that very low $\theta_{slope_{i}}$ were commonly associate with close to 0% anticipatory responses throughout the training, however the proportion of anticipatory responses did not increase monotonically with $\theta_{slope_{i}}$.

To evaluate the predictive power of the proportion of anticipatory responses as a simple alternative to $\theta_{slope_{i}}$, we repeated the same logistic regression models using the proportion of anticipatory responses as a sole independent variable instead of $\theta_{slope_{i}}$. We examined if the simple alternative marker could predict probe choices in the subsequent probe phase and compare the two models’ AIC score. In Study 1, we found that the proportion of anticipatory responses were predictive of preference modification effect (OR = 5.49, 95% CI = [2.80, 10.66], *Z* = 4.99, *p* = 6.1E^-7^; two-sided mixed model logistic regression), and even provided better explanatory power compared with the original model using $\theta_{slope_{i}}$ (ΔAIC = -560.68). This pattern was not found in Study 2, where the proportion of anticipatory responses were not found to be strongly indicative of subsequent preference modification effect (preliminary exp. 50% contingency: OR = 2.07, 95% CI = [0.22, 19.28], *Z* = 0.64, *p* = 0.26; preliminary exp. 100% contingency: OR = 1.98, 95% CI = [0.24, 16.31], *Z* = 0.63, *p* = 0.26; replication exp. 50% contingency: OR = 0.94, 95% CI = [0.41, 2.15], *Z* = -0.64, *p* = 0.55; replication exp. 100% contingency: OR = 1.39, 95% CI = [0.54, 3.54], *Z* = 0.69, *p* = 0.25; one-sided mixed model logistic regression), and using the proportion of anticipatory responses provided worse explanatory power compared to $\theta_{slope_{i}}$ (preliminary exp. ΔAIC = 10.95; replication exp. ΔAIC = 11.01).

To examine the added value of $\theta_{slope_{i}}$ above and beyond the simple proportion of anticipatory responses marker, we examined the significance of $\theta_{slope_{i}}$ in a nested model using both indicators to predict probe choices. In Study 1, $\theta_{slope_{i}}$ provided no additional explanatory power above and beyond the simple proportion of anticipatory responses indicators (ΔAIC = 1.81, $\chi_{\left( 1 \right)}^{2}$ = 0.19, *p* = 0.662; likelihood ratio test). In both experiments of Study 2, $\theta_{slope_{i}}$ significantly improved a predictive model using only anticipatory responses (preliminary exp. ΔAIC = 7.56, $\chi_{\left( 2 \right)}^{2}$ = 11.56, *p* = 0.003; replication exp. ΔAIC = 7.83, $\chi_{\left( 2 \right)}^{2}$ = 11.83, *p* = 0.003; likelihood ratio test).

Using alternative log-normal Bayesian model

In a post-hoc analysis, conducted following all other analyses, we examine the impact of changing the Gaussian Bayesian model with a log-normal modeling of RT distribution. The model had the same restrictions as the gaussian one, meaning the mean of anticipatory RTs must be lower than that of the cue-dependent RTs.

A few adaptions were required in order to help the log-normal model to converge: (i) we added a fixed value of 1,000ms to the effective RT to avoid negative values. (ii) We changed the RT scale to seconds (divided by 1000, while keeping milliseconds precision). (iii) we had to exclude out of this analysis participants which has a large number of more than 30% non-response. This resulted in exclusion of 26 participants from Study 1 meta-analyses and one participant from Study 2 replication experiment). The model is fully detailed in Supplementary Code S5.

*Meta analysis results:*

After running four independent chains, the Bayesian model converged to a stable solution (see Supplementary Fig. S12). The converged model estimated effective RTs as a time-dependent mixture of two RT distributions: one of early anticipatory responses ($\mu_{1}$ = 24.3ms, 95%CI [21.2, 28.4], $\sigma_{\varepsilon_{1}}$ = 298.2ms, 95%CI [295.6, 300.8]) and late cue-dependent responses ($\mu_{2}$ = 285.3ms, 95%CI [284.0, 286.6], $\sigma_{\varepsilon_{2}}$ = 60.5ms, 95%CI [60.3, 60.8]). Overall, participants demonstrated an increase in proportion of anticipatory responses as training progressed, as manifested in the positive group-level parameter ($\theta_{slope}$ = 3.46, 95%CI [3.16, 3.75]), with variation between participants ($\sigma_{\theta_{slope}}$ = 4.17, 95% CI [3.93,4.41]).

Posterior predictive checks (simulated distributions of RT based on estimated model parameters) revealed a good fit of the model to the actual data. Simulated posterior distributions recreated the patterns observed empirically, showing more rapid transition to anticipatory responses in participants with higher $\theta_{slope_{i}}$ parameter estimate as well as capturing the skewed right tail shape of RTs (Supplementary Fig. S13).

Similarly to the two-gaussian model, we examined whether variation in the slope parameter fit to RTs correlated with probe performance. The meta-analysis showed $\theta_{slope_{i}}$ was positively associated with the preference modification effect – i.e., participants with higher $\theta_{slope_{i}}$ parameter estimate, also demonstrated greater odds of choosing Go stimuli (OR = 1.05, 95% CI = [1.03, 1.07], *Z* = 6.05, *p* = 1.4E^-9^; one-sided mixed model logistic regression; Supplementary Fig. S14). The model’s intercept was significantly greater than zero, i.e., even when extrapolated to very low $\theta_{slope_{i}}$ value, the model forecasted enhanced preference for Go stimuli (intercept odds = 1.22, 95% CI = [1.15‚ 1.30], *Z* = 6.8, *p* = 1.0E^-11^).

*Study 2 experiments results:*

After running four independent chains, the Bayesian model converged to a stable solution (see Supplementary Fig. S15). The converged model estimated effective RTs as a time-dependent mixture of two RT distributions: one of early anticipatory responses (preliminary experiment: $\mu_{1}=-72.2ms, 95\%CI\left[ 95.1,39.2 \right], \sigma_{\epsilon_{1}}=263.6ms, 95\%CI[246.0,284.0]$; replication experiment: $\mu_{1}=-90.6ms, 95\%CI\left[ -104.2,-29.5 \right], \sigma_{\epsilon_{1}}=264.9ms, 95\%CI[258.6,277.6]$), and late cue-dependent responses (preliminary experiment: $\mu_{2}=339.1ms, 95\%CI\left[ 336.4,340.4 \right], \sigma_{\epsilon_{2}}=58.6ms, 95\%CI[51.2,59.7]$; replication experiment: $\mu_{2}=329.7ms, 95\%CI\left[ 323.1,343.1 \right], \sigma_{\epsilon_{2}}=59.7, 95\%CI[58.6,60.7]$). Overall, participants demonstrated an increase in proportion of anticipatory responses as training progressed, as manifested in the positive group-level parameter (50% condition: $\theta_{slope}$ = 1.59, 95%CI [0.92, 2.23]; 100% condition: $\theta_{slope}$ = 3.87, 95%CI [2.38, 4.1]), with variation between participants (50% condition: $\theta_{slope}$ = 1.82, 95%CI [1.38, 2.37]; 100% condition: $\theta_{slope}$ = 2.77, 95%CI [2.25, 3.41]).

Posterior predictive checks (simulated distributions of RT based on estimated model parameters) revealed a good fit of the model to the actual data. Simulated posterior distributions recreated the patterns observed empirically, showing more rapid transition to anticipatory responses in participants with higher $\theta_{slope_{i}}$ parameter estimate (Supplementary Fig. S16).

Examining the association of $\theta_{slope_{i}}$ parameter estimates with Go choices during probe, we found positive association with subsequent probe choice both in preliminary experiment (50% condition: log-OR = 0.03, *Z* = 2.45, *p* = 0.024, OR = 1.03, 95% CI [1.02, 1.04]; 100% condition: log-OR = 0.04, *Z* = 3.75, *p* = 0.001, OR = 1.04, 95% CI [1.03, 1.51]; one-sided mixed model logistic regression). In the replication study just in 50% condition found to be significant (log-OR = 0.014, *Z* = 2.12, *p* = 0.038, OR = 1.01, 95% CI [1.008, 1.02]; 100% condition: log-OR = 0.003, *Z* = 0.71, *p* = 0.48, OR = 1.003, 95% CI [0.998, 1.008]; one-sided mixed model logistic regression).

Controlling for initial value in Study 2

In an additional post-hoc analysis, we aimed to examine the impact of initial value on the preference modification effect during probe. In study 2 experiments, 80 stimuli were rank-ordered and categorized into ten value-groups of eight stimuli each (stimuli ranked 1-8 was categorized into the highest value group, stimuli 9-16 comprised the second-highest value group, etc.). The middle eight value groups each contained four Go stimuli and four NoGo stimuli of similar initial value, which were bided against each other in the probe trials (see details in methods section and Figure 9). Early CAT studies that used snack food stimuli and two value categories (high-value and low-value stimuli), reported a stronger preference modification effect for Go stimuli in the high value category, compared with low-value stimuli (Bakkour et al., 2017; Schonberg et al., 2014). However, this differential value effect has not been consistently replicated in later experiments (Bakkour et al., 2016; Salomon et al., 2018, 2019), which also used non-consumable stimuli, similar to the face stimuli used in Study 2 of the current work.

Under the assumption that the effect of CAT on choices is consistent across value categories, we analyzed and reported in the main text the aggregated results from all value categories combined. To validate this assumption, we reanalyzed the probe data using categorical variables representing the eight value groups and examine the significance of an interaction term of value-group with contingency. To evaluate the statistical significance of the (value X contingency) interaction effect, we compared a full model consisting of both main effects and interaction, with a restricted model containing only the main effects. Similarly, to examine the main effect of value-group, we compared a model containing the two main effects to restricted model in which the coefficients associated with value-group were omitted.

In the preliminary experiment we did not find significant value-group and contingency interaction effect (ΔAIC = -4.94, $\chi_{\left( 7 \right)}^{2}$ = 9.06, *p* = 0.249; likelihood ratio test). We did find a main effect for value group (ΔAIC = 16.34, $\chi_{\left( 7 \right)}^{2}$ = 30.34, *p* = 8.2E^-5^) as well as a main effect for contingency condition (ΔAIC = 5.74, $\chi_{\left( 1 \right)}^{2}$ = 7.74, *p* = 0.005). In the replication experiment we identified a significant interaction effect (ΔAIC = 4.35, $\chi_{\left( 7 \right)}^{2}$ = 18.35, *p* = 0.010), with no main effect for value group (ΔAIC = -10.11, $\chi_{\left( 7 \right)}^{2}$ = 3.89, *p* = 0.792) and a significant main effect for contingency condition (ΔAIC = 30.08, $\chi_{\left( 1 \right)}^{2}$ = 32.08, *p* = 1.5E^-8^), see Supplementary Fig. S17.

Descriptively, the value group effects did not show a consistent or monotonic trend, neither within-experiment nor between experiments. In the preliminary experiment, a main effect of value group was manifested as smaller overall preference for Go stimuli in some value categories (e.g. 54.6% choices of Go stimuli in the 7^th^ value group compared with 70.1% in the higher 5^th^ value group; See Supplementary Fig. S17). This trend was not monotonic, i.e. we did not observe a more robust preference modification effect in the higher value groups, nor replicated in the following experiment. Likewise, in the replication experiment an interaction effect was detected, which indicated variability in the contingency effect across different value groups. Like in the preliminary experiment, this effect was not monotonic with the value group (higher value groups did not show consistently more or less robust contingency effects), nor was it found in the previous experiment. In contrast to the inconsistent effects of value group, the effect of cue-contingency was more robust and was replicated in both experiments, above and beyond the contribution of value categories.

In an additional post-hoc analysis, we further examined the impact of adding the (standardized) value difference between Go and NoGo stimuli as covariates to the probe analysis. We found that in both experiment value differences had a significant explanatory power when accounting for participant choices (preliminary experiment: OR = 2.00, 95% CI = [1.07, 1.33], *Z* = 3.24, *p* = 0.002; replication experiment: OR = 1.16, 95% CI = [1.09, 1.23], *Z* = 5.13, *p* = 5.5E^-7^; two-sided mixed model logistic regression). Importantly however, the effect of enhanced preferences for Go stimuli remained consistent above and beyond value-differences effect (preliminary experiment, 50% contingency: OR = 1.33, 95% CI = [1.09, 1.62], *Z* = 2.86, *p* = 0.004; 100% contingency: OR = 1.81, 95% CI = [1.28, 2.57], *Z* = 3.38, *p* = 7.3E^-4^; replication experiment, 50% contingency: OR = 1.41, 95% CI = [1.24, 1.59], *Z* = 5.50, *p* = 3.8E^-8^; 100% contingency: OR = 1.97, 95% CI = [1.60, 2.43], 6.40, *p* = 1.6E^-10^; one-sided mixed model logistic regression).

Cross validation of Study 2 predictions

To validate the generalizability of our prediction model, we examined the effect of using out-of-sample data predictions. In a five-fold cross validation (CV) analysis, the data of all users were split into five groups. In each CV iteration, predictions of the proportion of Go choices was generated to the validation group, based on a mixed logistic regression model that was trained using the data of remaining four groups. In case our model suffered from high overfit, we would expect that the CV predictions would be dissimilar to the model reported in the main manuscript, which was based on the entire dataset.

Examining the correlation between the predictions made by the full data model (reported in the main manuscript), to the predictions made for the held-back data in the CV analysis, we found that in both experiments, the two prediction models resulted in nearly predictions (preliminary experiment: *r* = 0.975, mean absolute prediction difference = 1.51%, range = [0.04%, 4.53%]; replication experiment: *r* = 0.994, mean absolute prediction difference = 0.53% range = [0.002%, 2.43%]; see Fig. S18.). These results imply that the prediction model was not sensitive to the specific data it was exposed to, thus these results are reassuring that the prediction model is likely to generalize well to out-of-sample data as well.

Fig. S1.

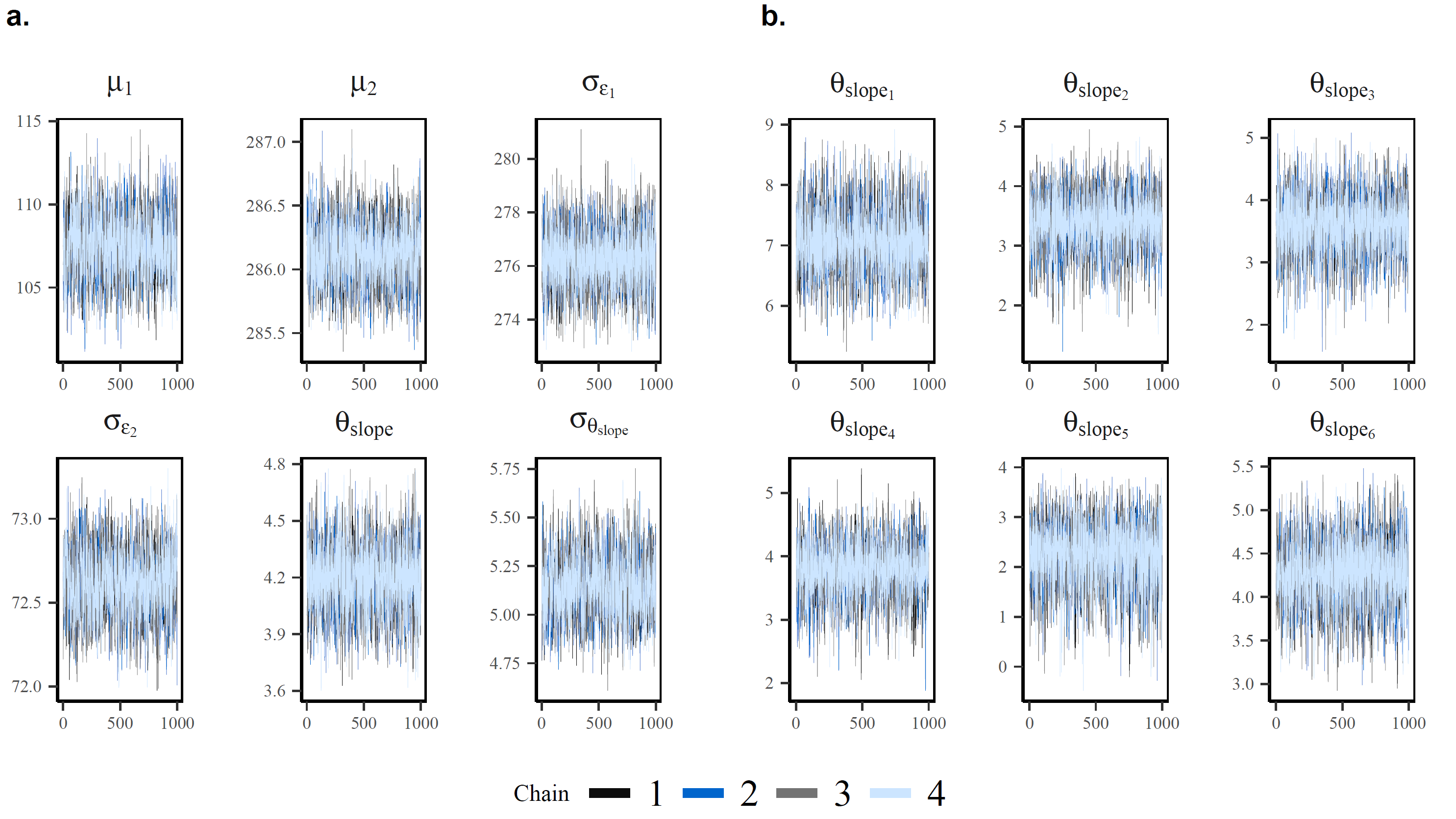

Trace plots of the Stan model of the meta-analysis. (a) Six hyper-parameters defined the shape of the RT data: two means ($\mu{}_{1}, \mu_{2}$) and standard deviations ($\sigma_{\varepsilon_{1}},\sigma_{\varepsilon_{2}}$) for the normal distribution of anticipatory and cue-dependent responses, and a mean and *SD* ($\theta_{slope}$, $\sigma_{\theta_{slope}}$) for the distribution from which individualized $\theta_{slope_{i}}$ parameters would be drawn for each participant. (b) Trace plot of the first six participants’ $\theta_{slope_{i}}$ parameters. A well-converged parameter is characterized with all four chains (in different colors) reaching stable solution around the same estimate for each parameter (‘hairy caterpillar’ pattern).

Fig. S2.

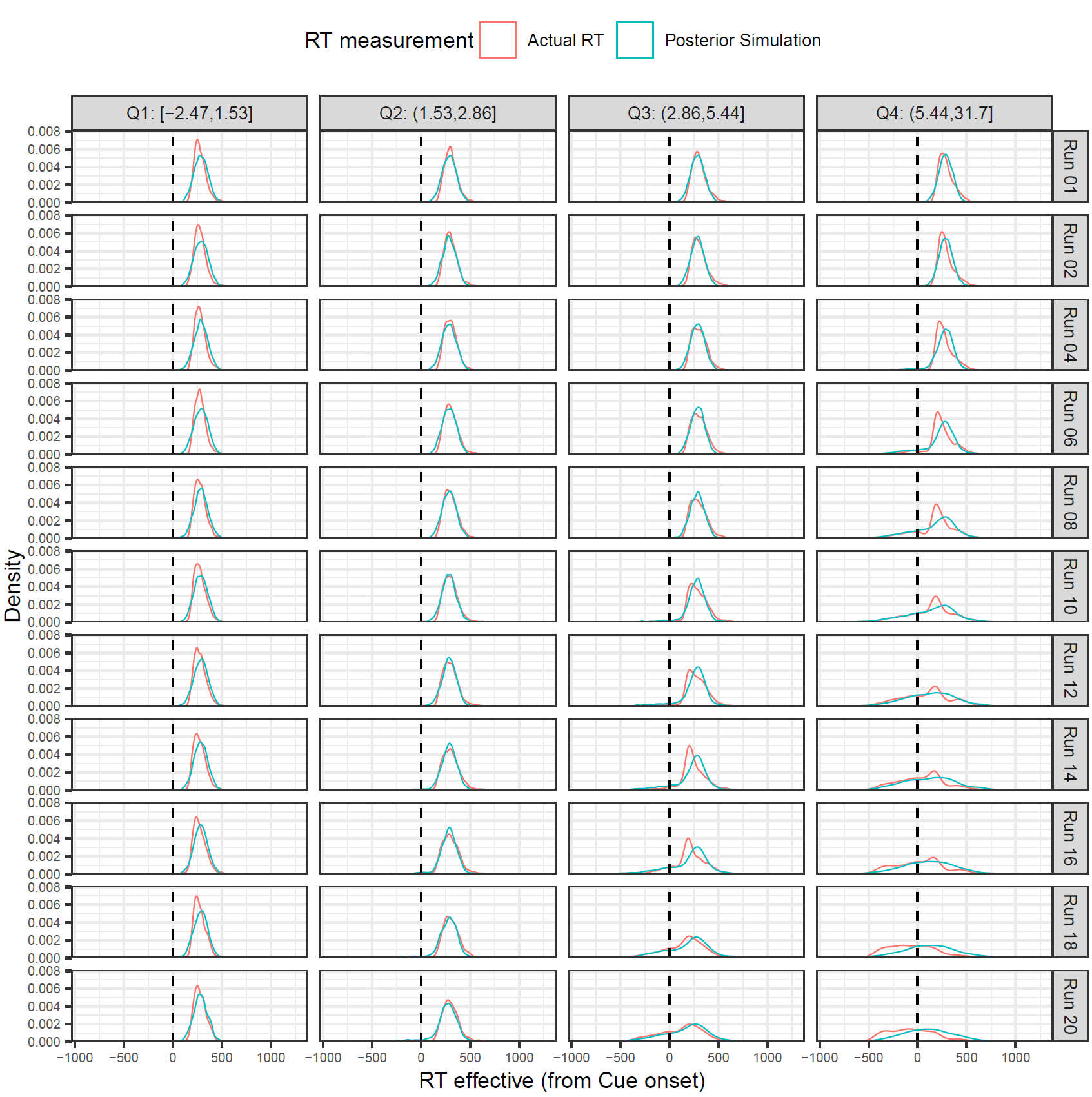

Actual RT distributions (red) versus simulated posterior distributions (teal), by $\theta_{slope_{i}}$ quantile group and run. Participants were categorized into four equal quantile groups, according to their ${\theta_{slope}}_{i}$ parameter estimates (denoted here as Q1-Q4; columns). Throughout the different training runs (rows; included here every other run) the posterior simulated RT distributions fitted well to the transition pattern.

Fig. S3.

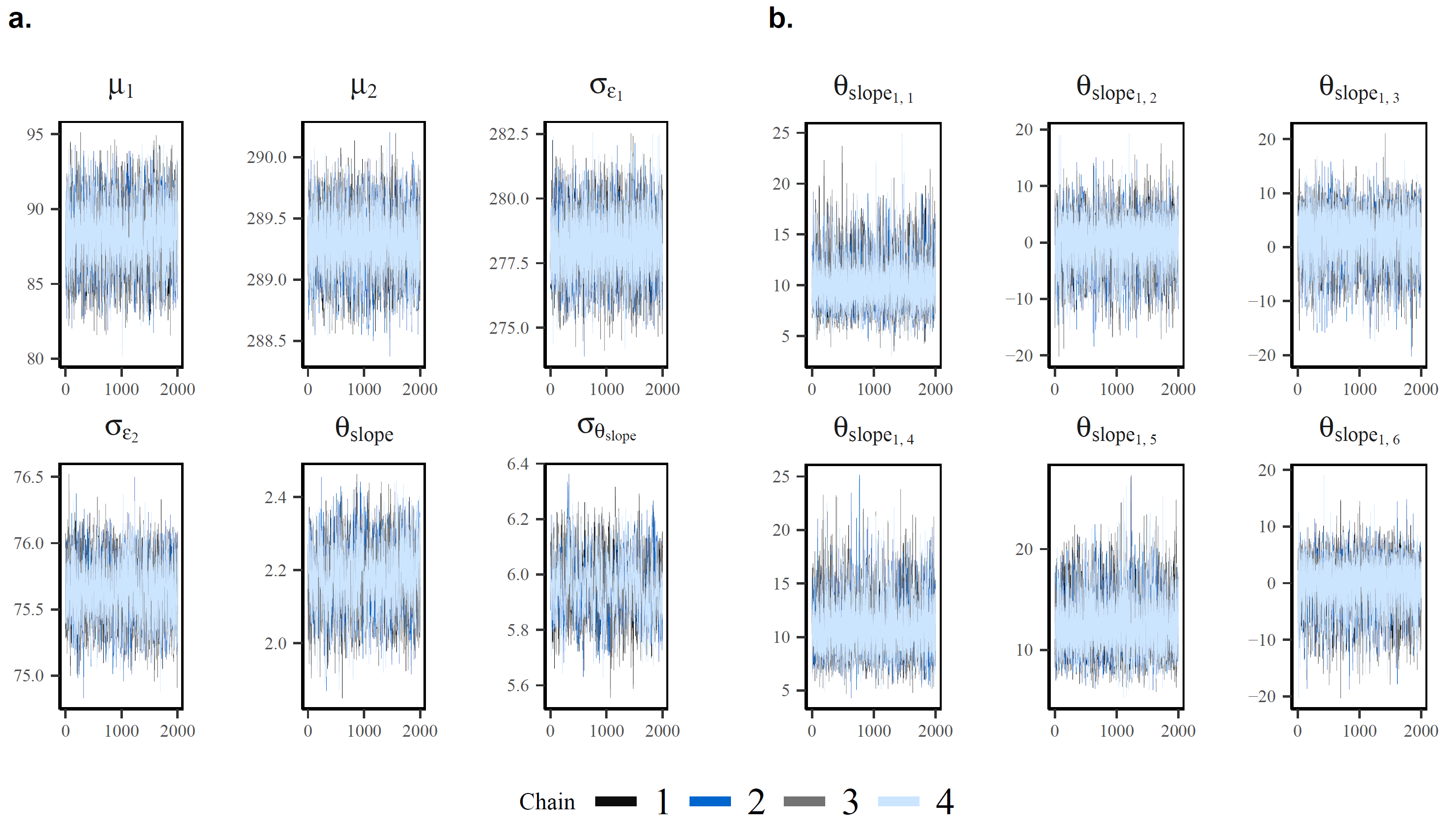

Trace plots of the meta-analysis Stan model, with stimulus-level ${\theta_{slope}}_{i,s}$parameter. (a) Six hyper-parameters defined the shape of the RT data: two means ($\mu{}_{1}, \mu_{2}$) and standard deviations ($\sigma_{\varepsilon_{1}},\sigma_{\varepsilon_{2}}$) for the normal distribution of anticipatory and cue-dependent responses, and a mean and *SD* ($\theta_{slope}$, $\sigma_{\theta_{slope}}$) for the distribution from which individualized $\theta_{slope_{i},s}$ parameters were drawn for each participant. (b) Trace plot of the first six $\theta_{slope_{i,s}}$ parameters, associated with six stimuli of the first participant in the meta-analysis. A well-converged parameter is characterized with all four chains (in different colors) reaching stable solution around the same estimate for each parameter (‘hairy caterpillar’ pattern).

Fig. S4.

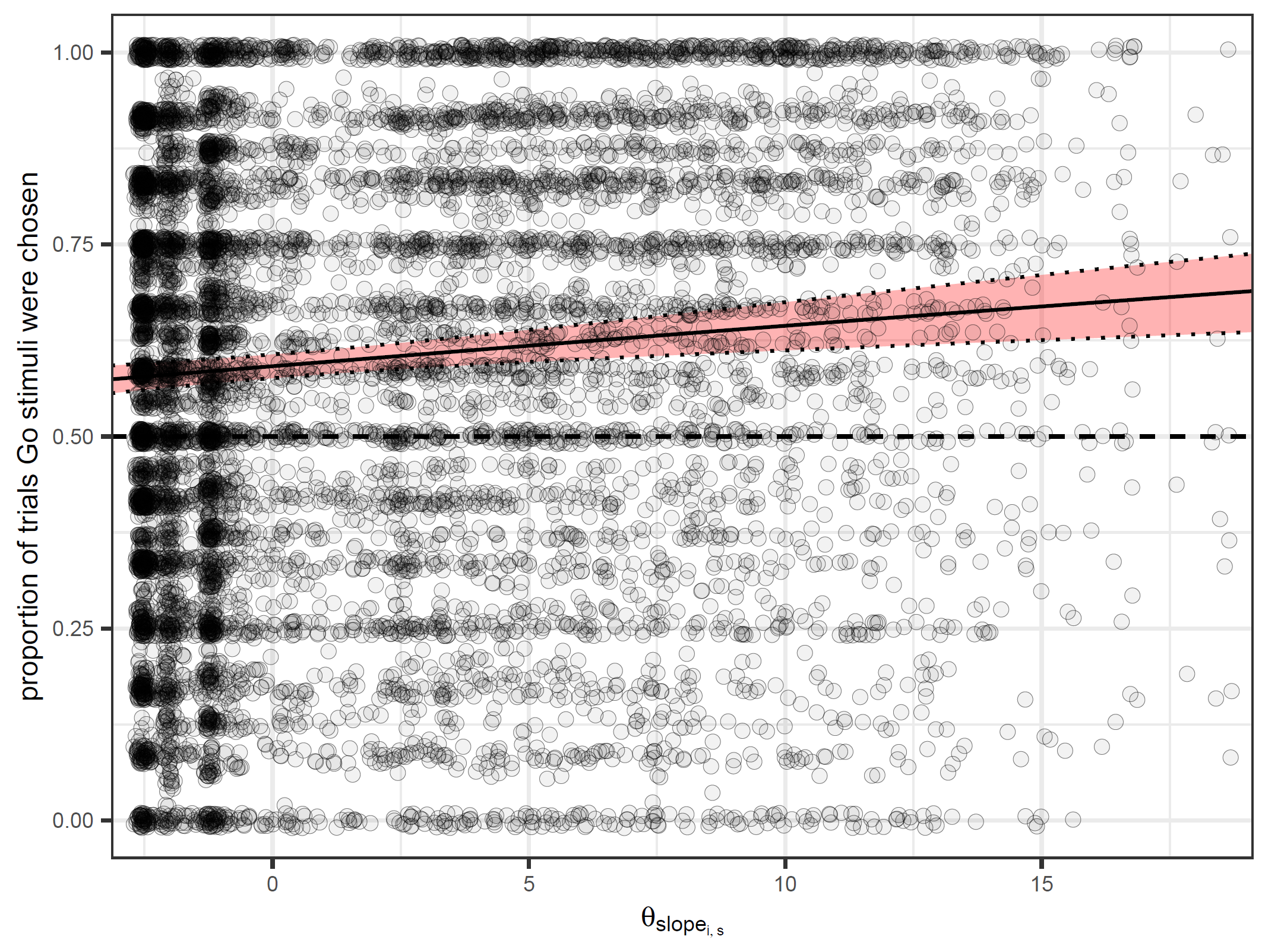

Association of probe choices with stimulus-level computational marker in study 1. Larger $\theta_{slope_{i,s}}$ estimates parameter estimates were positively associated with subsequent preference modification in probe phase, across the different experiments. Dots represent individual stimuli (small vertical jitter was added to better visualize high density areas). Trend lines and surrounding color margins represent estimated preference modification effect and 95% CI, respectively (mixed model logistic regression). Horizontal dashed line represents 50% chance level.

Fig. S5.

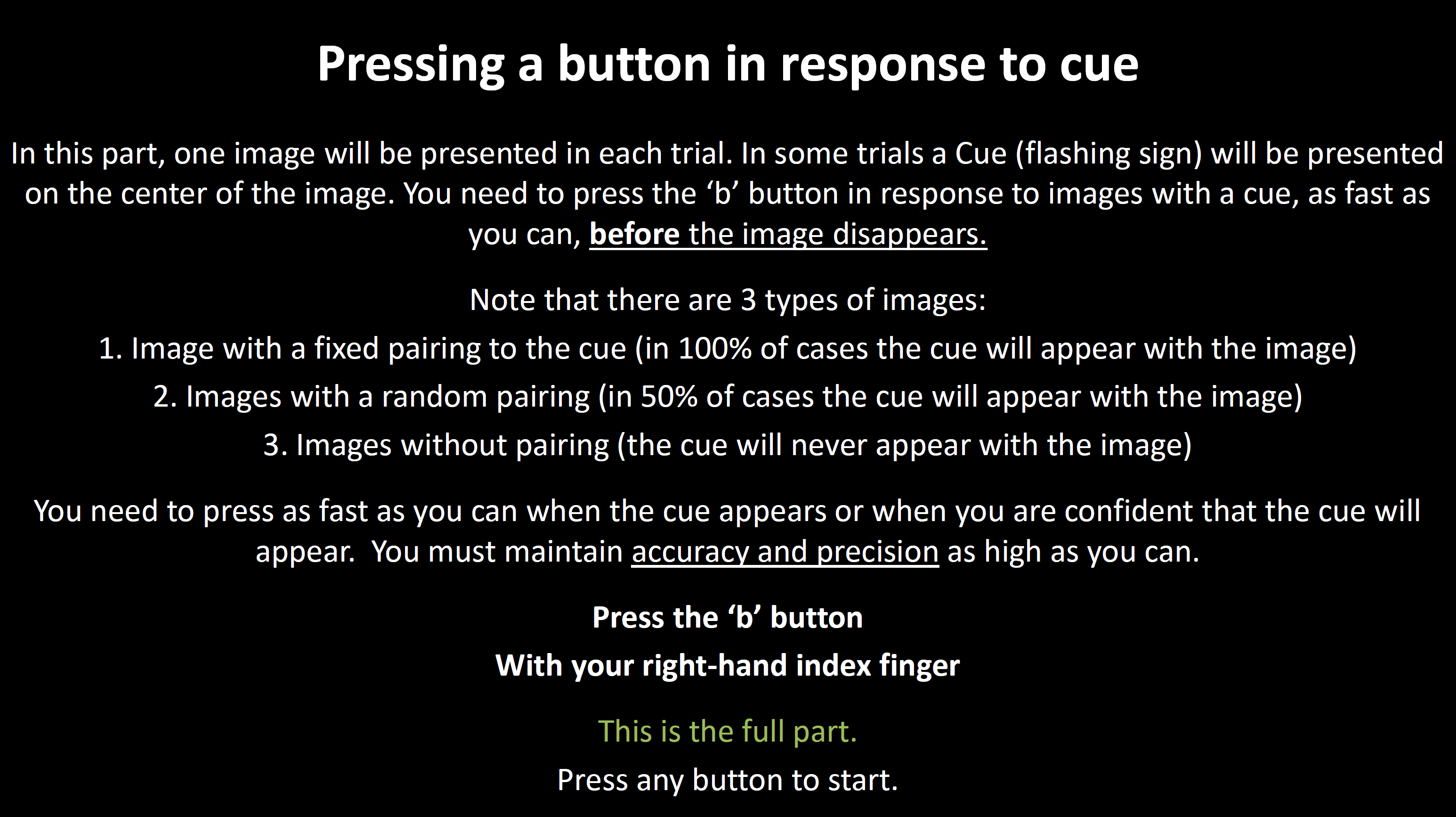

A translated version of the cue-approach training phase instruction presented on the screen before the CAT task in two experiments included in Study 2. Participants were clearly instructed that they may respond before the cue onset, when they anticipate that a cue will appear in association with the image.

Fig. S6.

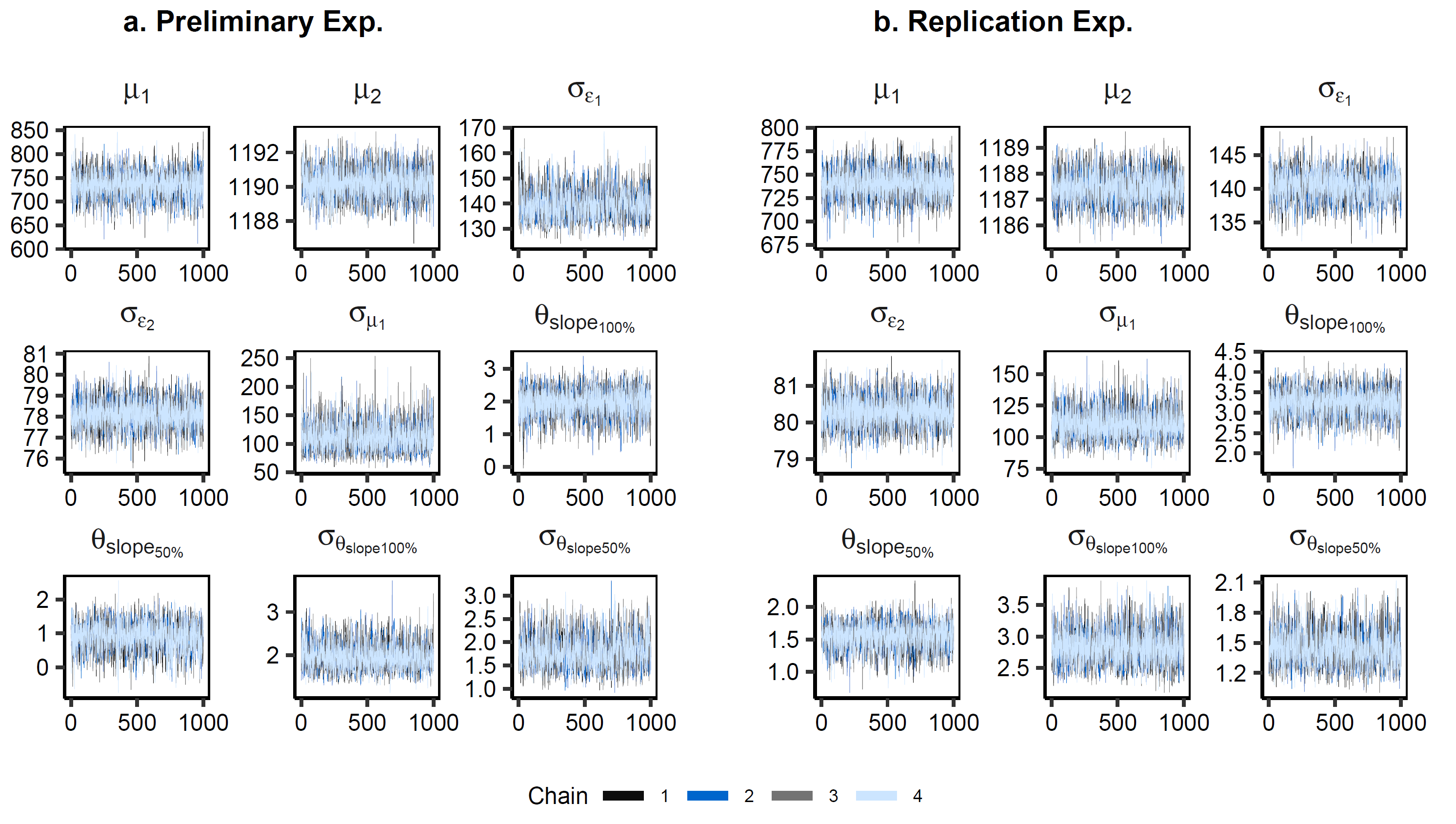

Trace plots of the Stan models in Study 2: (a) Preliminary experiment and (b) Replication experiment. Nine hyper-parameters defined the RT model: two means (μ­_1_, μ­_2_) and *SD* ($\sigma{}_{\varepsilon_{1}}$,$\sigma{}_{\varepsilon_{2}}$) for the normal distribution of anticipatory and cue-dependent RTs, respectively. A mean ($\theta_{slope_{100\%}}, \theta_{slope_{50\%}}$) and *SD* ($\sigma_{\theta_{slope100\%}}, \sigma_{\theta_{slope50\%}}$)for the distribution from which $\theta_{slope_{i}}$ parameters would be drawn for each i^th^ participant in the 100% and 50% contingency. A well-converged parameter is characterized with all four chains (in different colors) reaching stable solution around the same estimate for each parameter (‘hairy caterpillar’ pattern).

Fig. S7.

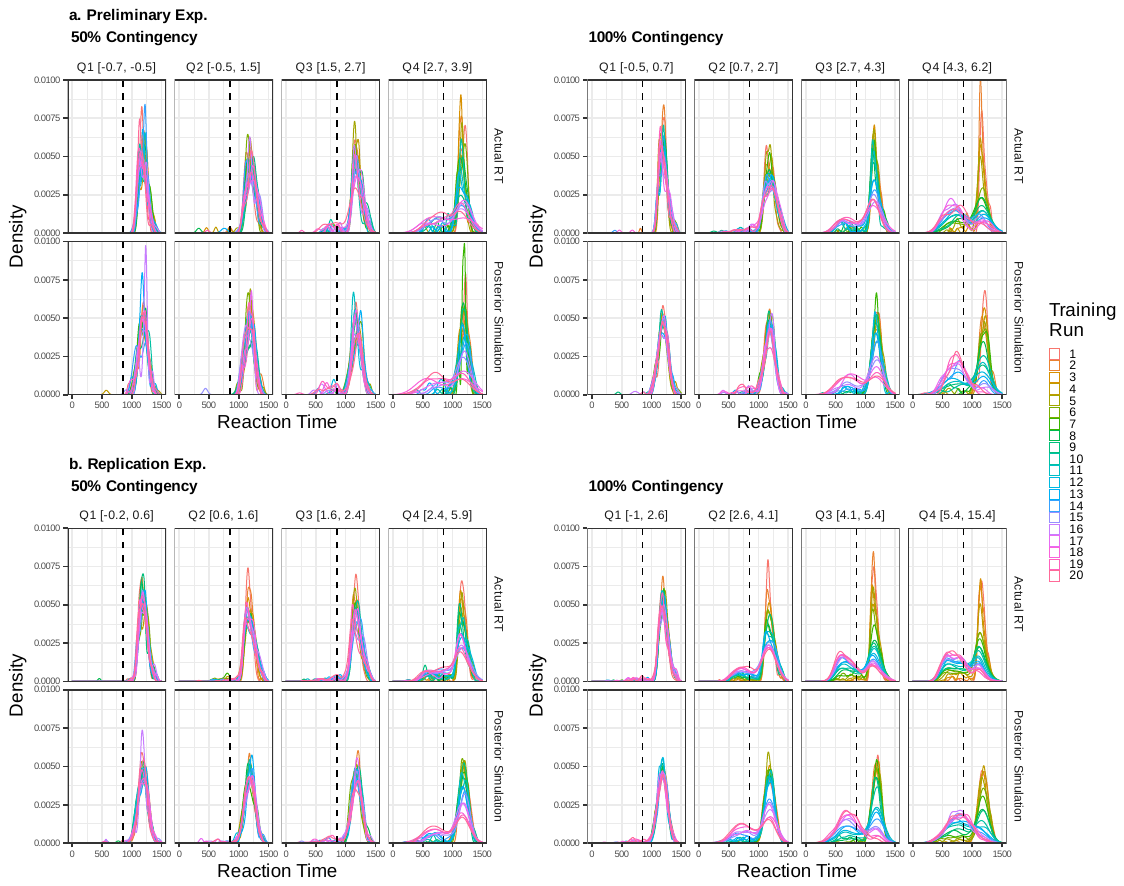

Actual RT distributions versus simulated posterior distributions, by $\theta_{slope_{i}}$ quantile group, in Study 2 (a) preliminary experiment and (b) replication experiment. Participants were categorized into four equal quantile groups, according to their ${\theta_{slope}}_{i}$ parameter estimates (denoted here as Q1-Q4; columns). Participants with higher ${\theta_{slope}}_{i}$ were characterized with faster transition to anticipatory responses (top row). Posterior simulated RT distributions using mixture of Gaussians (bottom row) captured relatively well this transition pattern. Vertical dashed line represents cue-onset.

Fig. S8.

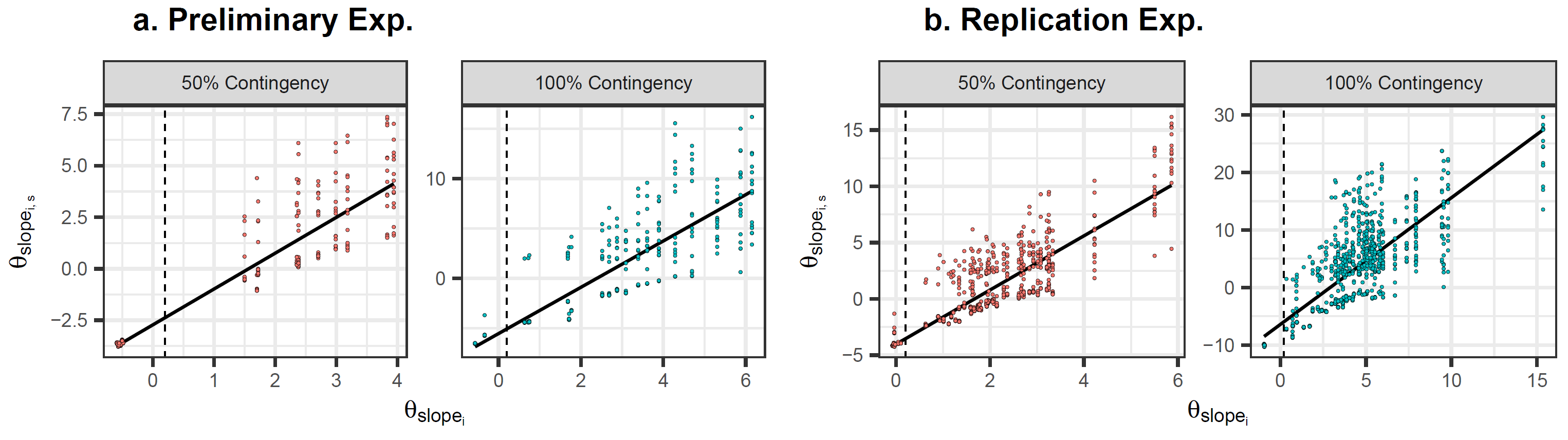

Correlation of $\theta_{slope_{i}}$ parameter fitted individually for each participant and $\theta_{slope_{i,s}}$ fitted individually for each Go stimulus within each participant, in Study 2 preliminary experiment (a) and replication experiment (b). Each dot represents a stimulus within participant. Each participant was fitted with one $\theta_{slope_{i}}$, and 16 $\theta_{slope_{i,s}}$ parameter estimates, per contingency condition. Very low $\theta_{slope_{i}}$ parameter estimates were also characterized with low variability in $\theta_{slope_{i,s}}$ estimates. Participants with small $\theta_{slope_{i}}$ estimate in either condition, were excluded from further logistic regression analysis (vertical dashed line represents a threshold of $\theta_{slope_{i}}$= 0.2).

Fig. S9.

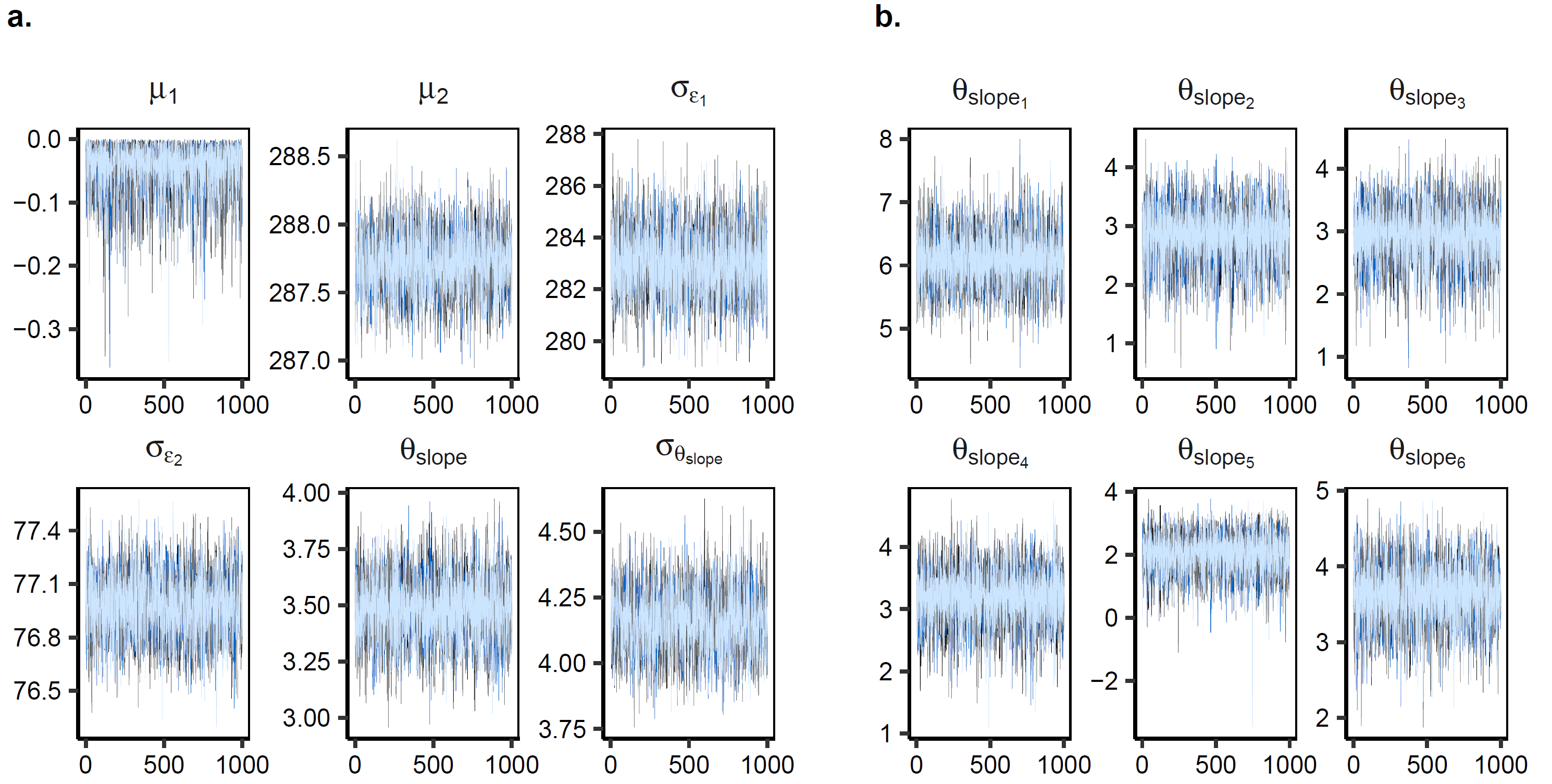

Trace plots of the Stan model of the meta-analysis, with upper limit of μ­_1_ < 0. (a) Six hyper-parameters defined the shape of the RT data: two means ($\mu_{1},\mu_{2}$) and standard deviations($\sigma_{\varepsilon_{1}}, \sigma_{\varepsilon_{2}}$) for the normal distribution of anticipatory and cue-dependent responses, and a mean and *SD* ($\theta_{slope}$, $\sigma_{\theta_{slope}}$) for the distribution from which individualized $\theta_{slope_{i}}$ parameters would be drawn for each participant. The MCMC values of $\mu_{1}$ converged around the upper limit restriction. (b) Trace plot of the first six participants’ $\theta_{slope_{i}}$ parameters. A well-converged parameter is characterized with all four chains (in different colors) reaching stable solution around the same estimate for each parameter (‘hairy caterpillar’ pattern).

Fig. S10.

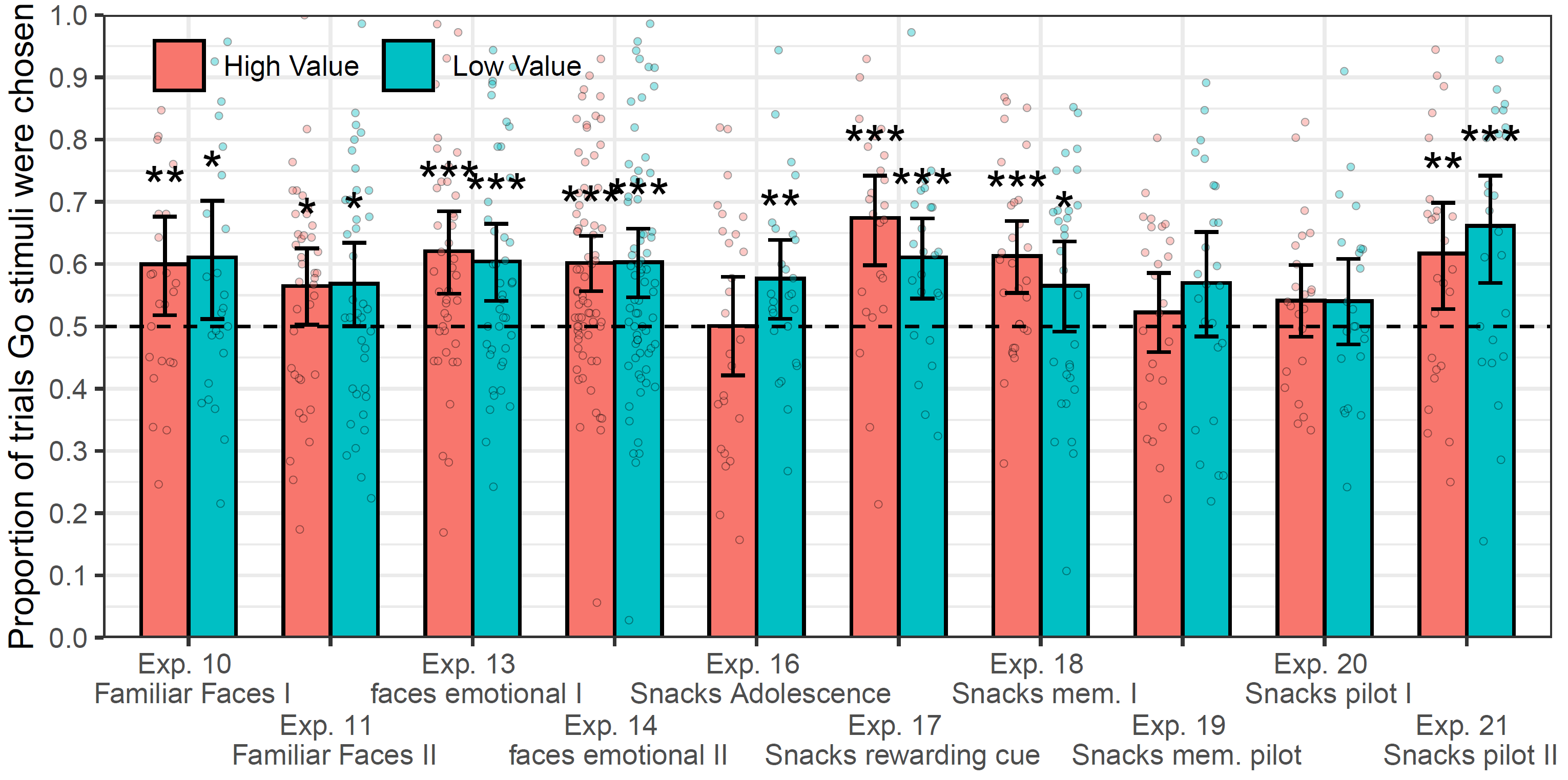

Probe results of unpublished CAT experiments. Proportion of trial participants chose Go stimuli over NoGo stimuli, by experiment and value category. Go stimuli were pitted against NoGo stimuli of similar initial value (both either high- or low-value). In Experiments 13 and 14 (emotional faces), the value category indicates the effect of facial expression (high value = positive affect, low value = neutral affect). Dots represents individual participants. Bars, error bars and asterisks represent the results of a logistic regression analysis, summarizing the mean estimated effect, 95% confidence interval and statistical significance, respectively. *** - *p* < 0.001, ** = *p* < 0.01, * - *p* < 0.05; one-sided mixed-model logistic regression. Models’ results were transformed from log-odds to probabilities unites (odds = exp(log-odds); prob. = odds/(1 + odds)).

Fig. S11.

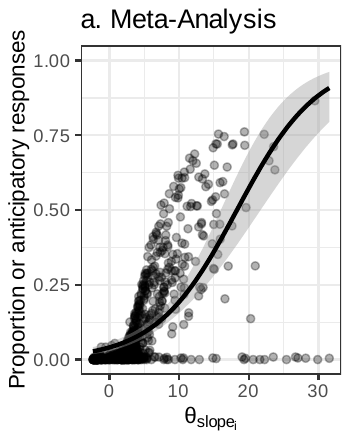

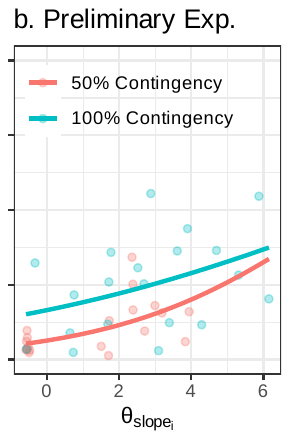

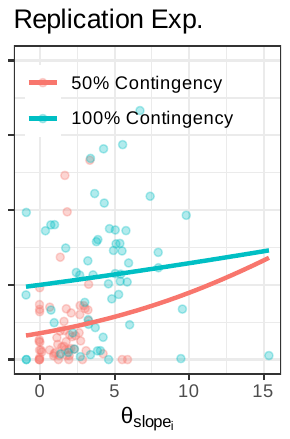

Association of $\theta_{slope_{i}}$ computational parameter and a simple alternative marker of learning, using the proportion of anticipatory responses. (a) association in study 1 meta-analysis (anticipatory response threshold = 144.52ms). (b) association in study 2 preliminary and replication experiments (anticipatory response threshold = 210.2ms). Points represents individual participants, line represents a logistic regression model trendline predicting the proportion of anticipatory responses using $\theta_{slope_{i}}$.

Fig. S12.

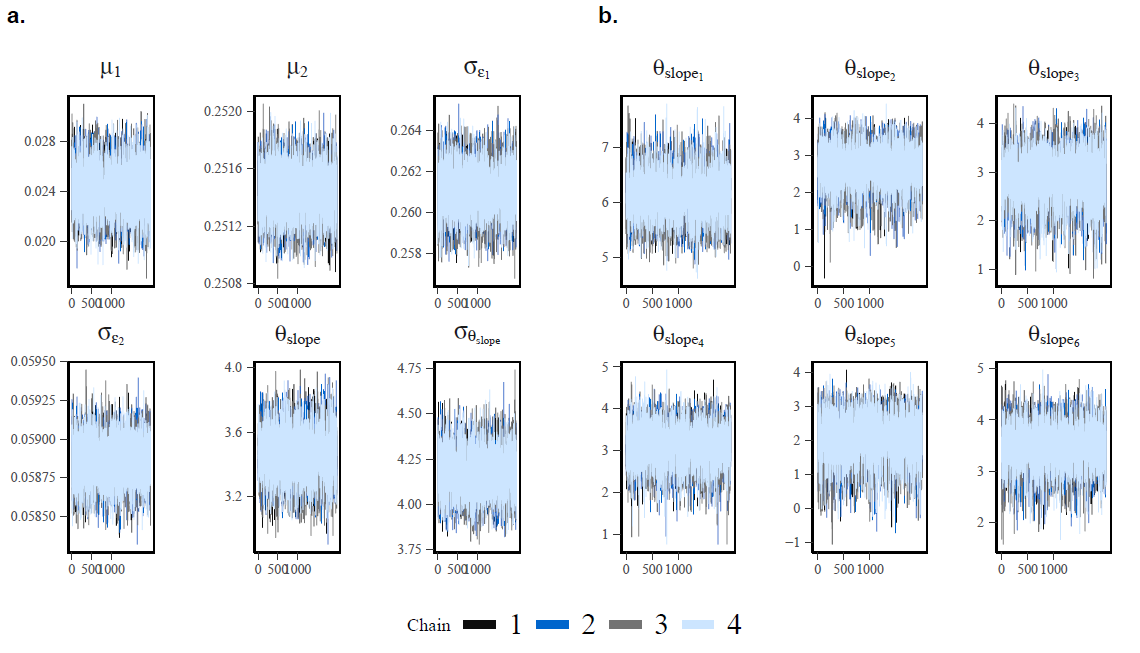
Trace plots of the Stan log-normal model of the meta-analysis. (a) Six hyper-parameters defined the shape of the RT data: two means ($\mu{}_{1}, \mu_{2}$) and standard deviations ($\sigma_{\varepsilon_{1}},\sigma_{\varepsilon_{2}}$) for the normal distribution of anticipatory and cue-dependent responses, and a mean and *SD* ($\theta_{slope}$, $\sigma_{\theta_{slope}}$) for the distribution from which individualized $\theta_{slope_{i}}$ parameters would be drawn for each participant. (b) Trace plot of the first six participants’ $\theta_{slope_{i}}$ parameters. A well-converged parameter is characterized with all four chains (in different colors) reaching stable solution around the same estimate for each parameter (‘hairy caterpillar’ pattern).

Fig. S13.

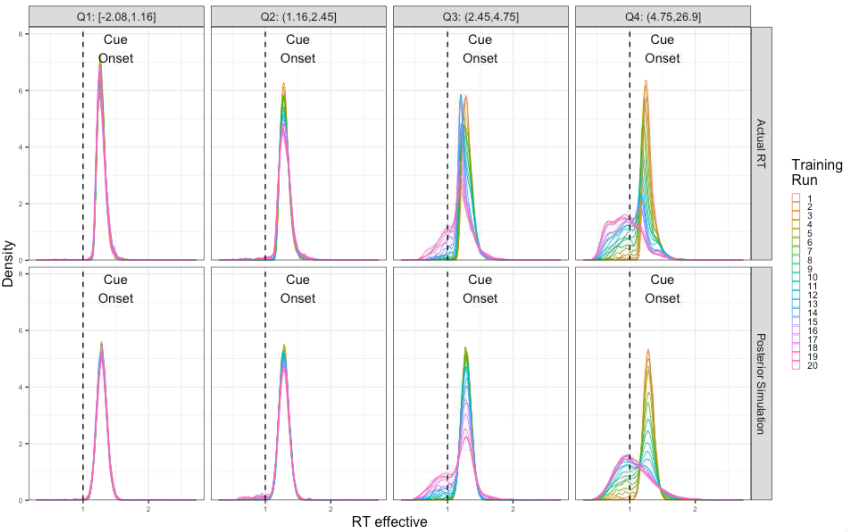

Actual RT distributions versus simulated posterior distributions, by $\theta_{slope_{i}}$ quantile group for Study 1 meta-analysis. Participants were categorized into four equal quantile groups, according to their parameter estimates (denoted here as Q1-Q4; columns). Participants with higher parameter estimates were characterized with faster transition to anticipatory responses (top row). Posterior simulated RT distributions using mixture of log-normal (bottom row) recreated relatively well this transition pattern. Vertical dashed line represents cue-onset.

Fig. S14.

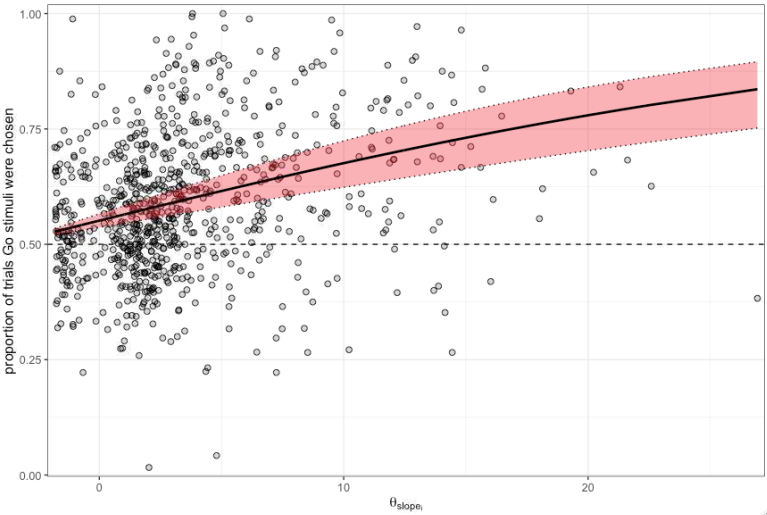

Meta-analysis results - computational marker and preference modification effect. Participants who transitioned faster to anticipatory response during cue-approach training (larger $\theta_{slope_{i}}$ estimates) also demonstrated stronger preference modification effect (proportion of trials Go stimuli were chosen). Trend line and surrounding red margins represent estimated preference modification effect and 95% CI, respectively (mixed model logistic regression). Dots represent individual participants.

Fig. S15.

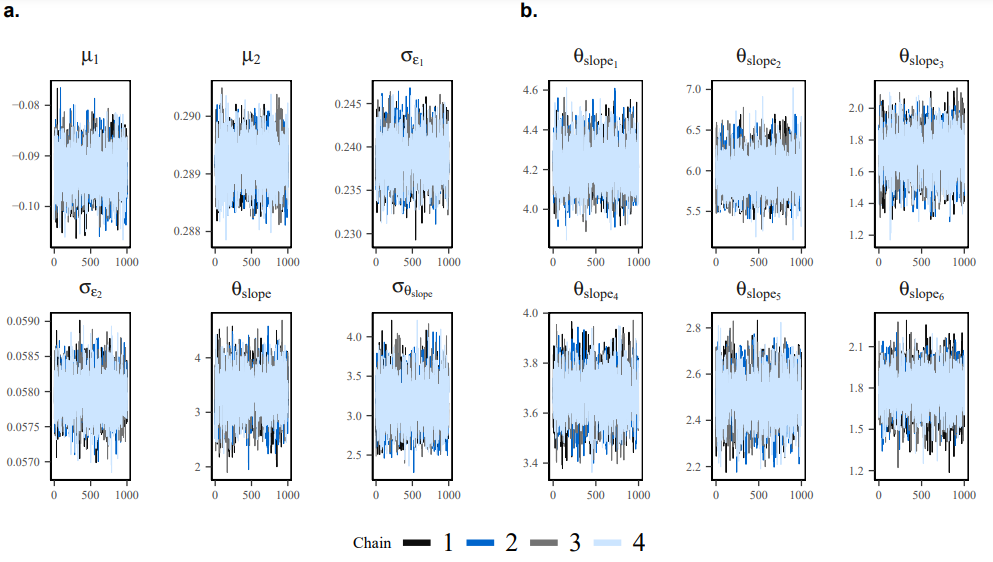

Trace plots of the Stan log-normal model of the designated experiments. (a) Six hyper-parameters defined the shape of the RT data: two means ($\mu{}_{1}, \mu_{2}$) and standard deviations ($\sigma_{\varepsilon_{1}},\sigma_{\varepsilon_{2}}$) for the normal distribution of anticipatory and cue-dependent responses, and a mean and *SD* ($\theta_{slope}$, $\sigma_{\theta_{slope}}$) for the distribution from which individualized $\theta_{slope_{i}}$ parameters would be drawn for each participant. (b) Trace plot of the first six participants’ $\theta_{slope_{i}}$ parameters. A well-converged parameter is characterized with all four chains (in different colors) reaching stable solution around the same estimate for each parameter (‘hairy caterpillar’ pattern).

Fig. S16.

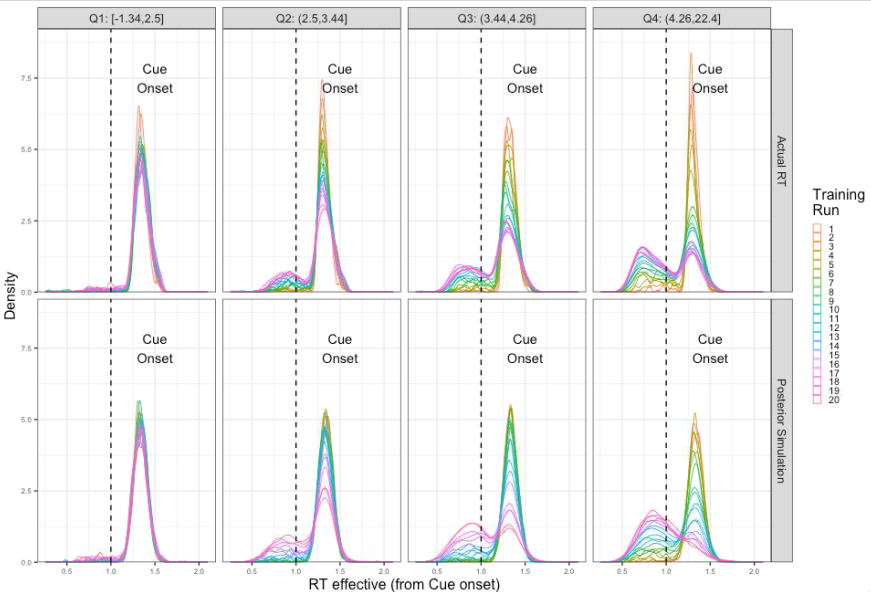

Actual RT distributions versus simulated posterior distributions, by $\theta_{slope_{i}}$ quantile group for Study 2. Participants were categorized into four equal quantile groups, according to their parameter estimates (denoted here as Q1-Q4; columns). Participants with higher parameter estimates were characterized with faster transition to anticipatory responses (top row). Posterior simulated RT distributions using mixture of log-normal (bottom row) recreated relatively well this transition pattern. Vertical dashed line represents cue-onset.

Fig. S17.

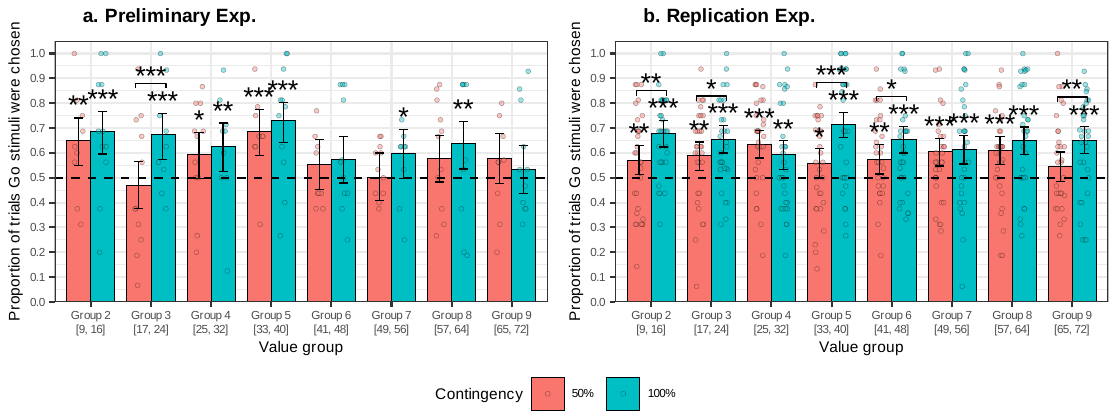

Study 2 probe results, by contingency and value-group. Proportion of trials participants chose Go stimuli over NoGo stimuli of similar initial value, in the preliminary experiment (a) and replication experiment (b). Dots represent individual participants, error-bars represents 95% CI based on a mixed model logistic regression. In square brackets are the value index of the stimuli in the group (lower index indicates higher value). No monotonic trend was observed for value group main effect or interaction with contingency conditions. Statistical significance is denoted with asterisks (* *p* < 0.05, ** *p* < 0.01, *** *p* < 0.001; one-sided mixed model logistic regression). Dashed line represents 50% chance level.

Fig. S18.

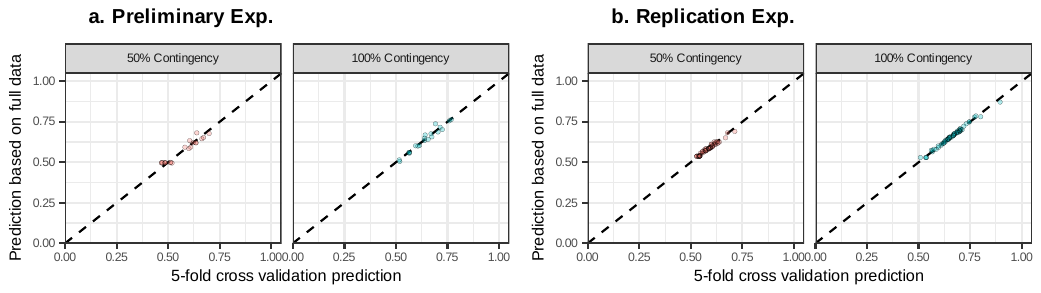

Study 2 five-fold cross validation results. The predicted proportion of trials participants chose Go stimuli over NoGo stimuli based on the full model using the entire dataset (y-axis), plotted against prediction made by CV model (x-axis) which used the data of 80% of the participants to predict the proportion of the held back 20% participant. Results presented for the preliminary experiment (a) and replication experiment (b). Dots represent individual participants; dashed line indicates the identity line. The CV model resulted in nearly identical prediction as the full model.

**Code S1.**

```
// Stan model - Meta analysis (29 experiments). Effective RT: 2 Gaussians mixture model
data {
	// Input: data shape
	int N_subjects; // N = 828 valid participants
	int N_trials; // maximal number of trials
	int N_trials_valid[N_subjects]; // actual number of trials of each participant
	// Input: Dependent (RT-effective) and independent (Run) data
	real RT[N_trials, N_subjects]; // dependent variable: effective RT (RT - Cue)
	real Run[N_trials, N_subjects]; // training run [0,1] indicating (1st, 20th run)
	real theta_b0; // b0 = -3.1, mixture proportion defined at 1st run (set to 0.1%)
}
parameters {
	// Group-level parameters of the two Gaussians
	real <upper=150> mu_fix_1; // early Gaussian Mean (upper limit at cue-onset)
	real <lower=200> mu_fix_2; // late Gaussian Mean (lower limit at cue-onset)
	real <lower=0> sigma_e[2]; // SD of the two Gaussians
	// Group-level (fixed-effect) parameters: theta-slope individualized learning parameter
	real theta_slope_fix; // Mean of individualized theta-slope parameter
	real <lower=0> theta_slope_sd; // SD of individualized theta-slope parameter
	// Participant-level (random-effect) parameter: individualized learning parameter
Real theta_slope[N_subjects]; // parameter to determine mixture proportion
}
model {
// Priors
mu_fix_1 ~ normal(-500,500);
mu_fix_2 ~ normal(500,500);
sigma_e ~ normal(0,1000);
theta_slope_fix ~ normal(0,1);
theta_slope_sd ~ normal(0,0.7);
// Likelihood
	for (i in 1:N_subjects){ // go over participants
	real theta; // mixture proportion
	// participant-level [random-effect] from group-level [fix-effect] parameters
	theta_slope[i] ~ normal(theta_slope_fix,theta_slope_sd);
	for (t in 1:N_trials_valid[i]){ // go over participant’s trials
/* define a linear function of training run with individualized theta-slope use the
inverse of the Normal distribution CDF to go from [-inf,inf] to [0,1] range */
	theta = Phi_approx(theta_b0 + theta_slope[i] * Run[t,i]); // Phi = pnorm function
	// maximize likelihood of parameters' fit to RT data
	target += log_mix(theta, // mixture proportion
normal_lpdf(RT[t,i] | mu_fix_1, sigma_e[1]), // anticipatory RT
normal_lpdf(RT[t,i] | mu_fix_2, sigma_e[2])); // cue-dependent RT
}
}
}
```

Stan model used in Study 1 meta-analysis.

**Code S2.**

```
// Stan model - Study 2. RT: mixture model, with stimulus-level learning parameters
data {
// Input: data shape
int N_subjects; // Number of participants
	int N_trials; // maximal number of Go trials
	int N_stim; // number of unique Go stimuli-participants combinations
	int N_trials_valid[N_subjects]; // actual number of trials of each participant
	// Input: Dependent (RT-effective) and independent (Run and contingency) data
	real RT[N_trials, N_subjects]; // dependent variable: effective RT (RT - Cue)
 	real Run[N_trials, N_subjects]; // training run [0,1] indicating [1st, 20th]
	int Stim[N_trials, N_subjects]; // stimulus indicator
real theta_b0; // b0 = -3.1, mixture proportion defined at 1st run (set to 0.1%)
}
parameters {
// Group-level parameters of the two Gaussians
real <upper=100> mu1_fix; // early Gaussian Mean (upper limit: 100ms after cue)
real <lower=200> mu2_fix; // late Gaussian Mean (lower limit: 200ms after cue)
	real <lower=0> sigma_e[2]; // SD of the two Gaussians
	// Group-level (fixed-effect) parameters: theta-slope individualized learning parameter
	real theta_slope_fix; // individualized theta-slope parameter Mean
	real <lower=0> theta_stim_sd; // SD of stimulus-level theta-slope parameter
	// Stimulus-level (random-effect) parameters: individualized learning parameters
	real theta_slope_stim [N_stim]; // stimuli individualized parameter
}
model {
	//priors
	mu1_fix ~ normal(500,500);
	mu2_fix ~ normal(1000,500);
	sigma_e ~ normal(0,1000);
	theta_slope_fix ~ normal(0,0.7);
	theta_stim_sd ~ cauchy(0,1);
	//likelihood
	// Stimulus-level [random-effect] from group-level [fix-effect] parameters
	theta_slope_stim ~ normal(theta_slope_fix, theta_stim_sd);
	for (i in 1:N_subjects){ // go over participants
	for (t in 1:N_trials_valid[i]){ // go over participant’s trials
	// identify stimulus index
int stim_i;
stim_i = Stim[t,i];
	/* define a linear function of training run with individualized theta-slope use the
	inverse of the Normal distribution CDF to go from [-inf,inf] to [0,1] range */
	// maximize likelihood of parameters' fit to RT data
	target += log_mix(Phi_approx(
theta_b0 + theta_slope_stim[stim_i] * Run[t,i]), // mixture proportion
	normal_lpdf(RT[t,i] | mu1_fix, sigma_e[1]), // anticipatory RT
	normal_lpdf(RT[t,i] | mu2_fix, sigma_e[2])); // cue-dependent RT
	}
	}
}
```

Stan model used in Study 1 with stimulus-level learning parameter.

**Code S3.**

```
// Stan model - Study 2. RT: 2 Gaussians mixture model, with 2 contingency conditions
data {
// Input: data shape
	int N_subjects; // n=20 or n = 59
	int N_trials; // maximal number of Go trials
	int N_trials_valid[N_subjects]; // actual number of trials of each participant
	// Input: Dependent (RT-effective) and independent (Run and contingency) data
	real RT[N_trials, N_subjects]; // dependent variable: RT
 	real Run[N_trials, N_subjects]; // training run [0,1] indicating [1st, 20th]
	real Contingency[N_trials, N_subjects]; // 50% (0) or 100% (1) contingency condition
	real theta_b0; // b0 = -3.1, mixture proportion defined at 1st run (set to 0.1%)
	real Cue; // fixed at 850ms
}
parameters {
// Group-level parameters of the two Gaussians
real <lower=0, upper=Cue-100> mu1_fix; // early Gaussian Mean (limit: 100ms before cue)
real <lower=Cue> mu2_fix; // late Gaussian Mean (lower limit at cue-onset)
	real <lower=0> sigma_e[2]; // SD of the two Gaussians
	real <lower=0> mu1_sd[1]; // SD of individualized early Gaussian mean parameter
	// Group-level (fixed-effect) parameters: theta-slope individualized learning parameter
	real theta_fix[2]; // individualized theta-slope parameter Means (2 conditions)
	real<lower=0> theta_sd[2]; // SDs of individualized theta-slope parameter
	// Participant-level (random-effect) parameters: individualized learning and early mean
	real <upper=Cue+100> mu1_rand[N_subjects]; // individualized mean for early RT
	real theta_slope_100[N_subjects]; // individualized learning-parameter - 100% contingency
	real theta_slope_50[N_subjects]; // individualized learning-parameter - 50% contingency
}
model {
	//priors
	mu1_fix ~ normal(500,500);
	mu2_fix ~ normal(1000,500);
	sigma_e ~ normal(0,1000);
	mu1_sd ~ normal(0,150);
	theta_fix ~ normal(0,0.7);
	theta_sd ~ normal(0.7,0.7);
	//likelihood
	for (i in 1:N_subjects){ // go over participants
	// participant-level [random-effect] from group-level [fix-effect] parameters
	mu1_rand[i] ~ normal(mu1_fix,mu1_sd); // individualized early RT mean
	theta_slope_100[i] ~ normal(theta_fix[1],theta_sd[1]); // 100% contingency
	theta_slope_50[i] ~ normal(theta_fix[2],theta_sd[2]); // 50% contingency
	for (t in 1:N_trials_valid[i]){ // go over participant’s trials
	/* define a linear function of training run with individualized theta-slope use the
	inverse of the Normal distribution CDF to go from [-inf,inf] to [0,1] range */
	// maximize likelihood of parameters' fit to RT data
	target += log_mix(Phi_approx(
theta_b0 + (Contingency[t,i]*theta_slope_100[i] +
(1-Contingency[t,i])*theta_slope_50[i]) * Run[t,i]), // mixture proportion
	normal_lpdf(RT[t,i] | mu1_rand[i] , sigma_e[1]), // anticipatory RT
	normal_lpdf(RT[t,i] | mu2_fix, sigma_e[2])); // cue-dependent RT
	}
	}
}
```

Stan model used in Study 2 main-analysis.

**Code S4.**

```
// Stan model - Study 2. RT: mixture model, with stimulus-level learning parameters (each contingency separately)
data {
// Input: data shape
	int N_subjects; // n=20 or n = 59
	int N_trials; // maximal number of Go trials
	int N_stim; // number of Go stimuli (=16)
	int N_trials_valid[N_subjects]; // actual number of trials of each participant
	// Input: Dependent (RT-effective) and independent (Run and contingency) data
	real RT[N_trials, N_subjects]; // dependent variable: RT
 	real Run[N_trials, N_subjects]; // training run [0,1] indicating [1st, 20th]
	int Stim[N_trials, N_subjects]; // stimulus indicator
real theta_b0; // b0 = -3.1, mixture proportion defined at 1st run (set to 0.1%)
	real Cue; // fixed at 850ms
}
parameters {
// Group-level parameters of the two Gaussians
real <lower=0, upper=Cue+100> mu1_fix; // early Gaussian Mean (limit: 100ms after cue)
real <lower=Cue> mu2_fix; // late Gaussian Mean (lower limit at cue-onset)
	real <lower=0> sigma_e[2]; // SD of the two Gaussians
	// Group-level (fixed-effect) parameters: theta-slope individualized learning parameter
	real theta_fix; // individualized theta-slope parameter Means
	real<lower=0> theta_sub_sd; // SD of participant-level theta-slope parameter
	real<lower=0> theta_stim_sd; // SD of stimulus-level theta-slope parameter
	// Participant-level (random-effect) parameters: individualized learning parameters
	real theta_slope_sub [N_subjects]; // participants individualized parameter
	real theta_slope_stim [N_subjects, N_stim]; // stimuli individualized parameter
}
model {
	//priors
	mu1_fix ~ normal(500,500);
	mu2_fix ~ normal(1000,500);
	sigma_e ~ normal(0,1000);
	theta_fix ~ normal(0,0.7);
	theta_sub_sd ~ cauchy(0,1);
	theta_stim_sd ~ cauchy(0,1);
	//likelihood
	// participant-level [random-effect] from group-level [fix-effect] parameters
	theta_slope_sub ~ normal(theta_fix,theta_sub_sd);
	for (i in 1:N_subjects){ // go over participants
	// stimulus-level parameter model variability around participant-level parameter
	theta_slope_stim[i,] ~ normal(theta_slope_sub[i], theta_stim_sd);
	for (t in 1:N_trials_valid[i]){ // go over participant’s trials
	// identify stimulus index
int stim_i;
stim_i = Stim[t,i];
	/* define a linear function of training run with individualized theta-slope use the
	inverse of the Normal distribution CDF to go from [-inf,inf] to [0,1] range */
	// maximize likelihood of parameters' fit to RT data
	target += log_mix(Phi_approx(
theta_b0 + theta_slope_stim[i, stim_i] * Run[t,i]), // mixture proportion
	normal_lpdf(RT[t,i] | mu1_fix, sigma_e[1]), // anticipatory RT
	normal_lpdf(RT[t,i] | mu2_fix, sigma_e[2])); // cue-dependent RT
	}
	}
}
```

Stan model used in Study 2 with stimulus-level learning parameter. The model was evaluated separately for each condition (100% contingency and 50% contingency).

**Code S5.**

```
// Stan model - Meta analysis (29 experiments). Effective RT: 2 log-normal mixture model
data {
	// Input: data shape
	int N_subjects; // N = 828 valid participants
	int N_trials; // maximal number of trials
	int N_trials_valid[N_subjects]; // actual number of trials of each participant
	// Input: Dependent (RT-effective) and independent (Run) data
	real RT[N_trials, N_subjects]; // dependent variable: effective RT (RT - Cue)
	real Run[N_trials, N_subjects]; // training run [0,1] indicating (1st, 20th run)
	real theta_b0; // b0 = -3.1, mixture proportion defined at 1st run (set to 0.1%)
}
parameters {
	// Group-level parameters of the two Gaussians
	real <upper=0.14> mu_fix_1; // early Gaussian Mean (upper limit at cue-onset)
	real <lower=0.18> mu_fix_2; // late Gaussian Mean (lower limit at cue-onset)
	real <lower=0> sigma_e[2]; // SD of the two Gaussians
	// Group-level (fixed-effect) parameters: theta-slope individualized learning parameter
	real theta_fix; // Mean of individualized theta-slope parameter
	real <lower=0> theta_sd; // SD of individualized theta-slope parameter
	// Participant-level (random-effect) parameter: individualized learning parameter
Real theta_b1[N_subjects]; // parameter to determine mixture proportion
}
model {
// Priors
mu_fix_1 ~ normal(-0.693,1);
mu_fix_2 ~ normal(0.4,0.4);
sigma_e ~ normal(0,1);
theta_fix ~ normal(0,1);
theta_sd ~ normal(0,0.7);
// Likelihood
	for (s in 1:N_subjects){ // go over participants
	real theta; // mixture proportion
	// participant-level [random-effect] from group-level [fix-effect] parameters
	theta_b1[i] ~ normal(theta_fix,theta_sd);
	for (t in 1:N_trials_valid[i]){ // go over participant’s trials
/* define a linear function of training run with individualized theta-slope use the
inverse of the Normal distribution CDF to go from [-inf,inf] to [0,1] range */
	theta = Phi_approx(theta_b0 + theta_b1[i] * Run[t,s]); // Phi = pnorm function
	// maximize likelihood of parameters' fit to RT data
	target += log_mix(theta, // mixture proportion
lognormal_lpdf(RT[t,s] | mu_fix_1, sigma_e[1]), // anticipatory RT
lognormal_lpdf(RT[t,s] | mu_fix_2, sigma_e[2])); // cue-dependent RT
}
}
}
```

Alternative Stan model used in Study 1 using log-normal distributions.

**Table S1.** Results - Unpublished experiments

| Experiment | *n* | Training  runs | Prop. Go stimuli  were chosen | *p* ^1^ | OR (95% CI) |
| --- | --- | --- | --- | --- | --- |
| Exp. 10 | 25 | 20 | High value 58.4% | 0.009 | 1.50 [1.07, 2.09] |
| Familiar Faces I |  |  | Low value 58.4% | 0.014 | 1.57 [1.05, 2.35] |
| Exp. 11 | 39 | 20 | High value 55.6% | 0.021 | 1.30 [1.01, 1.67] |
| Familiar Faces II |  |  | Low value 55.5% | 0.025 | 1.32 [1.00, 1.73] |
| Exp. 13 | 42 | 20 | Pos. affect 60.0% | 3.1e-04 | 1.64 [1.23, 2.17] |
| Emotional Faces I |  |  | Neut. affect 58.7% | 7.0e-04 | 1.53 [1.18, 1.98] |
| Exp. 14 | 70 | 20 | Pos. affect 59.1% | 6.3e-06 | 1.51 [1.26, 1.82] |
| Emotional Faces II |  |  | Neut. affect 58.4% | 1.9e-04 | 1.52 [1.21, 1.91] |
| Exp. 16 | 26 | 12 | High value 50.1% | 0.494 | 1.00 [0.73, 1.38] |
| Snacks Adolesce. |  |  | Low value 56.8% | 0.010 | 1.36 [1.05, 1.77] |
| Exp. 17 | 24 | 20 | High value 65.7% | 7.8e-06 | 2.07 [1.49, 2.88] |
| Snacks rewarding cue |  |  | Low value 60.1% | 5.9e-04 | 1.57 [1.20, 2.06] |
| Exp. 20 | 23 | 12 | High value 53.8% | 0.080 | 1.18 [0.94, 1.49] |
| Snacks pilot I |  |  | Low value 53.5% | 0.126 | 1.18 [0.89, 1.55] |
| Exp. 21 | 25 | 20 | High value 59.7% | 0.005 | 1.61 [1.12, 2.32] |
| Snacks pilot II |  |  | Low value 63.8% | 3.7e-04 | 1.95 [1.32, 2.88] |

*Notes:*

^1^ One-sided logistic regression.
